# Multiple losses of ecdysone receptor genes in nematodes: an alternative evolutionary scenario of molting regulation

**DOI:** 10.1101/2025.01.27.634690

**Authors:** Shumpei Yamakawa, Lisa-Marie Barf, Andreas Hejnol

## Abstract

Molting is a hallmark feature of ecdysozoans, including arthropods, tardigrades, and nematodes. Ecdysone hormones play a crucial role in regulating the molting process of different ecdysozoan taxa. Interestingly, despite this highly conserved function of ecdysone, the model nematode *Caenorhabditis elegans* has lost the ecdysone receptor (ECR) genes and their molting appears to be ecdysone-independent. The loss of *ecr* has only been reported in *Caenorhabditis* within ecdysozoans, and the evolutionary background behind this loss has remained enigmatic. Here, we show that loss of *ecr* is not exceptional in *Caenorhabditis*, but has occurred at least three times in Rhabditina and Tylenchina nematodes. Our genome-wide analysis of 160 nematode species revealed multiple losses of *ecr* and its typical heterodimer partner *usp* during nematode evolution. Furthermore, using transcriptomic, pharmacological, and *in silico/ in vivo* protein interaction analyses, we identified two factors that potentially underlie and buffer the loss of ECR gene/function: (1) molting regulation by an alternative nuclear receptor HR3 (NHR-23) and (2) a lineage-specific expansion of nuclear receptors in the *ecr-*deficient taxa. Taken together, this study shows how key regulators of ecdysozoan molting can be altered during evolution. We propose a novel scenario for the evolution of molting regulation in nematodes.

## Introduction

Ecdysozoa, which includes arthropods, nematodes, priapulids, and others, comprises more than 80% of all animal species (Aguinaldo, et al. 1997; Dunn, et al. 2008; Hejnol, et al. 2009). A defining feature of ecdysozoans is their ability to grow and change appearance by periodically shedding a cuticular exoskeleton (Nielsen 2012; Giribet and Edgecombe 2017; Schumann, et al. 2018). As the name Ecdysozoa derives from this unique growth strategy, molting or ecdysis (Aguinaldo, et al. 1997), the acquisition of a cuticular exoskeleton and molting process is a key innovation in ecdysozoan evolution. Despite its importance for understanding of ecdysozoan evolution, the evolutionary background of molting remains enigmatic.

Molting regulation has been well characterized in arthropods such as insects and decapods with ecdysone hormone as a master regulator (Fig. 1B) (Lachaise, et al. 1993; Thummel 1996; Yamanaka, et al. 2013; Niwa and Niwa 2014; Hyde, et al. 2019). In arthropods, ecdysone hormone is synthesized from cholesterol by a group of genes called the Halloween genes (e.g., *sad* and *phm*) in specific endocrine organs such as the prothoracic glands (Niwa and Niwa 2014). The secreted ecdysone is converted to 20-hydroxyecdysone (20E), which binds to the ecdysone receptor (ECR) in the target tissues (e.g., epidermal cells) (Yamanaka, et al. 2013). The ecdysone receptor (ECR) forms a heterodimer with another nuclear receptor USP to activate downstream cascade genes (e.g., HR3 and FTZ-F1, Fig. 1B) (Yamanaka, et al. 2013). Recently, we found that tardigrade molting is likely regulated by ecdysteroid hormone (Yamakawa and Hejnol 2024), suggesting that ecdysone-dependent molting is ancestral in panarthropods.

**Figure 1.**
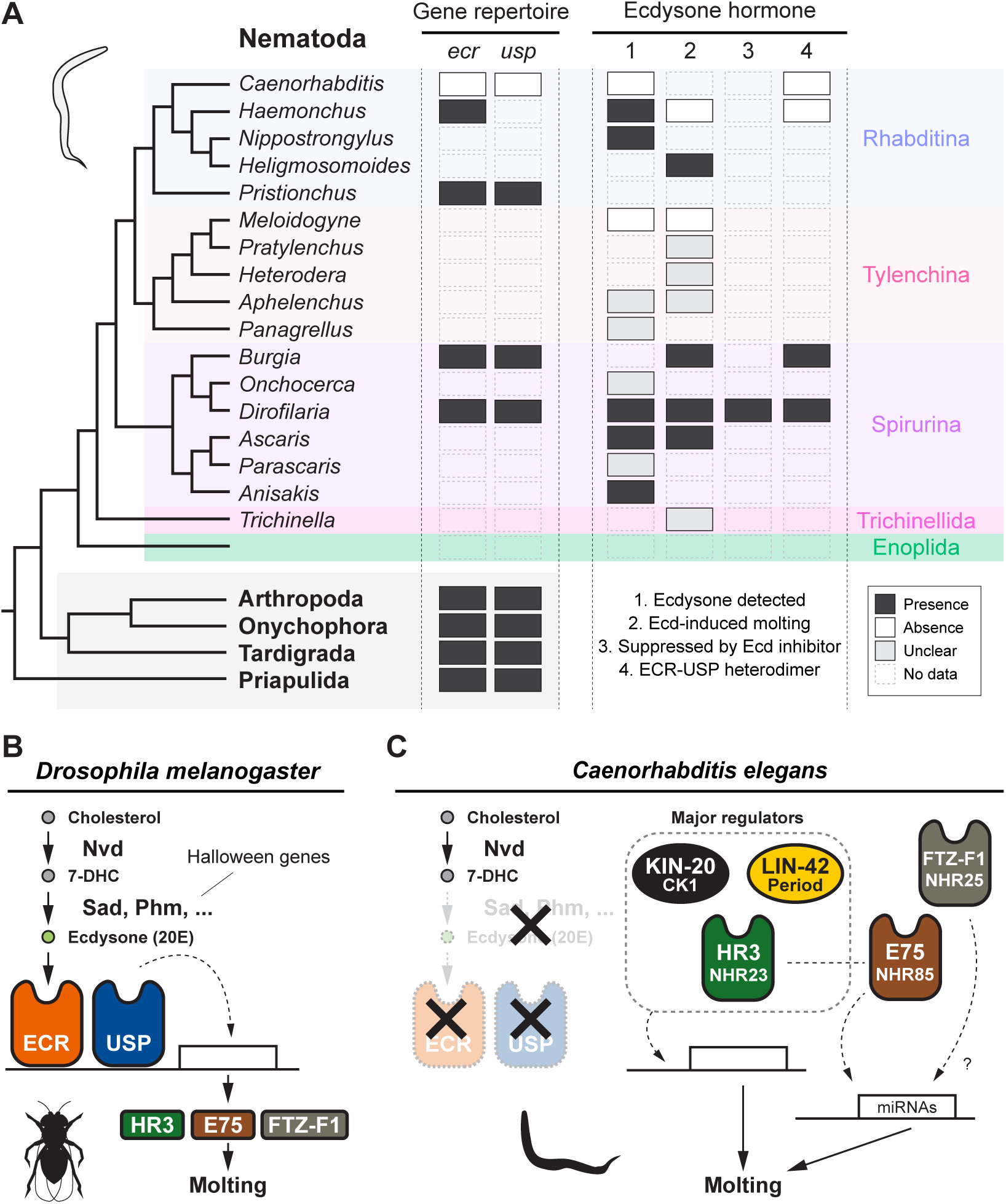
An overview of previous work on the role of ecdysone hormone and molting regulation in nematodes and arthropods. **A**: (left two columns) Presence and absence of the ecdysone receptor (*ecr*) and its heterodimer partner gene *usp* in ecdysozoan taxa. (right four column) A summary of the findings on ecdysone hormonal control in nematodes. 1: detection of ecdysteroid hormone in the internal body, 2: molting induction by exogenous ecdysteroids (=Ecd), 3: molting suppression by exogenous ecdysteroid antagonists, and 4: the functionality of ECR-USP heterodimer. As shown in the legends of the panels, black, white, gray, and empty boxes indicate the presence, absence, unclear, and no information, respectively. **B** and **C**: Schematic diagrams of the molting regulatory mechanisms in *Drosophila melanogaster* and *Caenorhabditis elegans*, respectively. 20E (20-hydroxyecdysone) is synthesized from cholesterol by the Nvd and Halloween genes, such as Sad (Shadow) and Phm (Phantom). The ECR-USP heterodimer binds to 20E, activating a cascade of downstream genes, including HR3, E75, and FTZ-F1. *C. elegans* is known to lack genes that synthesize ecdysone and its receptors, which play an important regulatory role in insect molting. See the main text for details on each regulatory mechanism. The reference and the details of **A** are shown in Table S1.

In nematodes, the developmental role of ecdysone hormone has been studied primarily in parasitic species (Fig. 1A and Table S1). For example, ecdysone and/or 20E have been detected in Rhabditina and Spirurina parasitic species, including *Haemonchus contortus*, *Anisakis simplex*, *Dirofilaria immitis*, and *Ascaris suum* (Fig. 1A, Table S1) (Fleming 1985; Cleator, et al. 1987; Evershed, et al. 1987; Fleming 1993). In *Ascaris suum*, 20E exhibits pulsatile fluctuations synchronized with the molting cycle, resembling patterns reported in arthropods (Fleming 1985). Pharmacological functional studies have also been conducted to test whether ecdysone hormone regulates molting process (Fig. 1A and Table S1). Exogenous ecdysone or 20E treatment is reported to induce or prompt the molting of *Heligmosomoides polygyrus* (molting of L4 [4th larval stage] to adult), *Ascaris suum* (L4-adult), *Dirofilaria immitis* (L3-L4), and *Burgia malayi* (L3-L4) (Fig. 1A and Table S1) (Dennis 1976; Fleming 1985; Warbrick, et al. 1993; Mhashilkar, et al. 2016). Notably, Warbrick et al., 1993 showed that the ecdysone agonist RH5849 and antagonists Azadirachtin have a potential to induce and suppress L3-L4 molting of *D. immitis*, respectively. Similar to the pleiotropic functions of ecdysone in arthropods (Kozlova and Thummel 2000; Subramoniam 2000), effects of pharmacological treatment were also found in reproduction-related processes (ex., oocyte development in *D. immitis* and microfilarial release in *Brugia pahangi*) (Delves, et al. 1986; Barker, et al. 1991). Finally, genetic components of ecdysone signaling, including ecdysone receptor and its heterodimer functionality, have also been identified in Rhabditina and Spirurina nematodes (Fig. 1A and Table S1) (Tzertzinis, et al. 2010). The above findings support the involvement of ecdysone hormone in the molting regulation of some nematode lineages such as Spirurina.

Interestingly, however, inconsistent results with ecdysone-dependent molting have been also reported in some nematode species (ex., no effects of ecdysone treatment in the molting of *Haemonchus contortus* (L3–L4) and *Meloidogyne javanica* (L2–L3) (Fig. 1A and Table S1) (Soriano, et al. 2004; Graham, et al. 2010). Remarkably, the model nematode *Caenorhabditis elegans* is known to lack any genes coding the ecdysone hormone receptor (ECR) (Schumann, et al. 2018; Taubenheim, et al. 2021). As *ecr* has been identified in different nematode taxa (Shea, et al. 2004; Graham, et al. 2010; Parihar, et al. 2010; Shea, et al. 2010; Tzertzinis, et al. 2010), loss of *ecr* in *C. elegans* is so far the only demonstrated case among ecdysozoans (Fig. 1A). *C. elegans* is also known to lack any genes coding *usp* (Schumann, et al. 2018; Taubenheim, et al. 2021), and molting of *C. elegans* must be therefore independent of ecdysone (Chitwood and Feldlaufer 1990; Frand, et al. 2005; Hill, et al. 2013). Rather, the molting of *C. elegans* is regulated by a mechanism similar to the biological/circadian clock (Fig. 1C) (Hendriks, et al. 2014; Meeuse, et al. 2020; Tsiairis and Großhans 2021; Meeuse, et al. 2023). For example, evolutionary conserved components of circadian clock, HR3/NHR-23 (ROR), LIN-42 (Period), and KIN-20 (CK1ε/δ) dominantly regulate the molting periods in *C. elegans* (Kostrouchova, et al. 1998, 2001; Banerjee, et al. 2005; Frand, et al. 2005; Kouns, et al. 2011; Monsalve, et al. 2011; Rhodehouse, et al. 2018; Patel, et al. 2022; Johnson, et al. 2023; Kinney, et al. 2023; Hiroki and Yoshitane 2024; Lamberti, et al. 2024; Spangler, et al. 2024). Another circadian clock component, E75/NHR-85 (Rev-Erb), is also known to be involved in the molting clock of *C. elegans*; the temporal formation of HR3/NHR-23 and E75/NHR-85 heterodimer generates a transcriptional pulse of the microRNA *lin-4* (Kinney, et al. 2023). Feedback loops involving HR3/NHR-23 and other genes/microRNAs (e.g., *let-7*) have also been identified in *C. elegans* (Kouns, et al. 2011; Patel, et al. 2022; Johnson, et al. 2023). NHR-25, an ortholog of FTZ-F1 which regulates insect molting, is a regulator of *C. elegans* molting, while its function is likely minor as through miRNA expression control (Fig. 1B, C) (Gissendanner, et al. 2004; Hayes, et al. 2006; Hada, et al. 2010; Meeuse, et al. 2023). Other transcription factors such as GRH-1 have been also identified as a molting regulator in *C. elegans* (Fig 1C) (Hauser, et al. 2022; Meeuse, et al. 2023).

Molting regulation of *C. elegans* is likely a derived state within nematodes. The loss of *ecr in C. elegans* is also inconsistent with the idea that molting regulation is conservative and stereotypic in nematodes, in which most species consistently molt four times regardless of the divergence in the life cycles (Malakhov 1986). Thus, it is generally considered that the loss of *ecr* in nematodes is a specific and rare event unique to *C. elegans* (Taubenheim, et al. 2021), and elucidating the evolutionary background of *ecr* loss has been a long-standing challenge. The loss of *ecr* in *C. elegans* is a critical subject for understanding the evolution of ecdysozoan molting since it is implied that the key regulator for molting can be altered during evolutionary processes. In summary, the case of *C. elegans* raises doubts on the evolutionary conserved role of ecdysone.

This study investigated the evolutionary trajectory leading to the loss of *ecr* and *usp* in nematodes, aiming to elucidate how key regulators of ecdysozoan molting can be altered. We examined the presence of *ecr* and *usp* in a total of 160 nematode species and elucidated the evolution of nematode molting regulation using genomic, transcriptomic, and protein prediction analyses.

## Results

### 1. Multiple losses of *ecr* and *usp* genes in Rhabditina and Tylenchina nematodes

Although *ecr* genes have been identified in some nematodes outside *Caenorhabditis* (Fig. 1A) (Shea, et al. 2004; Graham, et al. 2010; Parihar, et al. 2010; Shea, et al. 2010; Tzertzinis, et al. 2010), the specificity of *ecr* loss has been insufficiently explored within a phylogenetic framework (Fig. 1A). Thus, using publicly available genomic and transcriptome data, we searched for *ecr* across 160 nematode species including major nematode taxa (Fig. 2A, B, 159 genomic and 1 transcriptomic data, see also the methods). The genes of *ecr* were searched using BLAST and phylogenetic analysis (see the methods).

**Figure 2.**
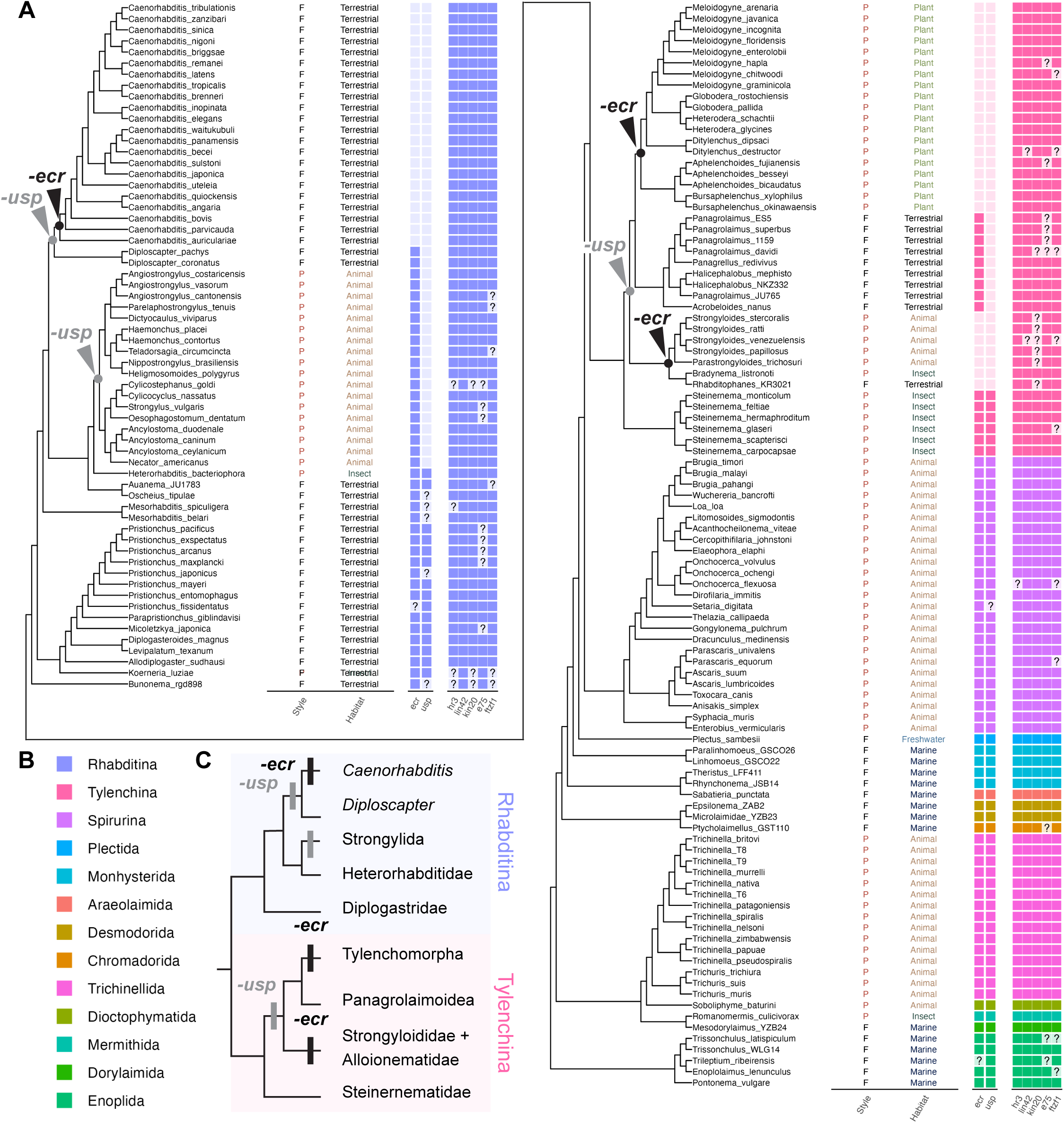
Gene repertory of the target genes (*ecr, usp, hr3, kin-20 lin-42, e75, and ftz-f1)* in 160 nematode species. **A**: Lifestyle (P: parasite and F: free-living), habitat, and gene repertories (*ecr, usp, hr3, lin-42, kin-20, e75, and ftz-f1*) are indicated with the phylogeny of the target nematode species (color represents the taxa, question marks represent missing information). The nodes of *ecr* and *usp* loss events are highlighted in the tree. **B**: Legends for color and taxa. **C**: A simpler tree clarifying the loss of ECR and USP in the Rhabditina and Tylenchina groups

#### Gene mining of ecr and usp in 160 nematode species

Our search identified *ecr* genes from all major nematode taxa, including the sister group of all remaining nematodes, Enoplida (Fig. 2A). In Rhabditina, *ecr* was not found in any *Caenorhabditis* species examined (Fig. 2A), and the loss of *ecr* is specific to this lineage. Unexpectedly, *ecr* is also absent in 26 of 41 Tylenchina species examined (Fig. 2A). No *ecr* was found in the species of Tylenchomorpha, Strongyloididae, and Alloionematidae, and the latest phylogenic analyses consistently support the polyphyly of the three groups (Fig. 2C, see methods) (Smythe, et al. 2019; Ahmed, et al. 2022; Qing, et al. 2024). Thus, the loss of *ecr* is indicated as occurring twice in Tylenchina (Fig. 2A, C). We also mapped the presence and absence of *ecr* onto the phylogenetic trees reconstructed by Ahmed et al., 2022, and the occurrence of multiple losses of *ecr* is still supported (Fig. S1). Together, the loss of *ecr* occurred at least three times during nematode evolution. Note that the *ecr* genes, previously identified in *Strongyloides* species (Gonzalez Akimori, et al. 2021), are incorrectly annotated (Fig. S2).

*C. elegans* is also known to lack the gene encoding USP (Taubenheim, et al. 2021), a typical heterodimer partner of ECR. Our gene mining revealed that *usp* is also absent in several lineages of Rhabditina and Tylenchina (Fig. 2A). Especially, the loss of *usp* is detected not only in the species lacking *ecr* but more broadly observed as no *usp* was identified in some Rhabditina species having *ecr* (ex., Strongylida: Fig. 2A, C). The loss of *usp* also precedes the loss of *ecr* in Tylenchina as only Steinernematidae have both *ecr* and *usp* (Figs 2A, C, and S1). Thus, it is suggested that the ECR-USP functionality has been lost several times in nematodes. Interestingly, *ecr* was lost only in the lineages which lost *usp* among the species examined, and *ecr* and *usp* losses are not related to specific lifestyle/habitat (Fig. 2A).

#### Additional analyses of ecr and usp loss

We confirmed that the absence of *ecr* and *usp* is not due to technical errors or low-quality genome assembly (Figs S3–S5, genome size, number of scaffolds, assembly N50, and BUSCO score). As these figures show, no association with specific metrics was observed.

We also surveyed ecdysone receptors (*ecr* and *usp*) and biogenetic genes (Halloween genes, Fig. 1B) in ecdysozoans and nematodes, respectively, motivated by our unexpected finding of multiple *ecr* losses. We obtained consistent results with previous reports. For example, *ecr* was newly identified in all ecdysozoan species examined (Fig. S6): nematomorphs (Gordioida and Nectonematoida), tardigrades (Heterotardigrada), kinorhynchs, and priapulids. No Halloween genes (*spo*, *phm*, *dib*, *sad*, or *shade*) were identified in any of the nematode species examined.

### 2. Putative modification of ecdysone hormone’s roles in molting regulation with the loss of *ecr* and *usp* genes

We subsequently examined the evolutionary background of multiple losses of *ecr* and *usp*. We first speculated that the involvement of ecdysone in molting regulation is specifically lost in parallel with *ecr* and *usp* losses. To test this hypothesis, we performed the following pharmacological functional and temporal gene expression analysis.

#### ECR antagonist suppresses the molting process of Plectida nematodes which possess ecr and usp genes

To investigate the function of ecdysone hormone in nematode molting, we examined the effects of the ECR antagonist, Cucurbitacin B (CucB), treatment on the molting of three nematode species (Fig. 3A): *C. elegans* (Rhabditina lacking *ecr* and *usp*), *Diploscapter pachys* (Rhabditina lacking *usp*), and *Plectus sambesii* (Plectida having *ecr* and *usp*). The phylogeny of the three species reflects the evolutionary changes of *ecr* and *usp* losses (Fig. 3A, B).

**Figure 3.**
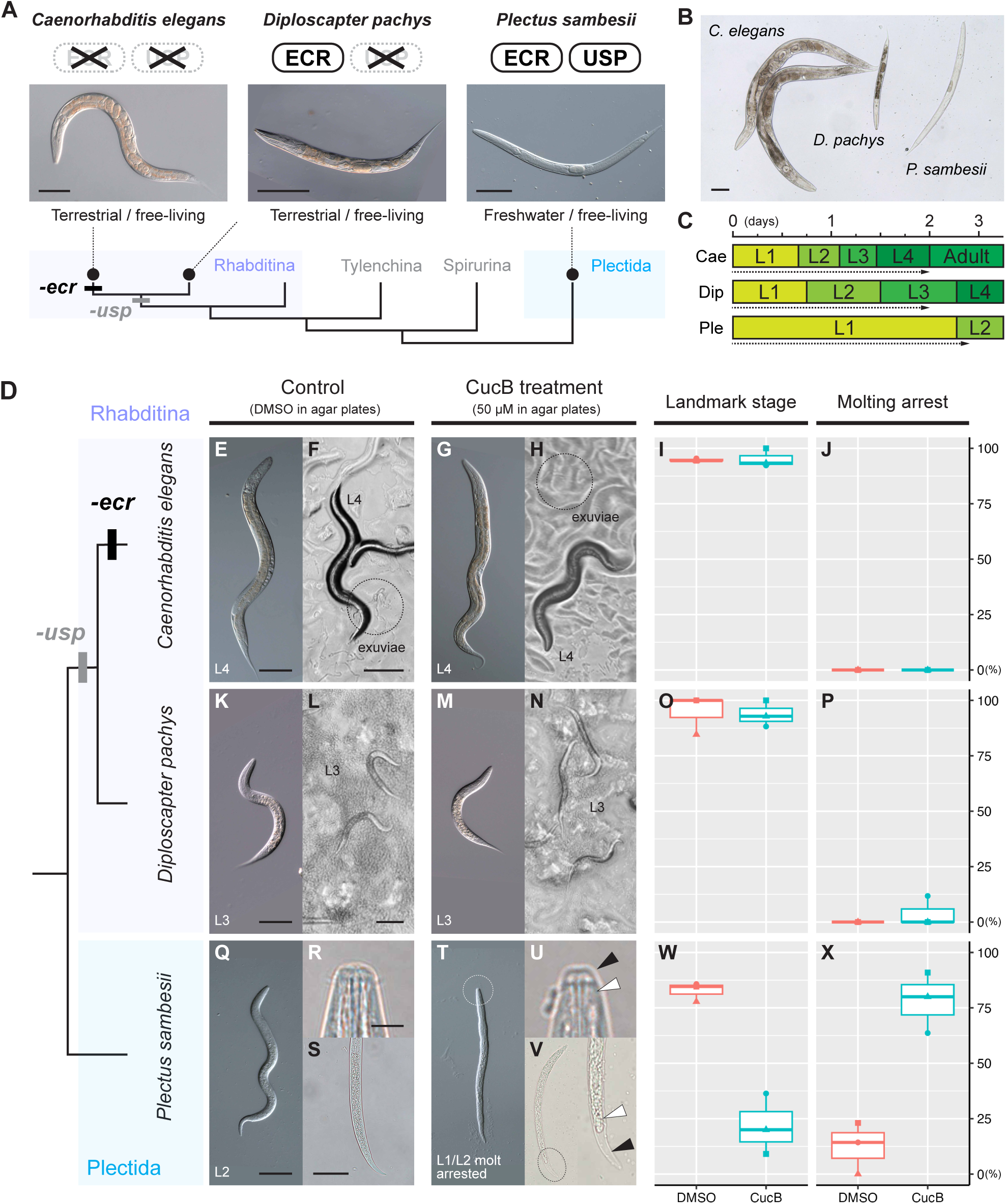
Effects of the ECR antagonist, Cucurbitacin B, treatment on the molting process in *Caenorhabditis elegans*, *Diploscapter pachys,* and *Plectus sambesii*. **A**: The phylogeny of the *ecr*/*usp* repertoire in the three target species. The photos show the adult gravid worms with eggs (scale bars: 100 µm). **B**: A comparison of body size among representative adult individuals of the three species (scale bar: 100 µm). **C**: Growth speed and molting timing after feeding on L1 worms in each species (Cae: *C. elegans*; Dip: *D. pachys*; and Ple: *P. sambesii*). L1–L4 represent the first to fourth larval stages. See the Methods and Fig. S7 for detailed information about larval and adult staging. Dotted arrows indicate the duration of cucurbitacin B (CucB) treatment. **D**–**X:** the results of CucB treatment. **D** shows the phylogeny of the target species. The results for *C. elegans*, *D. pachys*, and *P. sambesii* are shown in **E**–**J**, **K**–**P**, and **Q**–**X**, respectively. Representative worms after DMSO treatment are shown in **E**–**F**, **K**–**L**, and **Q**–**S**; CucB: **G**–**H**, **M**–**N**, and **T**–**V**. **E**, **G**, **K**, **M**, **Q**, and **T**: DIC images of the treated worms; **F**, **H**, **L**, and **N**; the worms on agar plates; **R**, **S**, **U**, and **V**: magnified images of the mouth (**R** and **U**) and tail (**S** and **V**) parts, indicating the process of apolysis during CucB treatment (the black and white arrowheads indicate the old and new cuticles, respectively). Scale bars for **E**/**G**, **F**/**H**, and **L**/**N**: 100 µm; for **K**/**M**, **Q**/**T**, and **S**/**V**: 50 µm; and for **R**/**U**: 5 µm. **I**, **Q**, and **W**: the percentage of DMSO– or CucB-treated worms that developed to the landmark stage. **J**, **P**, and **X:** the percentage of DMSO– or CucB-treated worms that showed molting arrest/defect phenotypes.

In all species examined, larval worms molted four times after hatching to become gravid adults (L1–L4 and adult stages). We collected synchronized L1 larvae and monitored growth speed and molting timing after feeding for each species on optimized agar culture plates at 20 °C (see Methods, Fig. S7A). Gonadal development was primarily used as the key for defining specific larval and adult stages (Fig. S7B, C) (Tahseen, et al. 1991; TAHsEEN, et al. 1992; Hubbard and Greenstein 2000; Schindler and Sherwood 2013). For example, elongated gonadal primordia were observed after the L3 stage, and the vulva developed clearly only after the L4 stage (Fig. S7B, C) (Hubbard and Greenstein 2000). Regarding molting cycle duration, *C. elegans* develops very rapidly, with young adults emerging as early as two days after feeding at 20 °C (Figs. 3C, S7A). Growth in *D. pachy* is slightly slower, requiring approximately three to four days after feeding until the young adults emerge (Figs. 3C, S7A). *P. sambesii* is the slowest developer and molts approximately every one to two days (Figs. 3C, S7A). Under our stable cultivation conditions, larval growth was comparatively synchronous within each species.

We designed the CucB treatment experiments to target specific molting stages, as shown in Figs 3C and S7A. For example, in *C. elegans* and *D. pachys*, CucB was applied for two days, and we assessed whether treated worms progressed to the L4/adult and L3 stages, respectively (Fig. 3C). Feeding started with the treatment of CucB, and DMSO was used as control. In *P. sambesii*, we performed two experiments (Fig. S7A). First, L1 worms (0 days after feeding) were treated with CucB for three days, and their development to the L2 stage was examined at the day three (Fig. 3C). Second, we targeted the progression from L3 to L4 by applying CucB for three days starting at day 4 after feeding (Fig. S7A). Based on preliminary toxicity assays, a concentration of 50 µM CucB was used consistently in all experiments (three replicates).

Results are shown in Figs 3D–X, S8 and Table S2, and the effects on molting were only observed in *P. sambesii*. Around 80% L1 worms developed to L2 worms in the control experiments (DMSO, 24 of 29 worms, Fig. 3Q–S, W), while only around 20% worms molted to L2 in CucB treatment (7 of 32 worms, Fig. 3T–V, W). In the remaining 25 of 32 CucB-treated L1 worms, apolysis process (a first step of molting process) occurred around mouth parts (Fig. 3T–U), but their development/growth was arrested at this stage. Some worms also molted the tail parts but did not complete molting process (Fig. 3V). Such phenotypes are considered as molting defects or growth arrest at molting stage. Although molting arrest was observed in some control worms (7/32, Fig. 3X), the difference between two treatments was statistically significant (*p*=0.0032, Two Sample t-test, Fig. 3X). We also observed similar phenotypic effects of CucB treatment in the experiments targeting L3-L4 molting (growth: 39/45 [DMSO], 8/58 [CucB]; molting arrest: 0/45 [DMSO], 49/58 [CucB]; Fig. S8). These effects of CucB on molting resemble the observations in the examples of insects and tardigrades (Toyofuku, et al. 2021; Yamakawa and Hejnol 2024). Especially, considering that the CucB-treated worms proceed to apolysis stage, CucB is suggested to specifically suppress the ecdysis process.

On the other hand, no effects on growth and molting were observed in *C. elegans* (Fig. 3E–J). Almost all worms developed to L4/adult stages (DMSO: 55/58, CucB: 48/50) and no molting defects were found (Fig. 3I, J). This result is consistent with the ecdysone-independent molting in *C. elegans*. Interestingly, CucB treatment did not affect the growth and molting of *D. pachys* which has *ecr* genes but not *usp* (Fig. 3K–P). Of 49 L1 worms, 47 and 46 worms grew to L3 stage in DMSO and CucB, respectively (Fig. 3O), and molting arrest was found in only two CucB treated worms (Fig. 3P). Since we used the same concentration of CucB among the three species, its effects may be more severe in smaller bodied worms. Nevertheless, molting was unaffected by CucB in *D. pachys*, the smallest of the three examined species, at a concentration sufficient to suppress molting in the larger species, *P. sambesii* (Fig. 3B).

#### Conserved temporal expression of ecr and usp during molting cycle in Trichinellida, Spirurina, and Rhabditina nematodes

We next examined the temporal expression patterns of *ecr* and *usp* in the representative nematodes. Previous studies have shown that molting-related genes, including *ecr*, exhibit oscillatory expression peaking just before ecdysis in various ecdysozoan species (Gao, et al. 2015; Liu, et al. 2022; Yamakawa and Hejnol 2024). Based on this, we investigated if the temporal expression of *ecr* and *usp* shows similar pattern during molting cycle among nematode species.

We examined the temporal expression of *ecr* during the molting cycles of four nematode species (Fig. 4A, B, Rhabditina: *Pristionchus pacificus* [having *ecr* and *usp*], *Ancylostoma ceylanicum* [lacking *usp*], Spirurina: *Brugia malayi*, and Trichinellida: *Trichuris suis*), for which transcriptome data during the molting cycle are publicly available (see methods and Fig. S9). In *T. suis*, *ecr* expression showed a sharp peak pattern before L3-L4 and L4-adult molting (Fig. 4B). Similar (but more moderate) peaked expression was observed in L3-L4 molting of *B. malayi* (Fig. 4B). On the other hand, although a pulsed expression of *ecr* was detected in L2-L3 and L3-L4 molting of *P. pacificus*, the expression peaks did not precede the start of molting but peaked at the same time (or later) as ecdysis, the final step of molting (Fig. 4B, see also the method and Fig. S9). Importantly, the expression of *ecr* in *A. ceylanisum*, which lacks *usp*, showed a comparatively minor increase during L3-L4 molting (Fig. 4B).

**Figure 4.**
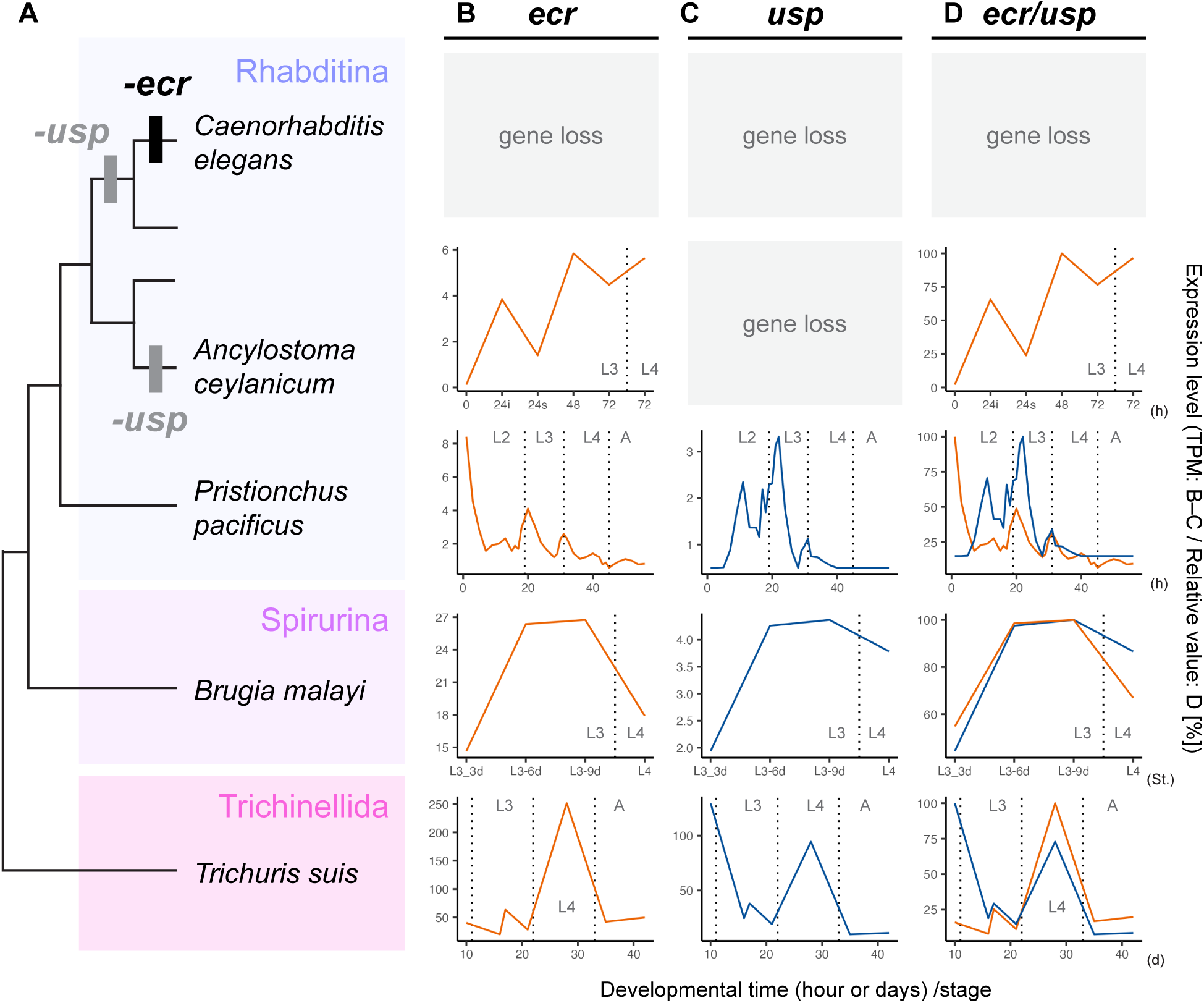
Temporal expression pattern of *ecr* and *usp* during the molting cycle in the representative nematode species. **A**: Phylogeny of the target nematode species and occurrence of *ecr* and *usp* loss events. **B**–**D**: Temporal expression patterns of *ecr*, *usp,* and both genes, respectively. The x-axis indicates developmental time (hours/h, days/d, or stages/St.), and the y-axis shows expression levels (transcripts per million [TPM] in **B** and **C** and relative expression level [%] in **D**). The vertical dotted lines indicate the timing of molting or ecdysis (L1–L4: first to fourth larval stages; A: adult stage).

We also investigated the expression of *usp* genes in three species which possess *usp* (Fig. 4A, C: *P. pacificus*, *T. suis*, and *B. malayi*). Similar to *ecr*, *usp* also shows a peaked expression pattern before molting in *T. suis* and *B. malayi* (Fig. 4B). The temporal expression of *ecr* and *usp* is similar in both species (Fig. 4B–D). *usp* of *P. pacificus* also shows comparatively similar expression pattern with *ecr* (Fig. 4B–D), although the expression pattern is less clearly overlapped than in *T. suis* and *B. malayi* (Fig. 4B–D). Similar expression of *P. pacifucs ecr* and *usp* was previously observed during molting cycles in a semi-quantitative PCR analysis (*Ppa-pnhr2* and *Ppa-pnhr-1*, respectively, 3–48 hours after hatching with a three-hour interval) (Parihar, et al. 2010).

#### Summary of pharmacological and expression analysis

ECR antagonist suppresses the molting process of a Plectida species, and the peaked expression of *ecr* and *usp* before molting is conserved in Trichinellida, Spirurina, and some Rhabditina nematodes (e.g., *P. pacificus*). They all possess both *ecr* and *usp* genes. Pharmacological suppression of ECR has been also reported to inhibit the molting process in the Spirurina species *Dirofilaria immitis* (Warbrick, et al. 1993). However, such molting suppression and peaked expression of *ecr* are not observed in the species lacking *ecr* and/or *usp*. Thus, it is plausible to interpret that involvement of ecdysone/*ecr* in molting process is altered and even lost in parallel with the loss of *ecr* and *usp* in the Plectida+Rhabditida (Rhabditina, Tylenchina, and Spirurina) taxon.

### 3. Screening for conserved molting regulators among nematodes

Although we found the putatively divergent molting function of ECR, the molting process itself appears consistent across nematodes (e.g., most nematodes consistently molt four times (Malakhov 1986)). This suggests the presence of other key regulators which buffer the loss of *ecr* and *usp*. As a first step toward identifying potential candidate genes, this study focused on the *C. elegans* molting regulatory genes (Fig. 1C). Specifically, we examined the expression pattern of the five representative regulators (Fig. 1C): *hr3* (*nhr-23*), *lin-42* (*per*), *kin-20 (ck1/εδ)*, *e75* (*nhr-85*), and *ftz-f1* (*nhr-25*).

First, our survey revealed that all genes are highly conserved across nematode lineages (Fig. 2A). Only *e75* genes may be lost in some *Pristionchus* and *Panagrolaimus* lineages (Fig. 2A). We then examined the expression pattern of the five genes during the molting cycle in six nematode species (Fig. 5, the above four species, *ecr*-lacking Rhabditina *C. elegans*, and *ecr-*lacking Tylenchina *Globodera pallida*). Notably, pulsed expression of *hr3* prior to molting was observed in all six species (Fig. 5B). For example, *hr3* expression shows a peaked pattern prior to molting in *A. ceylanisum* which does not show peaked *ecr* expression, and *G. pallida* which lacks *ecr* (Figs 4B, 5B). A similar expression pattern was observed in *lin-42* of *C. elegans*, *A. ceylanicum*, *P. pacificus*, and *B. malayi* (Fig. 5C). However, this pattern was not observed during the molting process of *G. pallida* and *T. suis* (Fig. 5C). The temporal expression patterns of *kin-20*, *e75* and *ftz-f1* are also not conserved among the six species examined (Fig. 5D–F).

**Figure 5.**
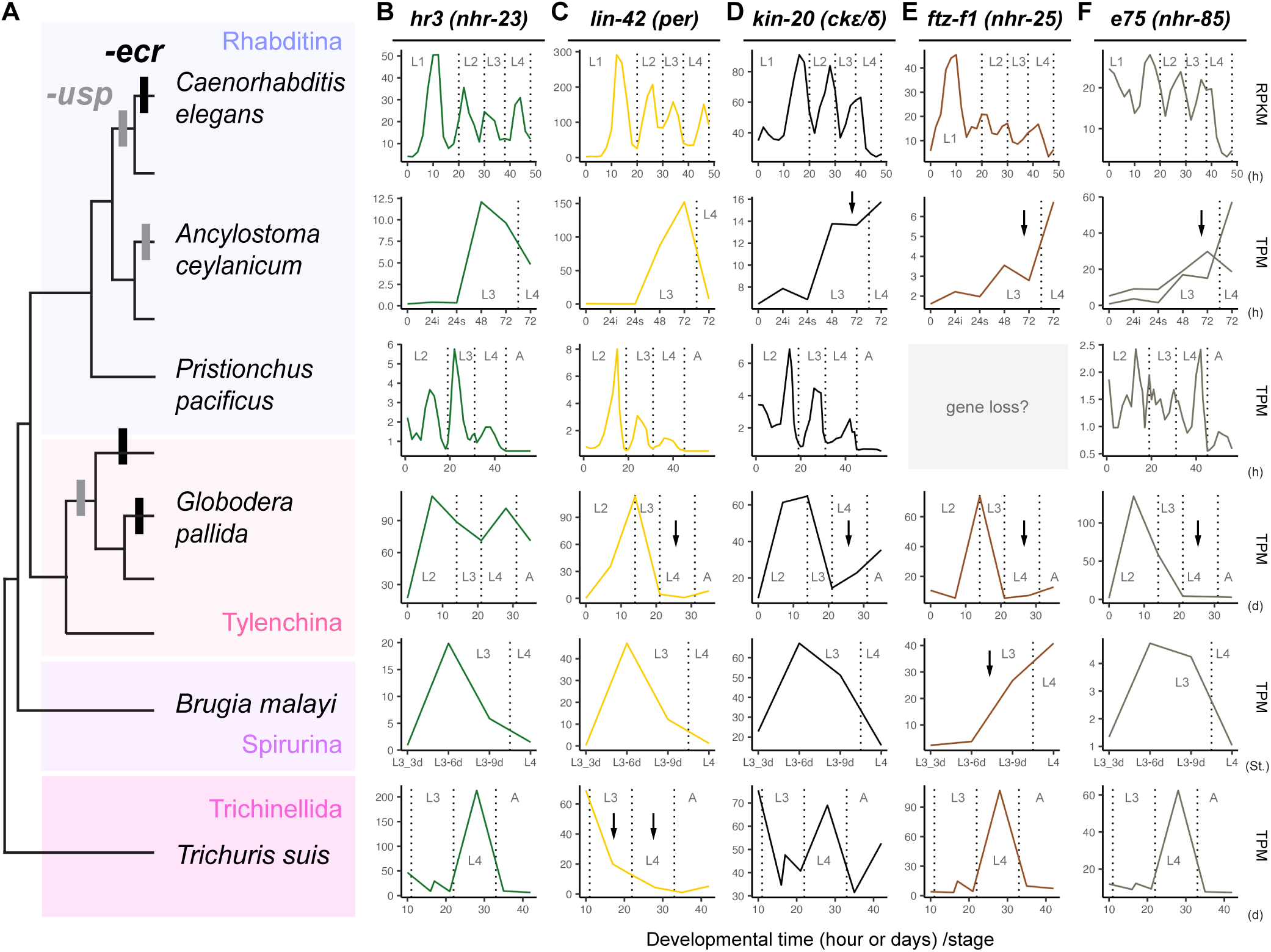
Temporal expression pattern of *hr3*, *lin-42*, *kin-20, e75,* and *ftz-f1* during the molting cycle in representative nematode species. **A**: Phylogeny of the target nematode species and occurrence of *ecr* and *usp* loss events. **B**–**F**: Temporal expression patterns of *hr3/nhr-23*, *lin-42/per, kin-20/ck1εδ*, *e75/nhr-*85, and *ftz-f1/nhr-25* genes, respectively. The x-axis indicates developmental time (hours/h, days/d, or stages/St.), and the y-axis shows expression levels (RPKM or TPM). The vertical dotted lines indicate the timing of molting or ecdysis (L1–L4: first to fourth larval stages; A: adult stage). The arrows indicate that there was no increased or peaked in expression just before molting/ecdysis.

We also performed systematic and unbiased screening of the candidate conserved molting regulators across the above six species. Previously, the genes showing oscillatory expression during the molting cycle were extracted from *C. elegans* (1,592 genes) and *P. pacificus* (2,964 genes) (Kim, et al. 2013; Sun, et al. 2021). For the remaining four species, we performed a fuzzy c-mean clustering analysis on each transcriptome dataset and extracted the cluster(s) showing peaked expression just before molting in each species (Fig. S10; green shades; *A. ceylanicum*: 8,110; *G. pallida*: 3,383; *B. malayi*: 7,114; and *T. suis*: 2,896 genes). Then, we used OrthoFinder to extract orthologous groups (OGs) among the screened genes (Emms and Kelly 2019). Of the total 5,180 OGs identified, we found that 55 OGs contain one or more genes from all species. For example, a total of 168 *C. elegans* genes were assigned to any of the 55 OGs (Table S2). Genes involved in cuticle development were included in the 55 OGs (e.g., nematode cuticle collagen [col-125, col-33, col-133/OG0000005]), supporting the feasibility of detecting molting-related genes. Finally, we searched for transcription factor genes among 168 *C. elegans* genes, based on their domain structure. We found that only three OGs include transcription factor genes. Notably, although molting regulatory functions have not been reported in two OGs (OG0000071/Homeobox: *lin-39* and *nob-1*; OG0000130/zf-C2H2: *sptf-2* and *znf-782*), our screening results included *hr3* (OG0000063/Hormone_receptor, zf-C4). No other nuclear receptor genes were identified as candidate genes from this screening. Note that some transcription factors, such as *grh-1* and *blmp-1*, which are involved in the molting regulation of *C. elegans* (Meeuse et al., 2023), were not extracted in this screening (Fig. S11). For example, the peak expression of both genes before molting is comparatively conserved among the examined species; this pattern was not observed, or expression was only observed very weakly in *G. pallida* (Fig. S11).

In conclusion, our targeted approach and unbiased screening both detected *hr3/nhr-23* as a potent candidate conserved regulator of nematode molting. This gene potentially buffers the loss of multiple ECR genes and their functions.

### 4. Extensive expansion of nuclear receptors and their structural diversification in Rhabditina and Tylenchina

In order to understand the evolutionary background that *ecr* and *usp* genes are specifically lost within Rhabditina and Tylenchina among nematodes, we focused on that Rhabditina and Tylenchina extensively expanded nuclear receptor (NR) genes (Sural and Hobert 2021). The following analyses tested the hypothesis that the expansion of NR contributes to the specific losses of ECR and USP among nematodes.

#### Stepwise expansion of nematode NRs

We first reconstructed the evolutionary process of NR expansion. NRs were identified by extracting proteins with NR conserved domains from 160 nematode species. Plotting the number of NRs in the phylogenetic tree shows that NR expansion proceeded in a stepwise manner under phylogenetic constraints (Fig. 6A). We then extracted orthogroups (OGs) of NRs from represented 11 species (1–3 species from each Rhabditina, Tylenchina, Spirurina, Trichinellida, and Enoplida; Figs 6, S12). This analysis showed that the common ancestor of nematodes has at least 12 OGs, and NRs increased to 34 OGs in the ancestor of Rhabditida (Figs 6A, B, S12: Spirurina/Rhabditina/Tylenchina). Finally, the ancestor of Rhabditina/Tylenchina acquired additional 22 OGs (total 56 OGs), and Rhabditina and Tylenchina ancestrally had 136 and 75 OGs, respectively (Figs 6A, B, S12). Some nematodes such as *Plectus sambesii* also increased NRs (Fig. 6A), while *ecr* and *usp* are not missing. Thus, although expansions of NRs was not strictly associated with the loss of *ecr* and *usp* in some lineages, it is still important to point out that Rhabditina and Tylenchina are unique to have abundant NRs among nematodes.

**Figure 6.**
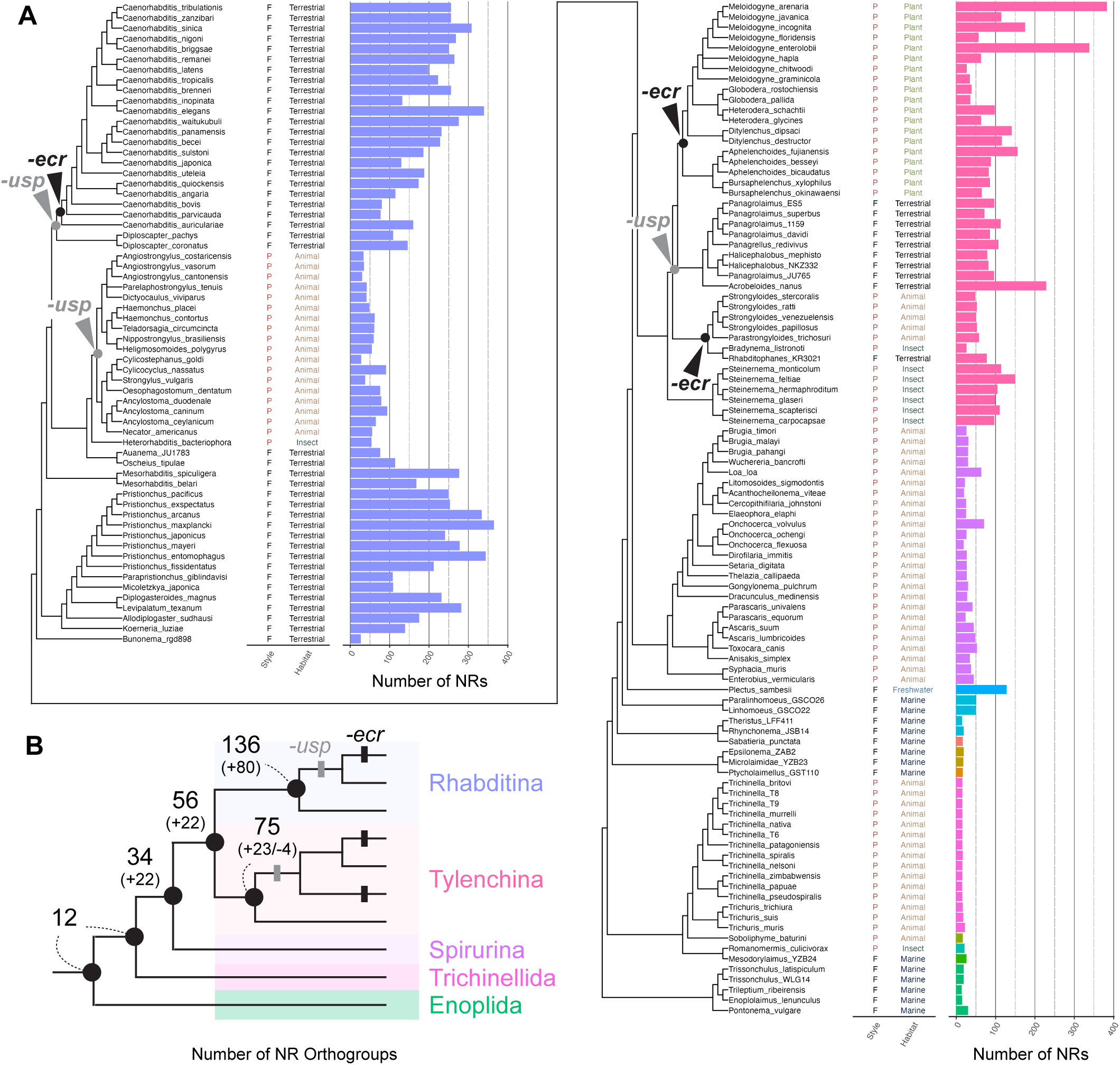
The evolutionary process of expansion of nematode nuclear receptors. **A**: Lifestyle (P: parasite and F: free-living), habitat, and the number of nuclear receptors (NRs) are indicated in the phylogeny of the target nematode species (color represents the taxa, see Fig. 1B). The nodes of *ecr* and *usp* loss events are highlighted in the tree. **C**: a simpler tree indicating the increase of NR orthogroups (OGs) at the nodes of representative taxa. The number above shows the number of OGs, and the number in the brackets below shows the increase (+) or decrease (-) of OGs at each node.

#### Structural divergence of expanded HNF4 proteins in nematodes

Since HNF4 subtype genes are specifically expanded in nematodes among various NR genes (Bonneton, et al. 2008; Kostrouchova and Kostrouch 2015), it is unclear if such HNF4 proteins contributed to the loss of different NR subtype proteins, ECR and USP. We assumed that the expanded HNF4 proteins diverged in structural functionality and convergently evolved a similar feature with ECR and USP. In order to test this hypothesis, we compared the structural features of ligand-binding domain (LBD), a core domain of NRs that interact with ligands and other proteins, between nematode “ancestral” and “expanded” HNF4s.

Nematodes ancestrally have a single HNF4 protein, and those proteins are conserved in early-branching taxa, such as Enoplida, Trichinnelida, and Monhynelida (Fig 2A, B). We predicted the protein structure of HNF4 LBD from 20 species of those groups using Colabfold/Alphafold2 (see the Methods). To investigate whether “ancestral” HNF4-LBDs have conserved HNF4-specific features, we compared each of the 20 predicted protein models with each NR LBD of the model insect *D. melanogaster* using TM-align (16 NRs, see also the Methods). The TM-score of each pairwise comparison was visualized as a heatmap in Fig. 7A. Highly specific and conserved LBD structural similarity is found between the nematode “ancestral” HNF4 and *D. melanogaster* HNF4 (Fig. 7A, B). Although some nematode HNF4s also show a comparatively high score with nonHNF4 NRs (Figs 7A, S13A), the TM-score from the comparison with *D. melanogaster* HNF4 is the highest among all species examined (Fig. S13B, C).

**Figure 7.**
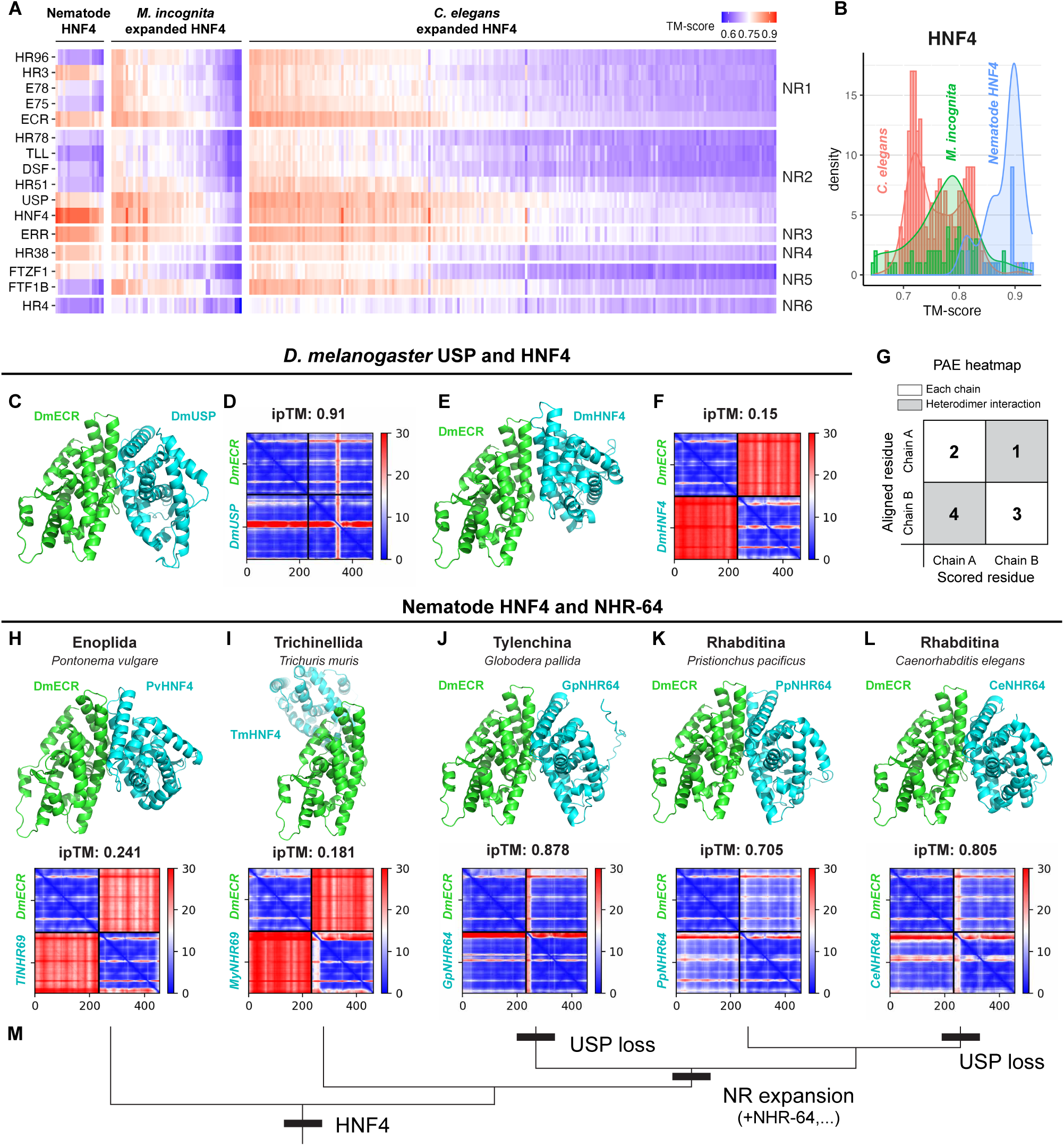
Structural divergence and the evolution of the heterodimer functionality of nematode HNF4 genes during NR expansion events. **A**: Heatmap of TM-scores for the pairwise structural alignment of the LBDs between nematode HNF4 and *Drosophila melanogaster* NR proteins. Each column represents the results of comparing a specific HNF4 protein from nematodes with each of the 16 *D. melanogaster* NRs (HR96, HR3, E78, etc., shown on the left y-axis). The NRs were also grouped into the subfamily which is indicated in the right y-axis. “Nematode HNF4” indicates HNF4 proteins from 20 species of early-branching taxa, such as Enoplida (see the main text). The center and right panels show the expanded HNF4 proteins extracted from the species *Meloidogyne incognita* and *Caenorhabditis elegans*, respectively. The x-axis labels (gene ID or name) are not shown in this heatmap, but a heatmap with labels is available in the supplementary dataset. **B**: Histogram of the TM score with *D. melanogaster* HNF4 in the three groups (blue: Nematode HNF4 [proteins from 20 species], *M. incognita*, and *C. elegans*). **C**–**F**: 3D models, ipTM values, and PAE heatmaps of the highest-scoring predicted heterodimer structure. All target proteins were investigated with ECR from *D. melanogaster* (DmECR). **C**–**D**: *D. melanogaster* USP (DmUSP), **E–F**: *D. melanogaster* HNF4 (DmHNF4), **H**: *Pontonema vulgare* HNF4 (PvHNF4), **I**; *Trichuris murris* HNF4 (TmHNF4), **J**: *Globodera pallida* NHR-64 (GpNHR-64), **K**: *Pristionchus pacificus* NHR-64 (PpNHR-64), and **L**; *Caenorhabditis elegans* NHR-64 (CeNHR-64). The ipTM values and PAE heatmaps are shown above the ipTM heatmaps. **M**: Phylogenetic tree of five species (*P. vulgare*, *T. muris*, *G.* pallida, *P. pacificus*, and *C. elegans*) showing the origin of HNF4 and NR expansion and loss of USP.

We conducted the same analysis on “expanded” HNF4s from Rhabditina and Tylenchina species. In this analysis, we predicted the structure of 260 and 60 HNF4 LBD which were extracted from *ecr*/*usp*-lacking Rhabditina *C. elegans* and Tylenchina *Meloidgyne Ingonita*, respectively (see also the Methods for the details). Unlike “ancestral” HNF4 proteins, more than 80% of examined proteins in both species do not necessarily show the highest TM-score with HNF4 among *D. melanogaster* 16 NRs (Fig. 7A, B).Notably, 52 (20%) and 85 (32.7%) proteins from *C. elegans* shows the highest TM-scores with ECR and USP, respectively (Fig. S13B). Similar results were also found in *M. Ingonita* (15 [25%] and 25 [41.6%], respectively, Fig. S13B). This analysis suggests the divergence of structural features of HNF4 LBDs during nematode NR expansion.

#### A case study finding nematode HNF4s which evolve a novel heterodimer functionality with ECR

The above analysis implied that some HNF4 proteins acquire new structural features during expansion events in Rhabditina and Tylenchina. To further test whether some HNF4 proteins evolved to functions equivalent to ECR or USP, we performed a case study to screen HNF4s which evolved USP-like functions in nematodes.

First, we searched for NRs with a structural feature similar to USP in the model nematode *C. elegans* using the Foldseek search tool (van Kempen, et al. 2024). The LBDs of ECR and USP have been experimentally determined to form a heterodimer (Hill, et al. 2013; Browning, et al. 2021). Thus, using the LBD of the USP protein from *D. melanogaster* as the query, we searched the AFDB-proteome database of *C. elegans* (UP000001940). As a result, we identified five candidate NRs with the top hit score (Table S3: NHR-64, NHR-69, NHR-35, NHR-14, and NHR-49). By investigating the presence of these proteins in other nematode species, we confirmed that all five proteins evolved from the HNF4 NR subtype within nematodes (=expanded HNF4s, Figs S14, S15F).

We used the ColabFold tool with AlphaFold2-Multimer to test if our target proteins could form a heterodimer with ECR (Jumper, et al. 2021; Evans, et al. 2022; Mirdita, et al. 2022). First, we validated that the ECR-USP heterodimer structure is correctly predicted in *D. melanogaster*. The predicted structural alignment of the ECR-USP heterodimer is highly consistent with the experimentally determined ECR-USP heterodimer (Fig. 7C) (Hill, et al. 2013; Browning, et al. 2021), and the ipTM (interface predicted template modeling) score is greater than 0.9. Furthermore, the PAE (predicted alignment error) heatmap clearly shows the high certainty of each ECR and USP prediction (Fig. 7D, second and fourth quadrants [Fig. 7G]) as well as the interaction of ECR and USP (Fig. 7D, first and third quadrants [Fig. 7G]). Conversely, a high-probability heterodimer interaction was not predicted when testing ECR and HNF4, which heterodimer functions have not been reported in *D. melanogaster* (Fig. 7E, F, ipTM = 0.15). The PAE heatmap shows that the interface of ECR and HNF4 is not accurately predicted (Fig. 7F; high error ratio [red color] in the first and third quadrants [Fig. 7G]), suggesting that the predicted alignment is likely due to random coincidence (Evans, et al. 2022). Thus, we confirmed that this analysis shows consistent results with the previous structural analysis.

We tested whether each candidate protein (NHR-64, NHR-69, NHR-35, NHR-14, and NHR-49) forms a heterodimer with *D. melanogaster* ECR. As shown in the analysis of ECR and HNF4 in *D. melanogaster*, heterodimer interaction was not clearly detected for three of the five proteins (Fig. S15A–E: NHR-35, NHR-14, and NHR-49; piTM = 0.209, 0.68, and 0.166, respectively). Notably, however, our results clearly predicted that NHR-64 and NHR-69 form a heterodimer with ECR (Fig. S15A, B, ipTM = 0.805 and 0.887, respectively). The structure of the putative heterodimer resembles that of the *D. melanogaster* ECR-USP heterodimer (Figs 7C, S15A-B). We furthermore performed yeast two-hybrid assay to validate the predicted heterodimer interaction of NHR-64 and NHR-69 in *in vivo* system (Fig. S15G, H). We first validated that our assay detected a weak and spontaneous interaction between ECR and USP without ligand in *D. melanogaster* (Fig. S15H), as shown in the previous studies (Lezzi, et al. 2002; Bitra and Palli 2009). We then investigated an interaction of *D. melanogaster* ECR and *C. elegans* NHR-64 or NHR-69 (Fig. S15H). Among the two proteins, interaction between *D. melanogaster* ECR and *C. elegans* NHR-64 was observed in this experiment.

We found that not only *C. elegans* but other Rhabditina and Tylenchina nematodes have orthologs of NHR-64 that exhibit similar structural features based on *in silico* structural prediction (Fig. 7J: *Globodera pallida* NHR-64 [ipTM = 0.878]; Fig. 7K: *Pristionchus pacificus* NHR-64 [ipTM = 0.705]; and Fig. S16B: Other examples). Our analysis also predicted that *C. elegans* NHR-64 can form a heterodimer with ECR of the nematode *Brugia malayi* (Fig. S16C). Moreover, we investigated the structural feature of “ancestral” HNF4 proteins (Figs 7H, I, and S16A). Interestingly, such HNF4s were not predicted to interact with ECR heterodimers (Fig. 7H: *Pontonema vulgare* [ipTM = 0.241]; Fig. 7I: *Trichuris muris* [ipTM = 0.181]; Fig. S16A: other examples). This is similar to the results of the ECR-HNF4 comparison in *D. melanogaster* (Fig. 6C). Taken together, our analyses suggests that some HNF4s evolved distinct structural features for dimerization during the NRs/HNF4s expansion within nematodes.

## Discussion

This study found that loss of *ecr* and *usp* is not unique to *Caenorhabditis* but has occurred multiple times within nematodes. Here we discuss that molting regulation by an alternative regulator and the expansion of NRs may explain the specific losses of *ecr* and *usp* within Rhabditina and Tylenchina species.

### Evolutionary scenario of nematode molting regulation

Based on the previous reports and our findings (ex., Figs 1, 3–4), it is implied that ecdysone is ancestrally involved in the molting regulation of Rhabditida (a clade of Spirurina, Rhabditina and Tylenchina) nematodes. Although it is still difficult to specify the regulatory role of ecdysone hormone in nematode molting (see the discussion below), involvement of ecdysone hormone plausibly changes in parallel with the losses of *ecr* and *usp* within Rhabditina and Tylenchina. We hypothesize that three aspects underlie this evolutionary process.

First, considering that molting of nematodes is consistent regardless of *ecr* and/or *usp* losses, the presence of other key molting regulators is implied. This study showed that HR3/NHR-23 is a potent candidate for the key regulator across nematode taxa. HR3/NHR-23 plays a key regulatory role in *C. elegans* molting, and this function may be conserved among a wide range of nematode taxa. Note that some components of *C. elegans* molting regulation, such as Lin-42, also show a conserved expression pattern during molting cycle among different nematodes (Fig. 5). This suggests that not only HR3 but also other genes may be involved in the molting of nematode ancestors. With such regulatory machinery, the loss of *ecr* and *usp* may be buffered without having catastrophic effects on the molting process itself.

Second, we propose that the loss of *usp* is a critical step toward the loss of *ecr*. This study revealed that *ecr* loss occurred only in lineages lacking *usp* (Fig. 2A). This may be because the loss of USP likely led to the altered regulatory interaction and functionality of ECR. Indeed, Graham et al., 2010 experimentally showed that ECR of *Haemonchus contortus* (usp-lacking Rhabditina) cannot form heterodimers with the USP of the red louse *Bovicola bovis*. Kostrouchova and Kostrouch 2015 also discussed changes of NR heterodimer functionality along loss of USP in *C. elegans.* <u>As shown in</u> our findings on the ECR-antagonist treatment and *ecr* expression in *usp*-lacking species (Figs 3, 4), the function of *ecr* in molting regulation may become minor or lost with the loss of *usp*. This is also consistent to that 20E treatment did not affect the L3-L4 molting of *H. contrtus* (Graham, et al. 2010). Although this hypothesis serves an interesting insight that usp-lacking species may represent an intermediate state of *ecr* loss, it is also important to note that some species of *usp*-lacking Rhabditina, such as *H. contortus* and *Heligmosomoides polygyrus*, are reported to exhibit endogenous ecdysone and pharmacological induction of molting by ecdysone (Dennis 1976; Fleming 1993). A detailed analysis of molting regulation in *usp*-lacking Rhabditina and Tylenchina is required for further assessing our hypothesis.

Finally, it seems critical that HNF4 subtype genes, the same NR subfamily with USP, have undergone extensive expansion in Rhabditina and Tylenchina (Bonneton, et al. 2008; Kostrouchova and Kostrouch 2015). This study suggests the divergence of the LBD structure of expanded HNF4 in *C. elegans* and *M. incognita* (Fig. 7A, B). We also screened a HNF4 protein which have a heterodimer functionality with ECR of *D. melanogaster* (Fig. 7C–M). These results are also consistent to the previous findings of the emergence of novel heterodimer pairs within expanded HNF4s in *C. elegans* (e.g., NHR-80 and NHR-49) (Pathare, et al. 2012). Based on these findings, it can be expected that expanded HNF4 can take over the function of USP, which may lead to USP loss and change in NR functionality. While this study did not address the causality of ECR loss, some HNF4 proteins may have convergently evolved the ECR-like structures/functions. Although NHR-64 represents an intriguing candidate for a representative protein which took over a USP-like function, considering that involvement of *nhr-64* in molting regulation is unclear (Gissendanner, et al. 2004; Liang, et al. 2010), further characterization of NHR-64 function and other expanded HNF4s across nematode species is required. In any case, this study raises the possibility that extensive NR expansion explains the specificity of *ecr*/*usp* losses among nematodes.

### Evolutionary impacts of structural divergence of nematode expanded NRs

The expansion of nematode NRs has been regarded as a “natural experiment made by a phylum” (Kostrouchova and Kostrouch 2015), and its evolutionary significance has been widely studied (Taubert, et al. 2011; Antebi 2015; Kostrouchova and Kostrouch 2015; Sural and Hobert 2021). For example, Sural and Hobert (2021) proposed that the expansion of NRs is associated with a free-living lifestyle. They reported that the expression of NRs, GPCRs, and insulin genes is enriched in the sensory neurons of *C. elegans* (Sural and Hobert 2021). Based on the findings, they hypothesizes that NRs may integrate sensory and/or internal information (Sural and Hobert 2021). It has also been suggested that NRs expanded in response to xenobiotics in free-living nematodes (Lindblom, et al. 2001; Lindblom and Dodd 2006). Notably, various other functions have been reported in addition to these examples (Taubert, et al. 2011; Antebi 2015; Kostrouchova and Kostrouch 2015). In this study, we found that NR expansion proceeded in a stepwise manner under phylogenetic constraints (Fig. 6) and that NRs evolved novel structural features (Fig. 7). We would like to discuss that our findings are not contradictory to previous research. For example, some parasitic lineages lost many NRs after the expansion of NRs in Rhabditina (Fig. 6A), which supports the association between NR expansion and a free-living lifestyle in this group. Additionally, our findings about changes in NR structural features shows affinity with the divergence of NR functions. We propose that changes in NR functionality, including heterodimeric potential, may be a driving force behind divergent NR functions in nematodes. Further investigation into the causes and consequences of NR expansion in nematode evolution is of great importance.

### Insights into evolution of nematode and ecdysozoan molting

Further experimental assessment of the role of the ecdysone hormone in nematodes is required for clarifying the evolutionary background of changes in the molting regulatory machinery. For example, the regulatory roles of *ecr and usp* need to be tested at the level of gene function in various nematodes. Recently, gene function assays have been established in several non-model nematode species, such as *Auanema*, *Steinernema*, and *Panagrolaimus* (Adams, et al. 2019; Cao 2023; Hellekes, et al. 2023). With this methodological development, it is critical to test if ecdysone/ecr regulates all four molting processes and it has a master regulatory role as seen in arthropods. Elucidating the ecdysone biogenesis process is also necessary. In particular, since ecdysone synthesis in nematodes is independent of Halloween genes (which are arthropod-specific P450 genes, Fig. 1B, Schumann et al., 2018), screening the P450 gene family is essential to identifying the genes responsible for ecdysteroid synthesis in nematodes. Finally, we note that ecdysone hormone likely has pleiotropic functions, including those related to reproduction, in nematodes. The relationship between such biological processes and the loss of ecr/usp is an important subject for future study.

Apart from ecdysone hormone, it is interesting that temporal expression of *hr3/nhr-23* during molting cycle is highly conserved among nematode species examined (Fig. 5B). Based on this finding, we provide the hypothesis that HR3 plays a dominant role as an ancestral and conserved molting regulator in nematodes and possibly even in ecdysozoans (note that HR3 is also a core molting regulator in arthropods (Yamanaka, et al. 2013) and putatively in tardigrades (Yamakawa and Hejnol 2024)). HR3 is also highly conserved among ecdysozoan species (Fig. S6). Considering that HR3 orthologues ROR and NHR-23 are known to be involved in regulation of circadian clocks among bilaterians (Patke, et al. 2020; Hiroki and Yoshitane 2024; Patil and van Zon 2024), the ancestry of HR3 shows an association with the origin of molting regulation, which control biological and developmental time. It should be also noted that other circadian clock components such as lin-42/per also shows a conserved temporal expression among different nematode species (Fig. 5). Our results also provide the possible use of HR3 regulation of molting for medical applications and pest control, in plant-parasitic and some human-parasitic nematodes, which have lost ECR (Fig. 2A: Tylenchomorpha and Strongylida, respectively).

## Materials and Methods

### 1. Data collection and target species

We obtained protein sequences of 159 nematode species from WormBase Parasite (version: WBPS19, WS291) (Howe, et al. 2017). These sequences/protein models were reconstructed from the genomic data. All accession numbers of the original dataset are available in the supplementary dataset. In this study, the gene ID was referenced to the original dataset, while also incorporating some modifications. For example, species identifiers were added to the beginning of the gene ID (i.e. *GeneID* → *NEMA_CaenoElega_GeneID*). Some characters, such as curly brackets, were also removed for later analysis. In addition, the transcriptome data of the Enoplida nematode *Pontonema vulgare* was obtained from NCBI (accession number: PRJNA504396). Quality of raw reads were examined using FastQC (version: v0.11.9; https://www.bioinformatics.babraham.ac.uk/projects/fastqc/). The subsequent filtering was performed using trimmomatic (version: 0.39) (Bolger, et al. 2014), and *de novo* assembly was conducted using Trinity (version: v2.13.2) (Henschel, et al. 2012). Transdecoder (version: 5.5.0; https://github.com/TransDecoder/TransDecoder) was used to obtain the peptide sequences of *P. vulgare* transcript models. The assembled sequences are available in the supplementary dataset.

The phylogenetic relationship of the target species was referenced to previous studies. For example, the topology of the major taxon phylogeny (grouped by color in Fig. 2) is consistent with three recent molecular phylogenetic reports (Smythe, et al. 2019; Ahmed, et al. 2022; Qing, et al. 2024). The relationship of Rhabditina and Tylenchina taxa is also consistent in the three reports (shown in Fig. 2C). Since the analysis by Ahmed et al. (2022) used the largest number of our target species, we modified their phylogenetic tree by adding our target species with referring to other literatures. If our target species were not included in their analysis, we referred to other literature. For example, we used Ahmed et al. (2022) to reconstruct the phylogeny of *Caenorhabditis* species, but the position of *C. auriculariae* was unclear in that study. In this case, we determined the position of *C. auriculariae* based on another study (Dayi, et al. 2021). The phylogenetic positions of the following species were determined not only with reference to Ahmed et al., 2022: *Angiostrongylus cantonensis* (Dumidae, et al. 2023), *Cylicocylus radiatus* (Hu, et al. 2020), *Meloidogyne chitwoodi* (Álvarez-Ortega, et al. 2019), *Steinernema hermaphroditum* (Schwarz, et al. 2025), *Aphelenchoides* species (Lai, et al. 2022), *Cercopithifilaria johnstoni* (McCann, et al. 2021), and Enoplida species (Qing, et al. 2024). The phylogeny of the two species, *Parelaphostrongylus tenuis* and *Bradynema listronoti*, was unclear. Their phylogenetic positions were estimated by identifying their respective family, Metastrongylidae and Allantonematidae, from the NCBI Taxonomy database (https://www.ncbi.nlm.nih.gov/taxonomy). A Newick file for visualizing the phylogenetic tree is available in the supplementary dataset. We also used the trees which were reconstructed by Ahmed et al., 2022 to test the multiple losses of *ecr* and *usp* in Rhabditina and Tylenchina. These files are available in the supplementary data of Ahmed et al., 2022, which was uploaded in Figshare (https://figshare.com/s/946b2bc6aef7ce4a9e6a, 3_TYLENCHINA and 4_RHABDITINA).

### 2. Gene identification

#### Screening

Gene mining of *ecr*, *usp, hr3, e75, ftz-f1*, *lin-42*, and *kin-20* was conducted based on the sequence similarity by combination of BLAST search and phylogenetic analysis with the following screening. All sequences and phylogenetic trees are available in the supplementary datasets.

We first performed a BLAST search using the query sequences of ECR, USP, HR3, E75, and FTZ-F1 from *Drosophila melanogaster* against the database of each nematode species. The query sequences are from UniProt (ECR: P34021, USP: P20153, HR3: P31396, E75: P17672, and FTZ-F1: P33244). The peptides showing the hit with less than 1E-20 e-value were extracted and used as query sequences for the second BLAST against the *D. melanogaster* database. This database was built using the CDS sequences from FlyBase (dmel-all-CDS-r6.45.fasta) (Öztürk-Çolak, et al. 2024). For the screening of LIN-42 and KIN-20, we used the query sequence and the database from *Caenorhabditis elegans* (UniProt id: Q65ZG8 and Q20471, respectively). Finally, the peptides showing the reciprocal top hit with less than 1E-10 e-value were extracted. Each candidate gene set of ECR, USP, HR3, E75, and FTZ-F1 (181, 256, 342, 196, and 199 peptides, respectively) was aligned with other known nuclear receptors (NRs) from different animals (the sequences were previously used to identify NR genes (Vogeler, et al. 2014; Yamakawa and Hejnol 2024) (see also the supplementary dataset) using MAFFT (version: v7.526, – –auto option) (Katoh and Standley 2013), and the alignments were filtered using trimAl (version: 1.4.1, threshold: 0.8) (Capella-Gutiérrez, et al. 2009). Candidate 203 proteins of LIN-42 were also aligned with other known Period, Clock, Arnt, and Bmal proteins using the same methods (the sequences are available in the supplementary dataset). Candidate 303 proteins of KIN-20 were aligned with KIN-19 and CSNK-1 proteins. Phylogenetic analysis of each data set was performed using RAxML (version: 8.2.12, bootstrap: 1,000 generations, raxmlHPC-PTHREADS-SSE3 –f a –x 12345 –p 12345 –# 1000 –m PROTGAMMAAUTO—-auto-prot=aic –s *input_file* –n *output_file*) (Stamatakis 2014). Gene identity was determined by examining, whether the candidate genes were included in the clade of target genes with high bootstrap values. By removing the genes that did not belong to the target clade, we finally identified 128, 74, 276, 151, 199, 132, and 220 peptide models as ECR, USP, HR3, E75, FTZ-F1, LIN-42, and KIN-20 respectively (see the trees in the supplementary dataset). Note that some species have multiple copies.

Next, the above identified target peptides were used as query sequences, and a BLAST search was performed against the database of each nematode species in which the target genes were not found in the first screening. Top-hit sequences with less than 1E-20 e-values were collected, and each sequence was used as a query for the second BLAST search against the database of the original species from which the first query originated. Then, genes showing the reciprocal one-to-one top-hit were finally accumulated as candidates (ECR: 197, USP: 277, HR3: 28, E75: 77, FTZ-F1: 95, LIN-42: 151, and KIN-20: 2 peptides). After selecting the peptides having DNA binding domain of NRs (except LIN-42 and KIN-20, HMMER domain search [version 3.4] (Johnson, et al. 2010), see the below for the details) and manually checking the accuracy of the gene domain (e.g., NCBI conserved domain search: https://www.ncbi.nlm.nih.gov/Structure/cdd/wrpsb.cgi), we finally investigated the identity of the selected peptides by phylogenetic analysis as mentioned above. In this analysis, 5, 21, 1, 16, 28, 17 and 0 peptide(s) were additionally identified as ECR, USP, HR3, E75, FTZ-F1, LIN-42, and KIN-20, respectively. Finally, the peptides obtained in the 2nd screening were used for the BLAST query sequences, and we repeated the BLAST analysis as performed in the 2nd screening. However, no genes were newly identified as ECR, USP, HR3, FTZ-F1, and LIN-42. Only 4 peptides were additionally identified as E75.

Target genes were also searched by BLAST, Hmmer domain search (screening of the genes having DNA or ligand binding domain of NRs, see also the below), and phylogenetic analysis. For example, the BLAST search was performed with a higher e-value threshold (ex. 1E-05). Different database was also explored for the species missing the target genes, and ECR of *Steinernema feltiae* was identified using the other dataset in WormBase Parasite (version: WBPS19, WS291) from the one which we originally used (accession number: PRJNA204661). Finally, eight additional genes (nine peptides) were identified to compensate for the missing target genes, and the results are summarized in Fig. 2. Note that no *ecr* and *usp* genes were identified in the tblastn-based search on the genome assembly of the species lacking these genes.

#### Criteria for the presence and absence of target genes

The summary of the screening and gene identification of 160 species is shown in Fig. 2A. In some species, the target genes were missing even though the genes were found in species of the same genus (or closely related species). Since this study avoided to distinguish between species-specific loss and technical errors such as low data quality, gene missing in these species were considered as missing information in Fig. 2A (question marks in Fig. 2A). We considered the loss to be reliable if the loss of the genes was observed in all the same genus and closely related species. According to these criteria, we detected loss of *ecr* only in *Caenorhabditis*, Tylenchomorpha, Strongyloididae and Alloionematidae (Fig. 2A). Loss of *usp* was also reliable in Strongylida, Anarhabditis, non-Steinernematidae Tylenchina (Fig. 2A).

#### Gene identification in other ecdysozoans

Previous works identified *ecr*, *usp* and *hr3* in insects (*Drosophila melanogaster*, *Drosophila pseudoobscura*, *Anopheles gambiae*, *Aedes aegypti*, *Bombyx mori*, *Tribolium castaneum*, and *Apis mellifera*), other arthropods (Myriapoda: *Strigamia maritima*; Chelicerata: *Limulus polyphemus* and *Stegodyphus mimosarum*; and Branchiopoda: *Daphnia pulex* and *Daphnia magna*), tardigrades (*Ramazzottius varieornatus* and *Hypsibius exemplaris*), onychophorans (*Euperipatoides rowelli*), nematodes (*Brugia malayi*, *Caenorhabditis elegans* [only *hr3*]), and priapulids (*Priaplus caudutus*) (Bonneton, et al. 2008; Schumann, et al. 2018). In addition, we newly identified *ecr*, *usp*, and *hr3* in tardigrades (*Echiniscus testudo*), nematomorphs (*Acutogordius australiensis*, *Gordionus montsenyensis*, *Nectonema munidae*, *Paragordius varius*), kinorhynchs (*Campyloderes vanhoeffeni*, *Franciscidere kalenesos*, *Dracoderes abei*) and priapulids (*Tubiluchus corallicola*). For gene identification, we collected genomic or transcriptomic data from publications or public databases (*E. testudo*: Murai et al., 2021 (Murai, et al. 2021), *A. australiensis* and *N. munidae*: Cunha et al., 2023 (Cunha, et al. 2023) [FigShare: doi.org/10.6084/m9.figshare.23419922], *G. montsenyensis*: Eleftheriadi et al., 2024 (Eleftheriadi, et al. 2024), *P. varius*: ERR1817116, ERR1817117, ERR1817118, and ERR1817119 [NCBI SRA]; kinorhynchs: Herranz et al., 2022 (Herranz, et al. 2022) [*C. vanhoeffeni*: SRR14509480, *F. kalenesos*: SRR14509483, and *D. abei*: SRR14509486]). Raw read data from *P. varius* and kinorhynch species were *de novo* assembled as described above. Gene identity was confirmed by BLAST and phylogenetic analysis using the same methods as above. The results are summarized in Fig. S6, and the gene tree and sequences are available in the supplemental dataset.

#### Reannotation of the ecr genes identified by Gonzalez Akimori et al., 2021

The seven ECR-1 genes used to identify ECR-1 of *Strongyloides stercoralis* in Gonzalez Akimori et al. 2021 were obtained from NCBI GenBank (Gonzalez Akimori, et al. 2021). The sequence of ECR-1 of *S. stercoralis* was not available due to unknown accession number after their gene name modification (Gonzalez Akimori, et al. 2021). Phylogenetic analysis was performed as described above, and the results are summarized in Fig. S2. We also included the sequences of the ecdysozoan THR (Thyroid hormone receptor) genes after preliminary test using the above dataset. All sequences for the tree are available in the supplemental dataset.

#### Genome assembly quality

We obtained the quality metrics information for the genome assembly used in this study from WormBase Parasite (version WBPS19, WS291) (Howe, et al. 2017). These metrics include genome size, N50, scaffold number, BUSCO annotation score, and BUSCO assembly score. These metrics are summarized in the phylogenetic trees in Figs S3–S5.

### 3. Extraction of NRs and transcription factors based on the domain conservation

The HMM profile of NRs (Hormone_recep: PF00104) was obtained from the Pfam database. Using this profile as a query, hmmscan was performed against all peptide data of the target species (threshold 1E-05) (Johnson, et al. 2010), and the identified genes were considered as NRs. The number of genes was counted considering the isoforms and shown in the linear bar graph of Fig. 5. We also used previously generated HMM profiles to screen transcription factors (Yamakawa, et al. 2023). Shortly, the profiles include the following targets: ARID (PF01388.17), AT_hook (PF02178.15), Basic (PF01586.12), CUT (PF02376.11), DM (PF00751.14), Ets (PF00178.18), Forkhead (PF00250.14), GATA (PF00320.23), GCM (PF03615.11), HLH (PF00010.22), HMG_box (PF00505.15), Hairy_orange (PF07527.9), Homeobox (PF00046.25), Hormone_recep (PF00104.26), OAR (PF03826.13), P53 (PF00870.14), P53_tetramer (PF07710.7), PAX (PF00292.14), Pou (PF00157.13), HPD (PF05044.8), RHD_DNA_bind (PF00554.18), Runt (PF00853.15), SCAN (PF02023.13), SIM_C (PF06621.8), SRF-TF (PF00319.14), T-box (PF00907.18), TBX (PF12598.4), TF_AP-2 (PF03299.10), TF_Otx (PF03529.9), bZIP_1 (PF00170.17), bZIP_2 (PF07716.11), zf-C2H2 (PF00096.22), zf-C2HC (PF01530.14) and zf-C4 (PF00105.14)

### 4. OrthoFinder analysis

OrthoFinder (version: 2.5.4, with default parameters) (Emms and Kelly 2019) was used to examine the orthogroups (OGs) of NRs in the represented nematode species of five major groups (Fig. 6): Rhabditina (*Ancylostoma ceylanicum*, *Caenorhabditis elegans*, *Heligmosomoides polygyrus*, and *Pristionchus pacificus*), Tylenchina (*Globodera pallida*, *Meloidogyne arenaria*, and *Steinernema monticolum*), Spirurina (*Ascaris suum* and *Brugia malayi*), Trichinellida (*Trichuris muris*), and Enoplida (*Pontonema vulgare*). Peptide sequences of NRs from each species were obtained as described above. After identifying 163 OGs in this comparison, we investigated the taxonomic origin of the genes in each OG (Rhabditina, Tylenchina, Spirurina, Trichinellida, and Enoplida). If the OG contained one or more genes from all taxa, the OG was considered to be the one that the common ancestor of the taxa had (Fig. 6B). Using these criteria, the evolutionary gain/loss events of NR OGs were resolved, as shown in Fig. 6B. Output file of the analysis is available in the supplementary dataset.

### 5. Transcriptome analysis

#### Data collection

We collected transcriptome data, including molting stages, for a total of six nematode species from public databases. The expression matrix (RPKM) for *Caenorhabditis elegans* was obtained from Kim et al., 2013 (Gene Expression Omnibus, accession number: GSE49043) (Kim, et al. 2013). WormBase ParaSite (version: WBPS19, WS291) (Howe, et al. 2017) was also the resource of this study to obtain the calculated expression matrix (TPM) for *Trichuris suis* (NCBI Sequence Read Archive [SRA] accession number: SRP039506) and *Globodera pallida* (NCBI SRA accession number: ERP001236). The expression matrix (TPM) for *Pristionchus pacificus* was obtained from the supplementary file of Sun et al. 2021(Sun, et al. 2021). In addition, raw read data of *Brugia malayi* and *Ancylostoma ceylanicum* were obtained from SRA (NCBI SRA, accession number: SRR4308250-51 and SRR9858310-17 [*B. malayi*] (Grote, et al. 2020) and SRR2125572, SRR2125581-86, SRR2125588-90, SRR2125594, SRR2125597, SRR2125600, SRR2125602, SRR2125606-7, SRR2125610-SRR2125611, SRR2125614, and SRR2125627 [*A. ceylanicum*] (Bernot, et al. 2020)), and the expression levels were calculated by us. As mentioned above, quality control and filtering of raw reads were performed by FastQC and trimmomatic, respectively. We also used samtools (version: 1.15.1) to remove the ribosomal RNA-derived sequences in the *B. malayi (Danecek, et al. 2021)*. The filtered reads were mapped to the reference transcript models (*B. malayi*: PRJNA1072 and *A. ceylanicum*: PRJNA231479, WormBase Parasite (Howe, et al. 2017)) using RSEM/bowtie2 (version: v1.3.1, rsem-calculate-expression ––paired-end ––bowtie2) to calculate expression levels (TPM) (Li and Dewey 2011). Molting stages were referred from original/previous papers (Kim, et al. 2013; Cotton, et al. 2014; Leroux, et al. 2018; Bernot, et al. 2020; Grote, et al. 2020; Sun, et al. 2021; Shears and Grencis 2022). Expression levels of the transcripts corresponding to the *ecr* and *hr3,* which were identified in the above screenings, were obtained from each transcriptome dataset (the average of TPM was calculated in the case of having replicates at the same stage). In the transcriptome data of *G. pallida*, the gene model of *hr3* was inappropriately reconstructed into two models (id: GPLIN_000052400 and GPLIN_000052600). Thus, the expression levels were calculated by summing the expression levels of the two models at each time point. *grh-1* and *blmp-1* were annotated by reciprocal blast search (*B. malayi*: Gene:WBGene00226940 [*grh-1*], Gene:WBGene00223835 [*blmp-1*]; *A. ceylanicum*: Acey_s0568.v2.g2260 [*grh-1*], Acey_s0580.v2.g13356 [*blmp-1*]; *T. suis*: D918_02299 [*grh-1*], D918_01175 [*blmp-1*]; *P. pacificus*: PPA28052 [*grh-1*], PPA04978 [*blmp-1*]; *G. pallida*: GPLIN_000211500 [*grh-1*], GPLIN_000875200 [*blmp-1*]; and *C. elegans*: grh-1 [*grh-1*], blmp-1 [*blmp-1*]). Their expression was investigated using the same dataset/methods (Fig. S11).

#### Data curation

Sun et al., 2021 originally provided the 1–2 h interval transcriptome data of *P. pacifucs* from 0 to 58 h after hatching (i.e., 0, 1, 3, 5, … 58 h) (Sun, et al. 2021). In our preliminary analysis, we found that the data contained an overall fluctuating expression pattern throughout the examined stages (Fig. S9A). Therefore, we divided the time points into overlapping windows of three time points each, and the mean was calculated for each window. For example, the first time-window was grouped by the data at 0, 1, and 3 h, and the mean value represents the data at 1 h. The second window included the data at 1, 3, and 5 h, and the mean value represents the data at 3 h. Using this method, we obtained the corrected expression profiles, which tends to remove the effects of fluctuating expression due to biological or technical differences on individual development (Fig. S9B).

#### Unbiased screening of NRs for identifying molting regulators

With reference to our previous work (Yamakawa and Hejnol 2024), the expression profiles (TPM matrices) of four nematode transcriptome data were classified into 5–7 clusters using fuzzy c-mean clustering (Fig. S10: *A. ceylanicum*, *G. pallida*, *B. malayi*, and *T. suis*). Analysis was carried out using R and its package e1071 (version: 1.7.11: https://cran.r-project.org/web/packages/e1071/index.html). From such clustering, the genes in the cluster(s) that showed a positively associated expression pattern with molting cycles were extracted (Fig. S10, green shades, *A. ceylanicum*: 8110, *G. pallida*: 3383, *B. malayi*: 7114, and *T. suis*: 2896 genes). In addition, Sun et al. 2021 and Kim et al. 2013 extracted the genes whose expression showed an oscillatory pattern along the molting cycle in *P. pacificus* (2,964 genes) and *C. elegans* (1,592 genes: assigned to clusters 1–6 in Kim et al., 2013), respectively (Kim, et al. 2013; Sun, et al. 2021). As described above, OrthoFinder (Emms and Kelly 2019) was used to extract OGs among the screened genes, and this result identified 5,180 OGs. Among 5,180 OGs, 55 OGs contain one or more genes from all species. We extracted 168 genes which were assigned into any 55 OGs in *C. elegans.* Finally, we searched for the presence of TF and NR domains in the represented the *C. elegans* 168 genes using Hmmer search (Johnson, et al. 2010) (see above).

### 6. Prediction of heterodimer structure

#### Foldseek search for the identification of the candidate proteins

First, we extracted the ligand-binding domain of USP from *Drosophila melanogaster* (UniProt: P20153) using the NCBI Conserved Domain Search tool (https://www.ncbi.nlm.nih.gov/Structure/cdd/wrpsb.cgi) (Blum, et al. 2024). We performed a Foldseek Search on the Foldseek web server (https://search.foldseek.com/search) and predicted the three-dimensional structure of USP with the 3Di/AA mode using sequence information (van Kempen, et al. 2024). We used this prediction as a query and targeted the AFDB-PROTEOME database in *C. elegans* (UP000001940). The results of this search are shown in Table S3. Subsequent analysis focused on five candidate proteins (NHR-64, NHR-69, NHR-35, NHR-49, and NHR-14). The orthologous genes were identified in several nematode species using BLAST and phylogenetic analyses (Enoplida: *Pontonema vulgare*, *Trissonchulus latispiculum*; Trichinellida: *Trichuris murris*; Dorylaimida: *Mesodorylaimus YZB24*; Spirurina: *Ascaris suum* and *Brugia malayi*; Tylenchina: *Globodera pallida*, *Steinernema monticolum*, and *Meloidogyne arenaria*; and Rhabditina: *Caenorhabditis elegans*, *Heligmosomoides polygyrus*, *Ancylostoma ceylanicum*, and *Pristionchus pacificus*). The phylogenetic tree is available in the supplementary dataset, and Fig. S14 shows part of the tree (the USP and HNF4 clade).

#### Prediction of heterodimer interaction

The ColabFold interface was used to perform AlphaFold2 and AlphaFold2-Multimer analyses (Evans, et al. 2022; Mirdita, et al. 2022). The sequences used in the analysis were obtained from the above analysis or from UniProt (*D. melanogaster* ECR: P34021 and HNF4: P49866). The ligand binding domains were extracted using the NCBI Conserved Domain Search tool. The sequences were assigned to the ColabFold interface and analyzed with the default settings (e.g., num_relax = 0 and template_mode = none). Five protein structure models were predicted in each analysis. We presented the 3D model and ipTM score of the most reliable model in the main text (Figs. 7 and S1). The 3D models of the proteins were generated using PyMOL (version 3.1, https://www.pymol.org/), and the heat maps were obtained from the ColabFold output files. The original output files of this analysis are available in the supplementary dataset.

### 7. TM-alignment

#### Extraction and prediction of NR and HNF4 LBDs

The predicted protein structure datasets for *C. elegans* (UP000001940_6239_CAEEL_v6) and *D. melanogaster* (UP000000803_7227_DROME_v6) were obtained from the AlphaFold Protein Structure Database (Fleming, et al. 2025). The expanded HNF4 genes of *C. elegans* were extracted by removing eight conserved non-HNF4 NR genes (*nhr-6*, *nhr-8*, *nhr-23*, *nhr-25*, *nhr-41*, *nhr-67*, *nhr-85*, and *nhr-91*) from the NR genes annotated as nuclear hormone receptors (*nhr-1*, *nhr-2*, …). The NRs of *D. melanogaster* were extracted based on previous annotations in UniProt (P16376: 7UP2, Q9VML1: DSF, P13055: E75, P45447: E78, and P34021: ECR, Q8WS79: ERR, Q05192: FTF1B, and P33244: FTZF1, P49866: HNF4, P31396: HR3, P49869: HR38, Q9W539: HR4, A1ZA01: HR51, Q24142: HR78, Q9VI12: HR83, Q24143: HR96, P18102: TLL, and P20153: USP). We used the NCBI Conserved Domain Search (https://www.ncbi.nlm.nih.gov/Structure/cdd/wrpsb.cgi) with the default settings to identify the NR LBD (ligand binding domain) in each protein sequence. Note that we removed genes with LBD regions of 150 amino acids or fewer to focus on genes with functional LBDs and reduce artificial errors. Finally, we extracted the structures of the specific LBD regions from each dataset using pdb-tools (218 and 16 proteins in *C. elegans* and *D. melanogaster*, respectively) (Rodrigues, et al. 2018).

Unlike *C. elegans* and *D. melanogaster*, the predicted protein structure of *Meloidogyne incognita* was not available in a public database, so we predicted the protein structure of *M. incognita* HNF4s. As mentioned above, 175 NRs were extracted from the *M. incognita* genome, and a phylogenetic analysis was performed on all NRs and reference NR proteins (500 bootstrap replications, the reconstructed tree is available in the supplementary dataset). We removed 24 non-HNF4 NRs, and the LBD regions were investigated in the remaining 151 proteins. Subsequently, LBD sequences were obtained for 64 proteins with more than 150 amino acid residues in the LBD. We predicted each protein structure using ColabFold/Alphafold2, as mentioned above. We applied the same analysis to HNF4 proteins from other nematodes. For example, we extracted HNF4 proteins from 28 species of Chromadorida, Desmodorida, Dioctophymatida, Dorylaimida, Enoplida, Mermithida, Monhysterida, and Trichinellida. Gene identity was confirmed by BLAST-based search (ex., reciprocal tophit with *D. melanogaster* HNF4). Finally, the HNF4 LBD structure was predicted in 20 species (*Epsilonema ZAB2, Pontonema vulgare, Ptycholaimellus GST110, Rhynchonema JSB14, Soboliphyme baturini, Trichinella T6, T. T8, T. T9, T. britovi, T. murrelli, T. nativa, T. nelsoni, T. papuae, T. patagoniensis, T. pseudospiralis, T. zimbabwensis, Trissonchulus WLG14,* and *Trissonchulus latispiculum*). In subsequent analyses, we used the best-predicted models for each prediction.

#### Scoring of TM-align

Using the above three protein structure datasets (nematode “ancestral” HNF4, *C. elegans* expanded HNF4, and *M. incognita* expanded HNF4 proteins), we performed pairwise TM-align (version: 20220412) analysis with reference *D. melanogaster* NR proteins (Zhang and Skolnick 2005). The analysis was performed without any additional options. We used the normalized TM-score with nematode HNF4 proteins to generate a TM-score heatmap and histograms. The results, including the script and raw TM-scores, are available in the supplementary dataset.

### 8. Yeast two-hybrid assay

A yeast two-hybrid assay was performed using the Matchmaker Gold Yeast Two-Hybrid System (Takara Bio, Shiga, Japan). In brief, the target sequences were inserted into the bait and prey vector plasmids (pGBKT7 and pGADT7, respectively). The plasmids were then co-transformed into the Y2HGold yeast strain and cultured on selective agar plates.

#### Amplification of the target sequences and plasmid construction

We commercially purchased *D. melanogaster* from Easy Zoo GmbH (Minden, Germany), and 5–15 pupae and adults were homogenized with liquid nitrogen and TRIzol Reagent (Invitrogen, MA, USA) to extract total RNA. The total RNA was collected using 1-bromo-3-chloropropane and further purified using isopropanol precipitation methods. After DNase treatment (Thermo Scientific, MA, USA), the RNA solution underwent further purification with ethanol precipitation. We applied the same methods to *C. elegans* worms of mixed stages. Reverse transcription and cDNA synthesis were conducted using a RevertAid First Strand cDNA Synthesis Kit (Thermo Scientific, MA, USA) by following the provided protocol. LBDs of *D. melanogaster ecr*, *usp*, *hnf4*, *C. elegans nhr-64*, and *nhr-69* were predicted as mentioned above. The primers used to amplify the LBD regions with additional sequence for plasmid construction are shown in Table S5. The pGBKT7 and pGADT7 plasmids were linearized by inverse PCR using the specific primers designed in the multiple cloning sites (see Table S5). The amplified and purified DNA fragments were cloned into the linearized plasmids using an In-Fusion Cloning Kit (Takara Bio). Accuracy of the insertion was confirmed by colony PCR and direct sequencing.

#### Co-transformation and detection of the protein interactions

The above plasmids were co-transformed into the yeast Y2HGold using the Yeastmaker Yeast Transformation System 2 (Takara Bio). The procedure was followed by the provided protocol. Liquid YPDA medium was used to grow the yeast cultures, and the transformation was performed with a standard lithium acetate method. See the main text for the bait/prey pairs of co-transformations. We used pGBKT7-53, pGBKT7-Lam, and pGADT7-T vectors, which were provided in the same kit, for positive/negative control. As shown in Fig. S15G, Y2HGold is auxotrophic for adenine (Ade), histidine (His), leucine (Leu), and tryptophan (Trp), and pGBKT7 and pGADT7 vector complements Trp and Leu, respectively. Thus, the co-transformants were first obtained by streaking the treated yeasts into the SD-Leu-Trp, and well-grown colonies were subsequently streaked onto the selective agar plates to test the expression of other reporter genes. In our preliminary screening, the test using the reporter gene *ADE2* is too strict to detect the interaction of our target proteins. For example, although the colonies of the co-transformants of p53-T were found on SD-Leu-Trp-His-Ade, no colonies of the co-transformants of DmECR-DmUSP, whose interaction was previously reported, were found. Thus, we decided to target the HIS3 and MEL1 reporter genes, whose stringency is comparatively mild, by using SD-Leu-Trp-His/X-alpha-gal plates (Fig. S15G). Specifically, the co-transformants were spread on SD-Leu-Trp for three days, and each of 6–12 well-grown colonies were further stroked onto SD-Leu-Trp/X-alpha-gal. We tested whether the colonies further amplify and turn to blue color within 24–48h. We used the following materials for making the agar plates: Minimal SD Base (Takara Bio), DO Supplement –Leu/-Trp (Takara Bio), DO Supplement –Ade/-His/-Leu/-Trp (Takara Bio), 5-Bromo-4-chloro-3-indolyl α-D-galactopyranoside (=X-alpha-gal, Sigma-Aldrich), and Adenine Hemisulfate Salt (Sigma-Aldrich).

### 9. Culture of nematodes and pharmacological experiments

#### Culture of C. elegans, D. pachys, and P. sambesii

We used *C. elegans* (N2 lineage), which were kept at the Institute of Nutritional Science in Friedrich Schiller University Jena, Germany. *D. pachys* and *P. sambesii* were obtained from Prof. Dr. Ralf Schnabel (Institute of Genetics, TU Braunschweig, Germany). They are cultured at room temperature on the optimized agar plates for each species. For *C. elegans,* we used a standard NGM/nematode growth media plate (Stiernagle 2006). *D. pachys* were cultured on the 2% agar plate in tap water, which was obtained in our lab (Institute of Zoology and Evolutionary Research, Friedrich Schiller University Jena, Germany). The agar solution was poured onto the plates without adding any materials after autoclaving. The plate for *P. sambesii* was made of 2% agar in distilled water. 5 g/L cholesterol (Carl ROTH, Karlsruhe, Germany) was made in 96% ethanol, and the 1/1000 amount of cholesterol solution was added to the agar solution after autoclaving. All of the three species are bacterial feeders, and we provided the *E. coli* OP50 to *C. elegans* and *P. sambesii*, and *E. coli* HT115 to *D. pachys*. The above cultivation methods for *D. pachys* and *P. sambesii* was kindly advised by Prof. Dr. Ralf Schnabel. Except the above differences in agar plates and bacterial food, manipulation and maintenance were followed by the manual of *C. elegans* (Stiernagle 2006).

#### Collection of synchronous L1 worms

With reference to the methods in *C. elegans* (Stiernagle 2006), synchronous L1 worms of each species were obtained by bleaching gravid adults or eggs, which were laid on the agar plates. A bleaching solution was made of a 2:1:2 mix solution of a hygiene cleaner, which was commercially purchased (DanKlorix, Colgate-Palmolive Deutschland GmbH, Germany), 10N NaOH solution, and M9 buffer, respectively. The worms and eggs were collected by washing the plate surface with 2–5 ml M9 buffer (several hundred-thousand worms in total). We also used a drawing brush to effectively collect the eggs which were laid beneath the agar surface in *D. pachys* and *P. sambesii*. The M9 buffers with worms and eggs were washed with M9 buffer two times using a centrifuge (12,000 g; for 30–60 sec), and finally, the solution was concentrated to around 100–500 µl. Two-times volume of the freshly made bleaching solution was added to the concentrated M9 buffer. The bleaching process was carried out in the 1.5 ml tube or small Petri dishes. In any case, the solutions were periodically shaken with pipetting or vortexing, and the bleaching process was carefully monitored using microscopes until most of the worm bodies were dissolved. Although the duration of bleaching was dependent on the species and the number of the worms, it usually took 5–15 min until we ceased the bleaching process. Before removing the bleaching solution by centrifuge, since eggs of some species (especially *P. sambesii*) did not sink to the bleaching solution, we added the same amount of M9 buffer. Subsequently, the eggs were washed with M9 buffer and centrifuged (12,000 g; for 30–60 sec) two-three times and further concentrated to 100–200 µl. Finally, the obtained solutions with eggs were placed onto the plain agar plates, which are optimized to each species. The solution was not sprayed to the plate, but spotted at a central part; the plate was also slowly dried in the clean bench. Within two to three days, the L1 worms hatched and moved from the area where the buffer solution was originally placed. We collected the L1 worms by removing the spotted area and washing with M9 buffer solution. We found that many *D. pachys* and *P. sambesii* worms did not survive in the M9 buffer without streaking onto plain agar plates.

#### Observation of molting process and staging

We began culturing of collected synchronous L1 worms by transferring them to an agar plate with an OP50 or HT115 lawn. To observe molting stages and perform pharmacological analysis, the worms were cultured at 20°C. We anesthetized the worms using sodium azide and observed the worms on the agar plate for differential interference constrast (DIC) imaging. We characterized the larval stages of *D. pachys* and *P. sambesii* based on observations of gonadal development, with reference to the previous reports on the same genus species (Tahseen, et al. 1991; TAHsEEN, et al. 1992; Hubbard and Greenstein 2000; Schindler and Sherwood 2013). As shown in Fig. S7A–C, gonadal primordia can be observed even in L1 worms. The primordia enlarge in the L2 stage and elongate further in the L3 stage. During the L4 stage, the primordiaelongate significantly, and the developed vulva can also be clearly observed. Adult worms can easily be distinguished by observing the developed gonads containing eggs. Male worms are very rare in all examined species, and we focused on hermaphroditic adults of *C. elegans* and parthenogenetic adults of *D. pachys* and *P. sambesii*. Nevertheless, since it is difficult to distinguish the sex of the worms during early larval stages (e.g., L1 and L2), L1 worms were used for the subsequent analysis without classifying their sex. We confirmed that the worms developed “female” morphology.

#### Pharmacological analysis

We commercially purchased cucurbitacin B (CucB, CAS number: 6199-67-3) from TargetMol (Wellesley Hills, MA, USA). We prepared a 50 mM stock solution of CucB in DMSO, and the experiments were performed in four-well plates. We added 500 µl of warm agar solution into a well after autoclaving, and 5 µl of CucB or DMSO was also added and mixed well before the agar hardened. In our preliminary tests, 50 µM is the highest concentration which can be solved in the agar plate. Subsequently, the bacterial food (10 µl in LB liquid culture) was added on the surface of the agar. Note that we used the optimized agar plates for culturing each species. Finally, synchronous L1 worms were transferred to the agar plates, and the plates were placed in the dark for two or three days at 20°C. For the experiments targeting the L3–L4 molting of *P. sambesii*, we used worms cultured for four days, as shown above. The periods of drug treatment are shown in Figs 3C and S7A. The morphology of the treated worms was observed as mentioned above. The larval and adult stages of the worms were also determined using the previously mentioned criteria. We judged the separation of old cuticles from the body as the apolysis process. We counted the number of worms that developed into landmark stages (shown in Fig. 3C) and the number of worms in the process of apolysis. All processes were repeated three times (three replicates), and the raw results are shown in Table S2. We performed a statistical analysis (Student’s t-test) using R (version 4.3.3).

### 10. Data visualization

We used R (version: 4.3.3) (Team 2024) and its packages such as Tydyverse (version: 2.0.0) (Wickham, et al. 2019), ggtree (version: 3.10.1) (Xu, et al. 2022), ggstance (version: 0.3.7, https://github.com/lionel-/ggstance), ggvenn (version: 0.1.10, https://github.com/yanlinlin82/ggvenn), and ape (version: 5.8, https://cran.r-project.org/web/packages/ape/index.html) to analyze and visualize the results. Images for Fig. S6 are from PHYLOPIC (https://www.phylopic.org/).

### 11. Data availability

Scripts and datasets (e.g. assembly sequences, phylogenetic trees, data accession numbers) are available in the supplementary dataset.

## Acknowledgments

We thank all current and former member of the Hejnol laboratory. In particular, we thank Nina Levin for proofreading of the manuscript. This study was supported by JSPS Oversea Fellowship (Japan Society for the Promotion of Science, to S.Y.), IMPULSE project (Friedrich Schiller University Jena, to S.Y.), Deutsche Forschungsgemeinschaft (project number 519107654, to A.H.), and the HFSP grant (RGP0041/2022, to A.H.).

## Supplementary information

Supplementary dataset

Supplementary figures (S1–S15, see the attached PDF file)

Supplementary tables (S1–S5, see the attached excel file)

## Author Contributions

Conceptualization, S.Y..; methodology, S.Y. and L.-M.B.; software, S.Y. and L.-M.B.; formal analysis, S.Y.; investigation, S.Y and L.-M.B..; writing—original draft, S.Y.; writing—review & editing, S.Y., L.-M.B., and A.H.; visualization, S.Y.; funding acquisition, S.Y. and A.H.

