## Supplementary figures for "Multiple losses of ecdysone receptor genes in nematodes: an alternative evolutionary scenario of molting regulation"

Figure S1. Validation of the multiple losses of *ecr* and *usp* in the tree reconstructed by Ahmed et al., 2022

Figure S2. Reannotation of the *ecr* of *Strongyloides* species

Figure S3. Genome assembly metrics and loss of *ecr* and *usp* (logarithmic scale)

Figure S4. Genome assembly metrics and loss of *ecr* and *usp* (linear scale)

Figure S5. BUSCO scores and loss of *ecr* and *usp*

Figure S6. Gene repertory (ecr, *usp*, and *hr3)* and molting regulation in  ecdysozoans

Figure S7. Staging of larval and adult development in *C. elegans*, *D. pachys*, and *P. sambesii*

Figure S8. Effects of CucB treatment on the third molting in *P. sambesii*

Figure S9. Comparison of original and corrected expression pattern of *ecr* and *hr3* in *P. pacificus*

Figure S10. RNA-seq clustering of *A. ceylanisum*, *G. pallida*, *B. Malayi*, and *T. suis*

Figure S11. Temporal expression of *grh-1* and *blmp-1* in six nematode species

Figure S12. Venn diagram of orthogroups of nuclear receptors in the representative nematode groups

Figure S13. Subsequent analysis of pairwise TM-align analysis

Figure S14. Phylogeny of expanded nematode HNF4 genes

Figure S15. Screening of *C. elegans* HNF4 proteins which can interact with *D. melanogaster* ECR

Figure S16. Additional predictions of heterodimer structure with ECR

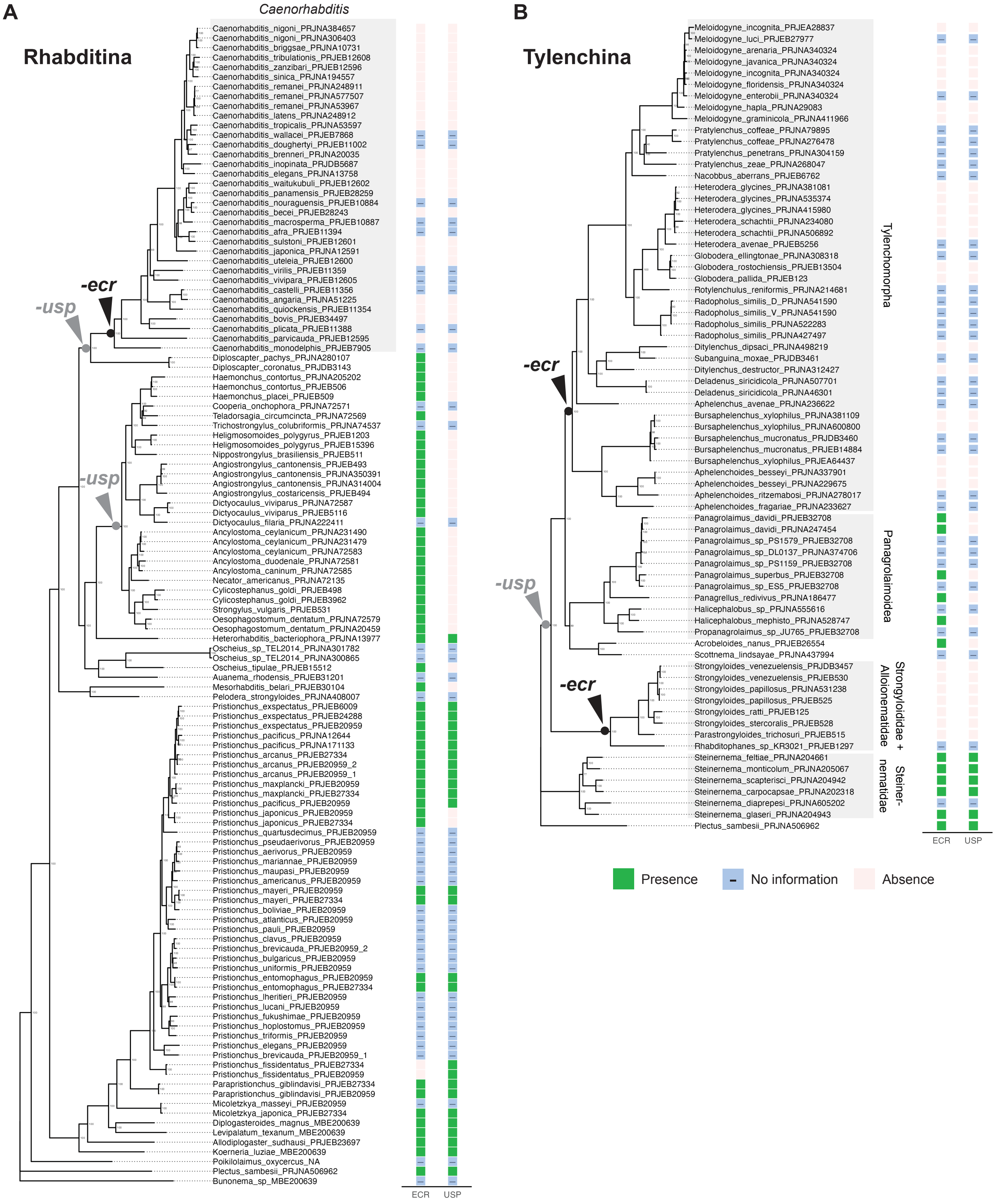


**Figure S1. Validation of the multiple losses of *ecr* and *usp* in the tree reconstructed by Ahmed et al., 2022**

**A**, **B**: the presence and absence of *ecr* and *usp* genes were mapped onto trees (**A**: Rhabditina and **B**: Tylenchina), which were reconstructed by Ahmed et al., 2022. Species which was not examined in this study are indicated with “No information”.

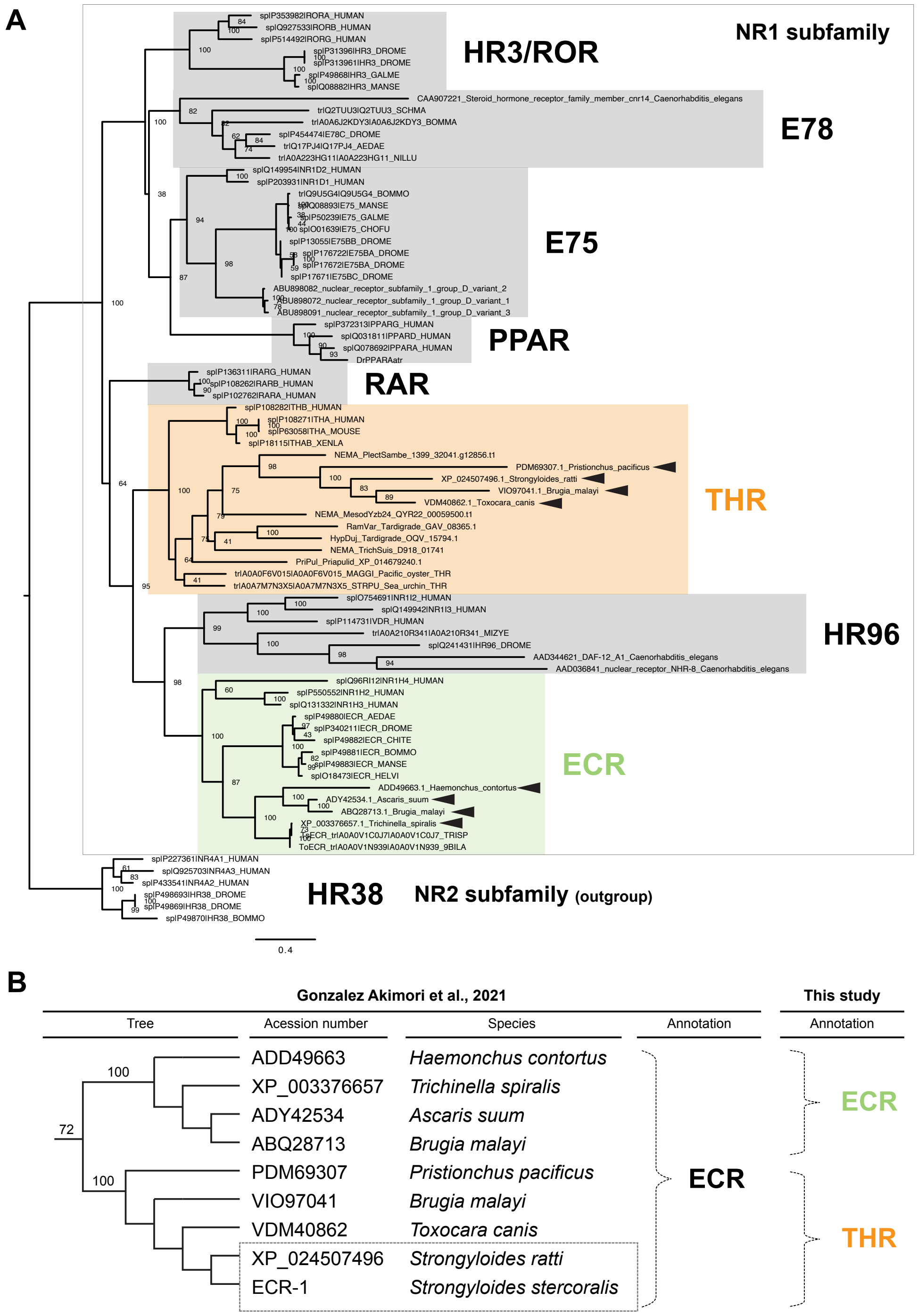


**Figure S2. Reannotation of the *ecr* of *Strongyloides* species**

**A**: Maximum likelihood tree of NR1 subtype nuclear receptors to reannotate the target genes which were previously identified in Fig. 3A of Gonzalez Akimori et al., 2021. The accession numbers of the target genes are shown in the tree and in panel **B** (arrowheads in the tree). The sequence of *Strongyloides stercoralis* could not be determined from the original paper. Shaded boxes indicate gene annotation (green: ECR, orange: THR [thyroid hormone receptor], and gray: other NR1 genes). THRs have also been identified in priapulids (*Priapulus caudutus*), tardigrades (*Hypsibius exemplaris* and *Ramazzottius varieornatus*), and other nematodes (*Plectus sambesii, Trichuris suis*, and *Mesodorylaimus sp. YZB2_4)*. **B**: Original annotation by Gonzalez Akimori et al., 2021 and correction by this study. The tree/bootstrap values shown on the left represent the original tree (without scaling: Gonzalez Akimori et al., 2021), and the correction of the annotation was summarized on the right.

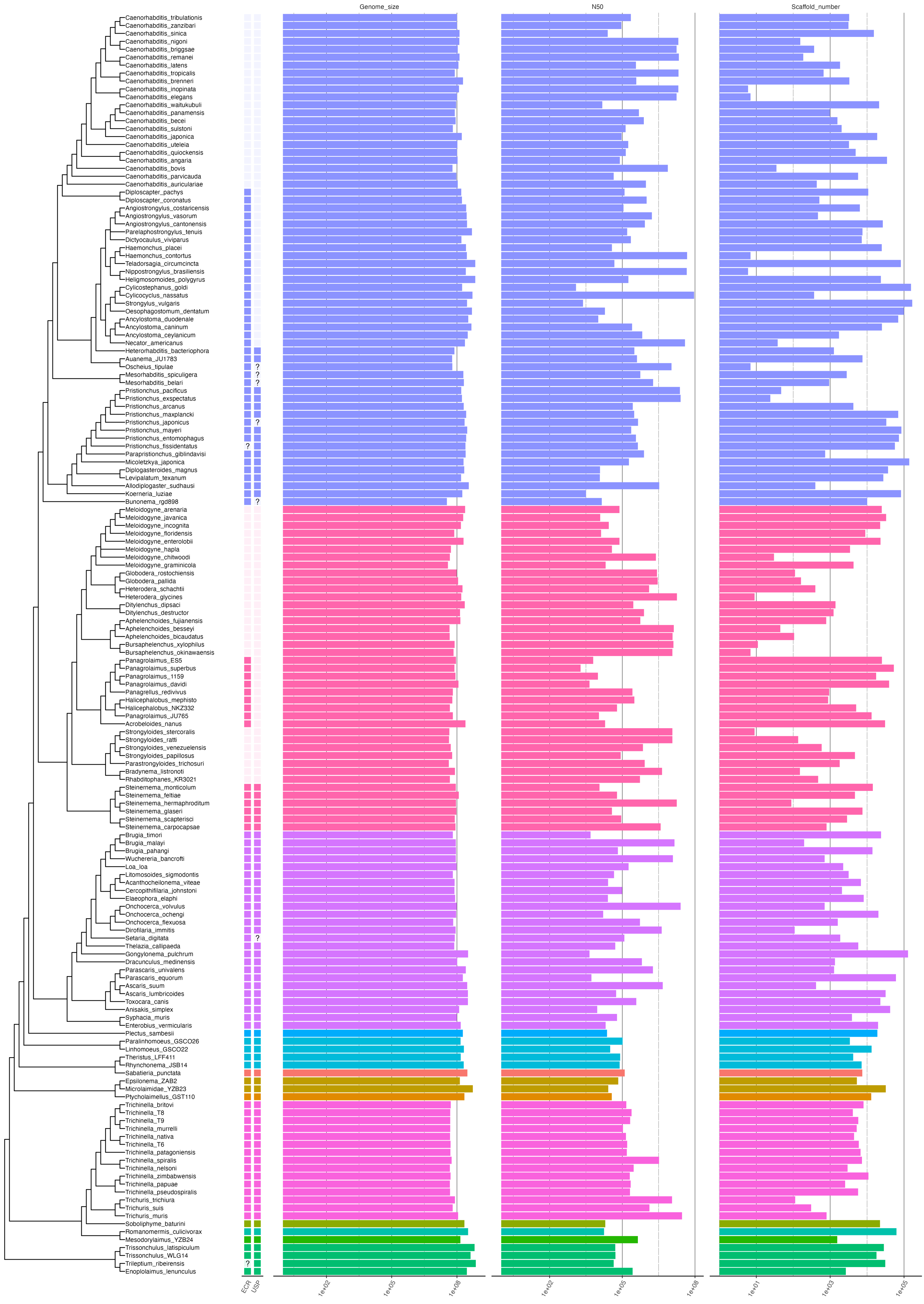
Genome_size
N50
Scaffold_number
Caenorhabditis_tribulationis
Caenorhabditis_zanzibari
Caenorhabditis_sinica
Caenorhabditis_nigoni
Caenorhabditis_briggsae
Caenorhabditis_remanei
Caenorhabditis_latens
Caenorhabditis_tropicalis
Caenorhabditis_brenneri
Caenorhabditis_inopinata
Caenorhabditis_elegans
Caenorhabditis_waitukubuli
Caenorhabditis_panamensis
Caenorhabditis_becei
Caenorhabditis_sulstoni
Caenorhabditis_japonica
Caenorhabditis_uteleia
Caenorhabditis_quiockensis
Caenorhabditis_angaria
Caenorhabditis_bovis
Caenorhabditis_parvicauda
Caenorhabditis_auriculariae
Diploscapter_pachys
Diploscapter_coronatus
Angiostrongylus_costaricensis
Angiostrongylus_vasorum
Angiostrongylus_cantonensis
Parelaphostrongylus_tenuis
Dictyocaulus_viviparus
Haemonchus_placei
Haemonchus_contortus
Teladorsagia_circumcincta
Nippostrongylus_brasiliensis
Heligmosomoides_polygyrus
Cylicostephanus_goldi
Cylicocyclus_nassatus
Strongylus_vulgaris
Oesophagostomum_dentatum
Ancylostoma_duodenale
Ancylostoma_caninum
Ancylostoma_ceylanicum
Necator_americanus
Heterorhabditis_bacteriophora
Auanema_JU1783
Oscheius_tipulae
Mesorhabditis_spiculigera
Mesorhabditis_belari
Pristionchus_pacificus
Pristionchus_exspectatus
Pristionchus_arcanus
Pristionchus_maxplancki
Pristionchus_japonicus
Pristionchus_mayeri
Pristionchus_entomophagus
Pristionchus_fissidentatus
Parapristionchus_giblindavisi
Micoletzkya_japonica
Diplogasteroides_magnus
Levipalatum_texanum
Allodiplogaster_sudhausi
Koerneria_luziae
Bunonema_rgd898
Meloidogyne_arenaria
Meloidogyne_javanica
Meloidogyne_incognita
Meloidogyne_floridensis
Meloidogyne_enterolobii
Meloidogyne_hapla
Meloidogyne_chitwoodi
Meloidogyne_graminicola
Globodera_rostochiensis
Globodera_pallida
Heterodera_schachtii
Heterodera_glycines
Ditylenchus_dipsaci
Ditylenchus_destructor
Aphelenchoides_fujianensis
Aphelenchoides_besseyi
Aphelenchoides_bicaudatus
Bursaphelenchus_xylophilus
Bursaphelenchus_okinawaensis
Panagrolaimus_ES5
Panagrolaimus_superbus
Panagrolaimus_1159
Panagrolaimus_davidi
Panagrellus_redivivus
Halicephalobus_mephisto
Halicephalobus_NKZ332
Panagrolaimus_JU765
Acrobeloides_nanus
Strongyloides_stercoralis
Strongyloides_ratti
Strongyloides_venezuelensis
Strongyloides_papillosus
Parastrongyloides_trichosuri
Bradynema_listronoti
Rhabditophanes_KR3021
Steinernema_monticolum
Steinernema_feltiae
Steinernema_hermaphroditum
Steinernema_glaseri
Steinernema_scapterisci
Steinernema_carpocapsae
Brugia_timori
Brugia_malayi
Brugia_pahangi
Wuchereria_bancrofti
Loa_loa
Litomosoides_sigmodontis
Acanthocheilonema_viteae
Cercopithifilaria_johnstoni
Elaeophora_elaphi
Onchocerca_volvulus
Onchocerca_ochengi
Onchocerca_flexuosa
Dirofilaria_immitis
Setaria_digitata
Thelazia_callipaeda
Gongylonema_pulchrum
Dracunculus_medinensis
Parascaris_univalens
Parascaris_equorum
Ascaris_suum
Ascaris_lumbricoides
Toxocara_canis
Anisakis_simplex
Syphacia_muris
Enterobius_vermicularis
Plectus_sambesii
Paralinhomoeus_GSCO26
Linhomoeus_GSCO22
Theristus_LFF411
Rhynchonema_JSB14
Sabatieria_punctata
Epsilonema_ZAB2
Microlaimidae_YZB23
Ptycholaimellus_GST110
Trichinella_britovi
Trichinella_T8
Trichinella_T9
Trichinella_murrelli
Trichinella_nativa
Trichinella_T6
Trichinella_patagoniensis
Trichinella_spiralis
Trichinella_nelsoni
Trichinella_zimbabwensis
Trichinella_papuae
Trichinella_pseudospiralis
Trichuris_trichiura
Trichuris_suis
Trichuris_muris
Soboliphyme_baturini
Romanomermis_culicivorax
Mesodorylaimus_YZB24
Trissonchulus_latispiculum
Trissonchulus_WLG14
Trileptium_ribeirensis
Enoplolaimus_lenunculus
ECR
USP
1e+02
1e+05
1e+08
1e+02
1e+05
1e+08
1e+01
1e+03
1e+05

**Figure S3. Genome assembly metrics and loss of *ecr* and *usp* (logarithmic scale)**

The left tree shows the phylogeny of the target species and the presence or absence of *ecr* and *usp* (see also Fig. 2 in the main text). The right bar graphs indicate the genome size, N50, and scaffold number in each species. The x-axis is on a log scale.

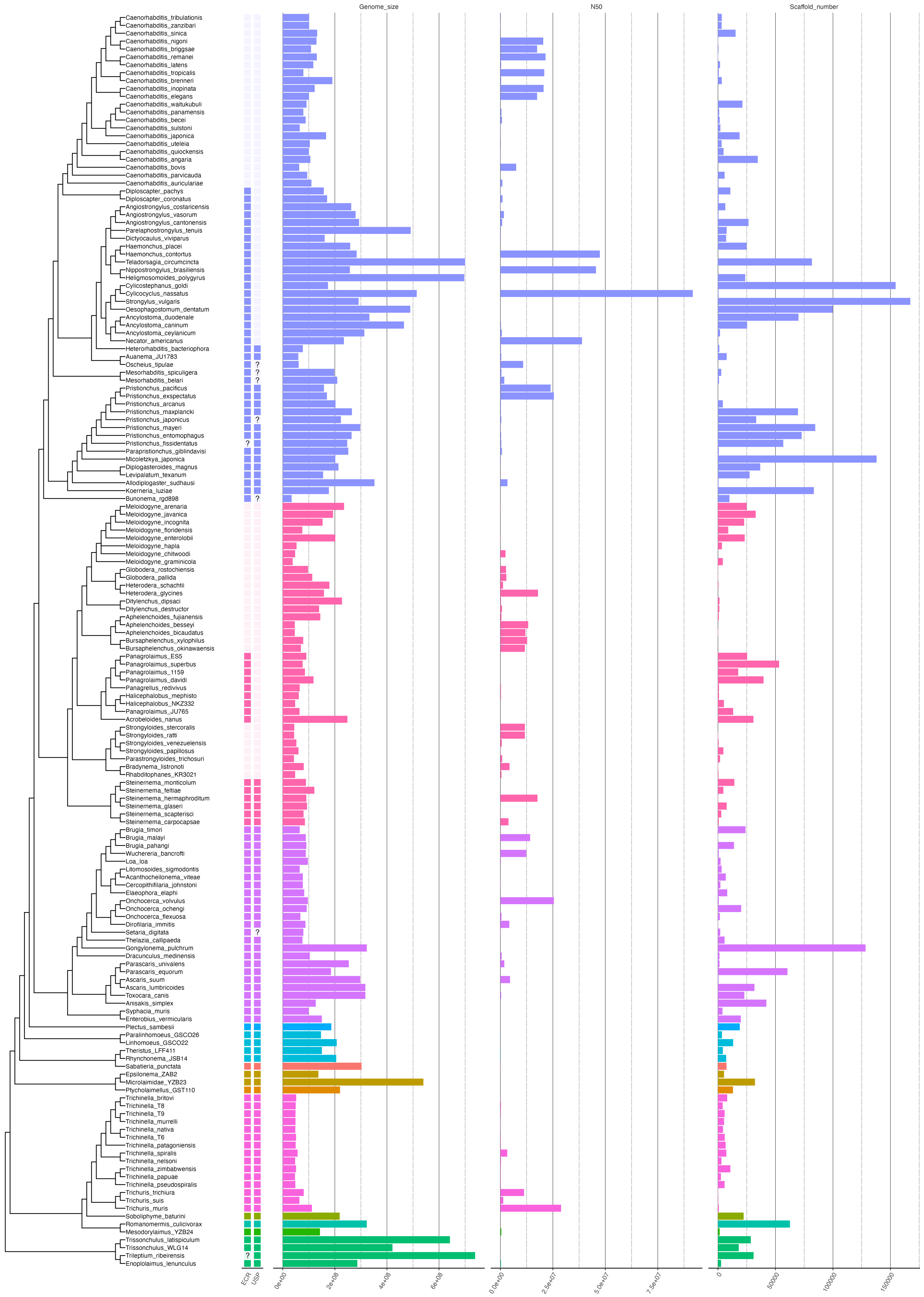

Genome_size
N50
Scaffold_number
Caenorhabditis_tribulationis
Caenorhabditis_zanzibari
Caenorhabditis_sinica
Caenorhabditis_nigoni
Caenorhabditis_briggsae
Caenorhabditis_remanei
Caenorhabditis_latens
Caenorhabditis_tropicalis
Caenorhabditis_brenneri
Caenorhabditis_inopinata
Caenorhabditis_elegans
Caenorhabditis_waitukubuli
Caenorhabditis_panamensis
Caenorhabditis_becei
Caenorhabditis_sulstoni
Caenorhabditis_japonica
Caenorhabditis_uteleia
Caenorhabditis_quiockensis
Caenorhabditis_angaria
Caenorhabditis_bovis
Caenorhabditis_parvicauda
Caenorhabditis_auriculariae
Diploscapter_pachys
Diploscapter_coronatus
Angiostrongylus_costaricensis
Angiostrongylus_vasorum
Angiostrongylus_cantonensis
Parelaphostrongylus_tenuis
Dictyocaulus_viviparus
Haemonchus_placei
Haemonchus_contortus
Teladorsagia_circumcincta
Nippostrongylus_brasiliensis
Heligmosomoides_polygyrus
Cylicostephanus_goldi
Cylicocyclus_nassatus
Strongylus_vulgaris
Oesophagostomum_dentatum
Ancylostoma_duodenale
Ancylostoma_caninum
Ancylostoma_ceylanicum
Necator_americanus
Heterorhabditis_bacteriophora
Auanema_JU1783
Oscheius_tipulae
Mesorhabditis_spiculigera
Mesorhabditis_belari
Pristionchus_pacificus
Pristionchus_exspectatus
Pristionchus_arcanus
Pristionchus_maxplancki
Pristionchus_japonicus
Pristionchus_mayeri
Pristionchus_entomophagus
Pristionchus_fissidentatus
Parapristionchus_giblindavisi
Micoletzkya_japonica
Diplogasteroides_magnus
Levipalatum_texanum
Allodiplogaster_sudhausi
Koerneria_luziae
Bunonema_rgd898
Meloidogyne_arenaria
Meloidogyne_javanica
Meloidogyne_incognita
Meloidogyne_floridensis
Meloidogyne_enterolobii
Meloidogyne_hapla
Meloidogyne_chitwoodi
Meloidogyne_graminicola
Globodera_rostochiensis
Globodera_pallida
Heterodera_schachtii
Heterodera_glycines
Ditylenchus_dipsaci
Ditylenchus_destructor
Aphelenchoides_fujianensis
Aphelenchoides_besseyi
Aphelenchoides_bicaudatus
Bursaphelenchus_xylophilus
Bursaphelenchus_okinawaensis
Panagrolaimus_ES5
Panagrolaimus_superbus
Panagrolaimus_1159
Panagrolaimus_davidi
Panagrellus_redivivus
Halicephalobus_mephisto
Halicephalobus_NKZ332
Panagrolaimus_JU765
Acrobeloides_nanus
Strongyloides_stercoralis
Strongyloides_ratti
Strongyloides_venezuelensis
Strongyloides_papillosus
Parastrongyloides_trichosuri
Bradynema_listronoti
Rhabditophanes_KR3021
Steinernema_monticolum
Steinernema_feltiae
Steinernema_hermaphroditum
Steinernema_glaseri
Steinernema_scapterisci
Steinernema_carpocapsae
Brugia_timori
Brugia_malayi
Brugia_pahangi
Wuchereria_bancrofti
Loa_loa
Litomosoides_sigmodontis
Acanthocheilonema_viteae
Cercopithifilaria_johnstoni
Elaeophora_elaphi
Onchocerca_volvulus
Onchocerca_ochengi
Onchocerca_flexuosa
Dirofilaria_immitis
Setaria_digitata
Thelazia_callipaeda
Gongylonema_pulchrum
Dracunculus_medinensis
Parascaris_univalens
Parascaris_equorum
Ascaris_suum
Ascaris_lumbricoides
Toxocara_canis
Anisakis_simplex
Syphacia_muris
Enterobius_vermicularis
Plectus_sambesii
Paralinhomoeus_GSCO26
Linhomoeus_GSCO22
Theristus_LFF411
Rhynchonema_JSB14
Sabatieria_punctata
Epsilonema_ZAB2
Microlaimidae_YZB23
Ptycholaimellus_GST110
Trichinella_britovi
Trichinella_T8
Trichinella_T9
Trichinella_murrelli
Trichinella_nativa
Trichinella_T6
Trichinella_patagoniensis
Trichinella_spiralis
Trichinella_nelsoni
Trichinella_zimbabwensis
Trichinella_papuae
Trichinella_pseudospiralis
Trichuris_trichiura
Trichuris_suis
Trichuris_muris
Soboliphyme_baturini
Romanomermis_culicivorax
Mesodorylaimus_YZB24
Trissonchulus_latispiculum
Trissonchulus_WLG14
Trileptium_ribeirensis
Enoplolaimus_lenunculus
ECR
USP
0e+00
2e+08
4e+08
6e+08
0.0e+00
2.5e+07
5.0e+07
7.5e+07
0
50000
100000
150000

**Figure S4. Genome assembly metrics and loss of *ecr* and *usp* (linear scale)**

The left tree shows the phylogeny of the target species and the presence or absence of *ecr* and *usp* (see also Fig. 2 in the main text). The right bar graphs indicate the genome size, N50, and scaffold number in each species. The x-axis is on a linear scale.

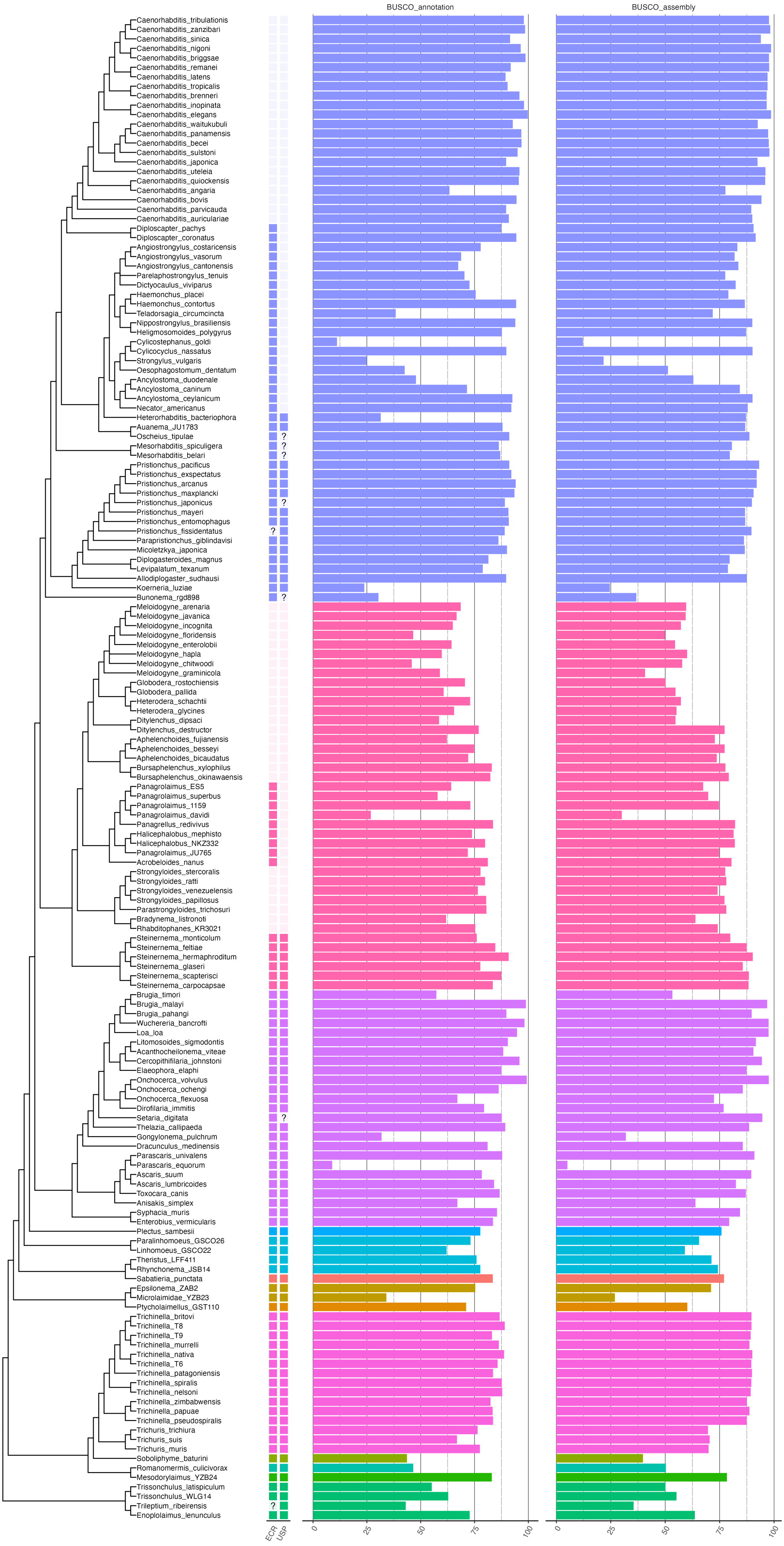

BUSCO_annotation
BUSCO_assembly
Caenorhabditis_tribulationis
Caenorhabditis_zanzibari
Caenorhabditis_sinica
Caenorhabditis_nigoni
Caenorhabditis_briggsae
Caenorhabditis_remanei
Caenorhabditis_latens
Caenorhabditis_tropicalis
Caenorhabditis_brenneri
Caenorhabditis_inopinata
Caenorhabditis_elegans
Caenorhabditis_waitukubuli
Caenorhabditis_panamensis
Caenorhabditis_becei
Caenorhabditis_sulstoni
Caenorhabditis_japonica
Caenorhabditis_uteleia
Caenorhabditis_quiockensis
Caenorhabditis_angaria
Caenorhabditis_bovis
Caenorhabditis_parvicauda
Caenorhabditis_auriculariae
Diploscapter_pachys
Diploscapter_coronatus
Angiostrongylus_costaricensis
Angiostrongylus_vasorum
Angiostrongylus_cantonensis
Parelaphostrongylus_tenuis
Dictyocaulus_viviparus
Haemonchus_placei
Haemonchus_contortus
Teladorsagia_circumcincta
Nippostrongylus_brasiliensis
Heligmosomoides_polygyrus
Cylicostephanus_goldi
Cylicocyclus_nassatus
Strongylus_vulgaris
Oesophagostomum_dentatum
Ancylostoma_duodenale
Ancylostoma_caninum
Ancylostoma_ceylanicum
Necator_americanus
Heterorhabditis_bacteriophora
Auanema_JU1783
Oscheius_tipulae
Mesorhabditis_spiculigera
Mesorhabditis_belari
Pristionchus_pacificus
Pristionchus_exspectatus
Pristionchus_arcanus
Pristionchus_maxplancki
Pristionchus_japonicus
Pristionchus_mayeri
Pristionchus_entomophagus
Pristionchus_fissidentatus
Parapristionchus_giblindavisi
Micoletzkya_japonica
Diplogasteroides_magnus
Levipalatum_texanum
Allodiplogaster_sudhausi
Koerneria_luziae
Bunonema_rgd898
Meloidogyne_arenaria
Meloidogyne_javanica
Meloidogyne_incognita
Meloidogyne_floridensis
Meloidogyne_enterolobii
Meloidogyne_hapla
Meloidogyne_chitwoodi
Meloidogyne_graminicola
Globodera_rostochiensis
Globodera_pallida
Heterodera_schachtii
Heterodera_glycines
Ditylenchus_dipsaci
Ditylenchus_destructor
Aphelenchoides_fujianensis
Aphelenchoides_besseyi
Aphelenchoides_bicaudatus
Bursaphelenchus_xylophilus
Bursaphelenchus_okinawaensis
Panagrolaimus_ES5
Panagrolaimus_superbus
Panagrolaimus_1159
Panagrolaimus_davidi
Panagrellus_redivivus
Halicephalobus_mephisto
Halicephalobus_NKZ332
Panagrolaimus_JU765
Acrobeloides_nanus
Strongyloides_stercoralis
Strongyloides_ratti
Strongyloides_venezuelensis
Strongyloides_papillosus
Parastrongyloides_trichosuri
Bradynema_listronoti
Rhabditophanes_KR3021
Steinernema_monticolum
Steinernema_feltiae
Steinernema_hermaphroditum
Steinernema_glaseri
Steinernema_scapterisci
Steinernema_carpocapsae
Brugia_timori
Brugia_malayi
Brugia_pahangi
Wuchereria_bancrofti
Loa_loa
Litomosoides_sigmodontis
Acanthocheilonema_viteae
Cercopithifilaria_johnstoni
Elaeophora_elaphi
Onchocerca_volvulus
Onchocerca_ochengi
Onchocerca_flexuosa
Dirofilaria_immitis
Setaria_digitata
Thelazia_callipaeda
Gongylonema_pulchrum
Dracunculus_medinensis
Parascaris_univalens
Parascaris_equorum
Ascaris_suum
Ascaris_lumbricoides
Toxocara_canis
Anisakis_simplex
Syphacia_muris
Enterobius_vermicularis
Plectus_sambesii
Paralinhomoeus_GSCO26
Linhomoeus_GSCO22
Theristus_LFF411
Rhynchonema_JSB14
Sabatieria_punctata
Epsilonema_ZAB2
Microlaimidae_YZB23
Ptycholaimellus_GST110
Trichinella_britovi
Trichinella_T8
Trichinella_T9
Trichinella_murrelli
Trichinella_nativa
Trichinella_T6
Trichinella_patagoniensis
Trichinella_spiralis
Trichinella_nelsoni
Trichinella_zimbabwensis
Trichinella_papuae
Trichinella_pseudospiralis
Trichuris_trichiura
Trichuris_suis
Trichuris_muris
Soboliphyme_baturini
Romanomermis_culicivorax
Mesodorylaimus_YZB24
Trissonchulus_latispiculum
Trissonchulus_WLG14
Trileptium_ribeirensis
Enoplolaimus_lenunculus
?
ECR
USP
0
25
50
75
100

**Figure S5. BUSCO scores and loss of *ecr* and *usp***

The left tree shows the phylogeny of the target species and the presence or absence of *ecr* and *usp* (see also Fig. 2 in the main text). The right bar graphs indicate the BUSCO scores (annotation and assembly).

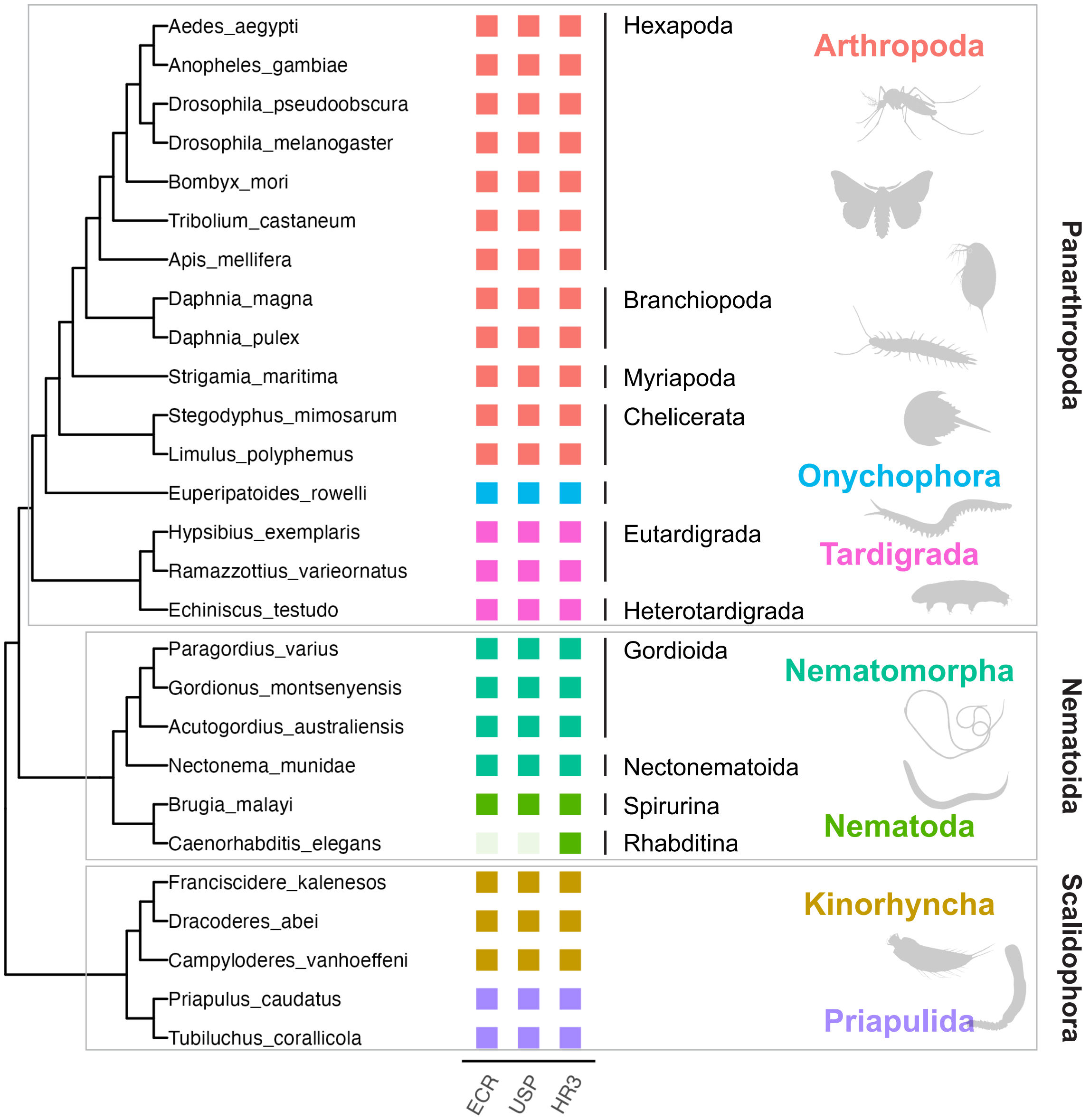


**Figure S6. Gene repertory (ecr, *usp*, and *hr3)* and molting regulation in ecdysozoans**

Phylogeny of Panarthropoda, Nematoida, and Scalidophora is based on Howard et al., 2022. The presence of *ecr*, *usp*, and *hr3* was indicated in the boxes next to each species name. Gene identification was referred to the previous annotation and this study (see Methods, Bonneton et al., 2018 and Schumann et al., 2018).

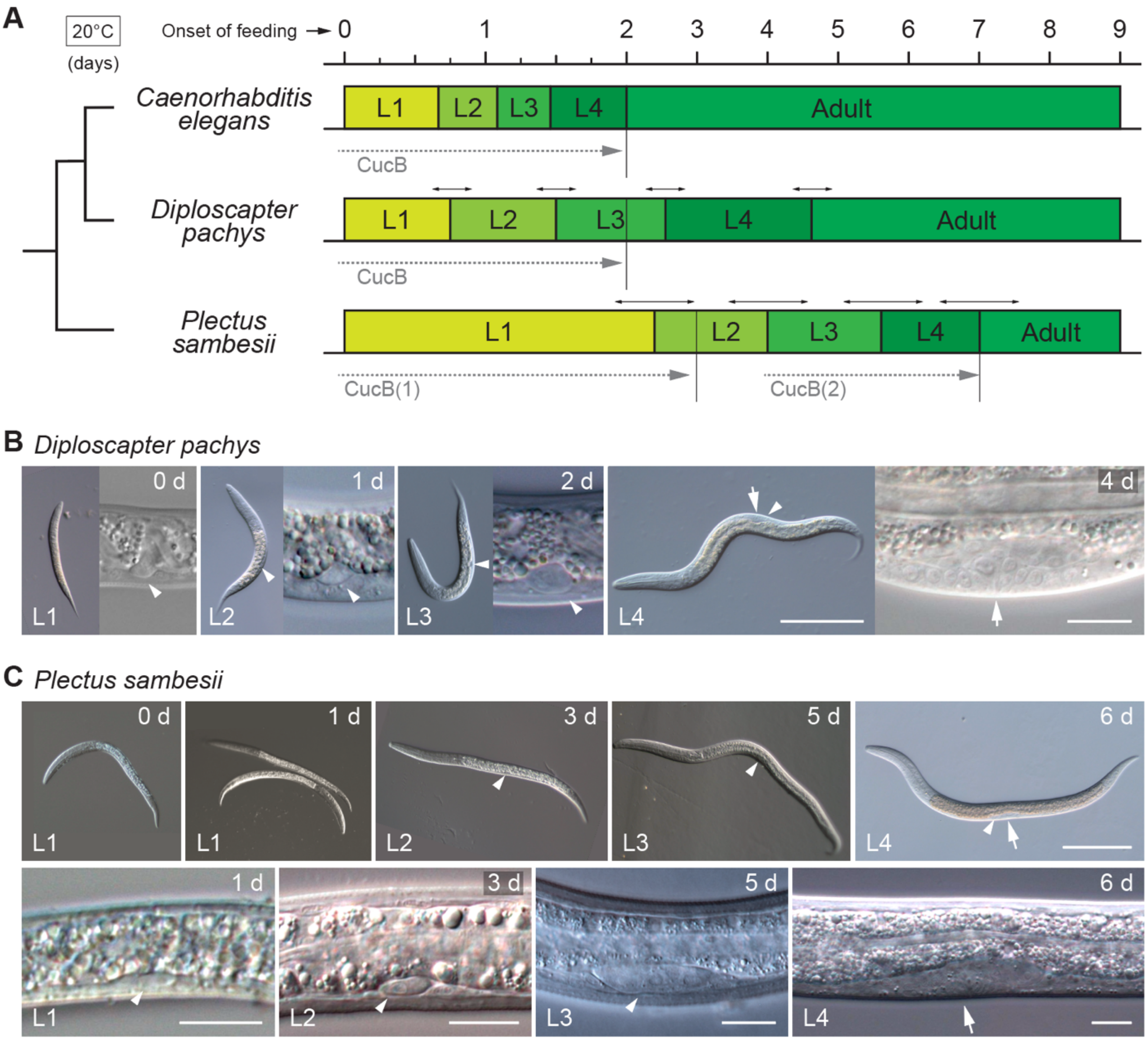


**Figure S7. Staging of larval and adult development in *C. elegans*, *D. pachys*, and *P. sambesii***

**A**: Duration of larval and adult stages after the commencement of feeding in tree species at 20°C. L1–L4 represent the first through fourth larval stages. The colors/boxes represent the duration of each stage (0–9 days after feeding). The double-headed arrows at the boundary of each stage indicate the possible stage shift observed in this study. Dotted arrows indicate the duration of CucB treatment, and vertical lines indicate the end of treatment. **B**, **C**: Representative worms and gonadal development at the L1–L4 stages (**B**: *D. pachys* [0, 1, 2, and 4 days after feeding] and **C**: *P. sambesii* [0, 1, 3, 5, and 6 days after feeding]). Arrowheads and arrows indicate the gonads and vulva, respectively. Scale bars: 100 µm for photos of whole worms and 10 µm for photos of gonads.

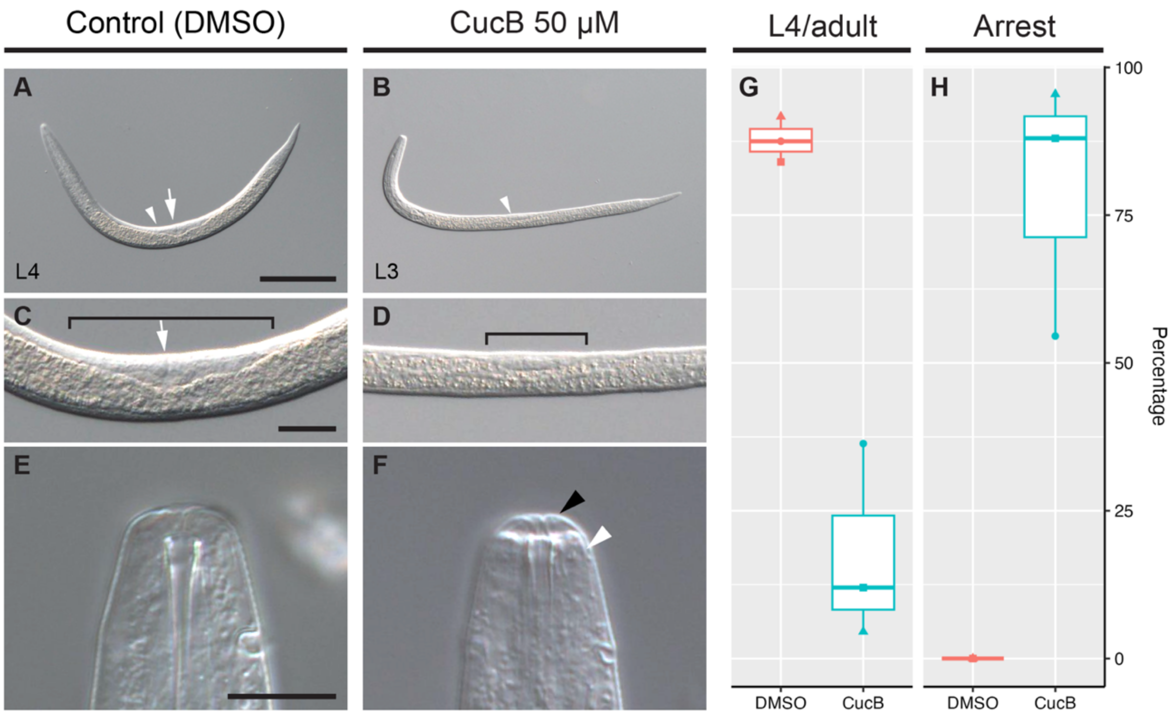


**Figure S8. Effects of CucB treatment on the third molting in *P. sambesii***

**A**–**F**: Representative worms after a three-day treatment with DMSO (**A**, **C**, and **E**) or 50 µM CucB (**B**, **D**, and **F**). **A**–**B**, **C**–**D**, and **E**–**F** shows the whole worms, magnified view of gonads, and magnified view of mouthparts, respectively (white arrow heads in **A**, **B**: gonads; white arrows in **A**, **C**: vulva; square brackets in **C**, **D**: gonads; and arrow heads in **F**: new [white] and old [black] cuticles). Scale bars: 100 µm for **A**, **B**; 25 µm for **C**, **D**; and 10 µm for **E**, **F**. **G**, **H**: The percentage of treated worms that developed to the L4 or adult stage (**G**) and those that arrested at the third molting stage (**H**).

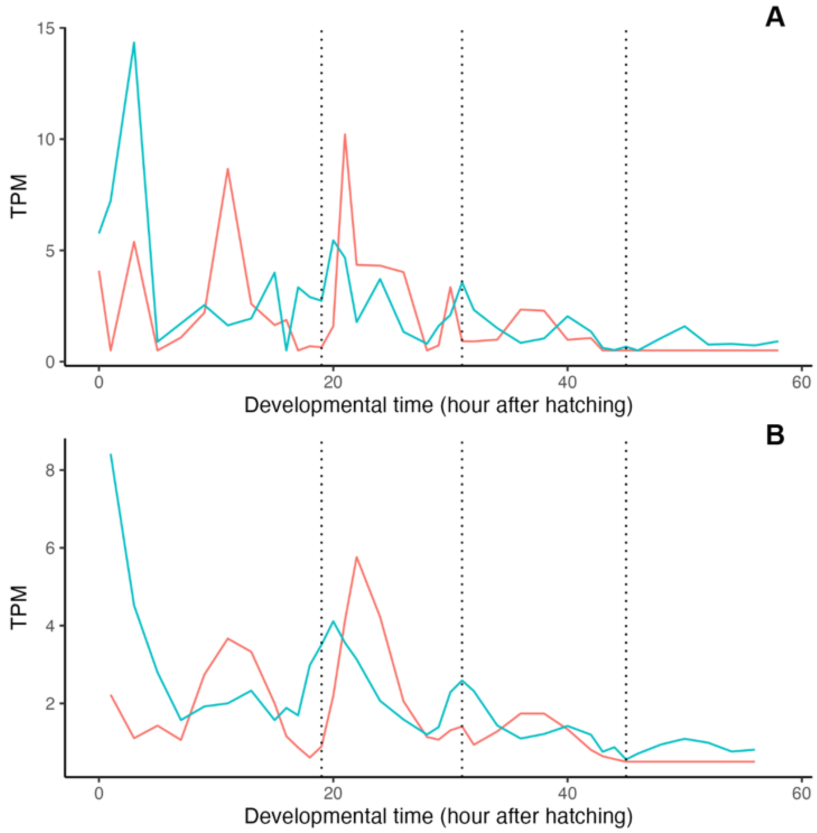


**Figure S9. Comparison of original and corrected expression pattern of *ecr* and *hr3* in *P. pacificus***

**A**: Original expression profiles of *ecr* (blue) and *hr3* (red) from Sun et al., 2021(2). **B**: corrected expression profiles of *ecr* and *hr3* by calculating the average expression levels (TPM) between the three time point windows from Sun et al., 2021 (see Methods).

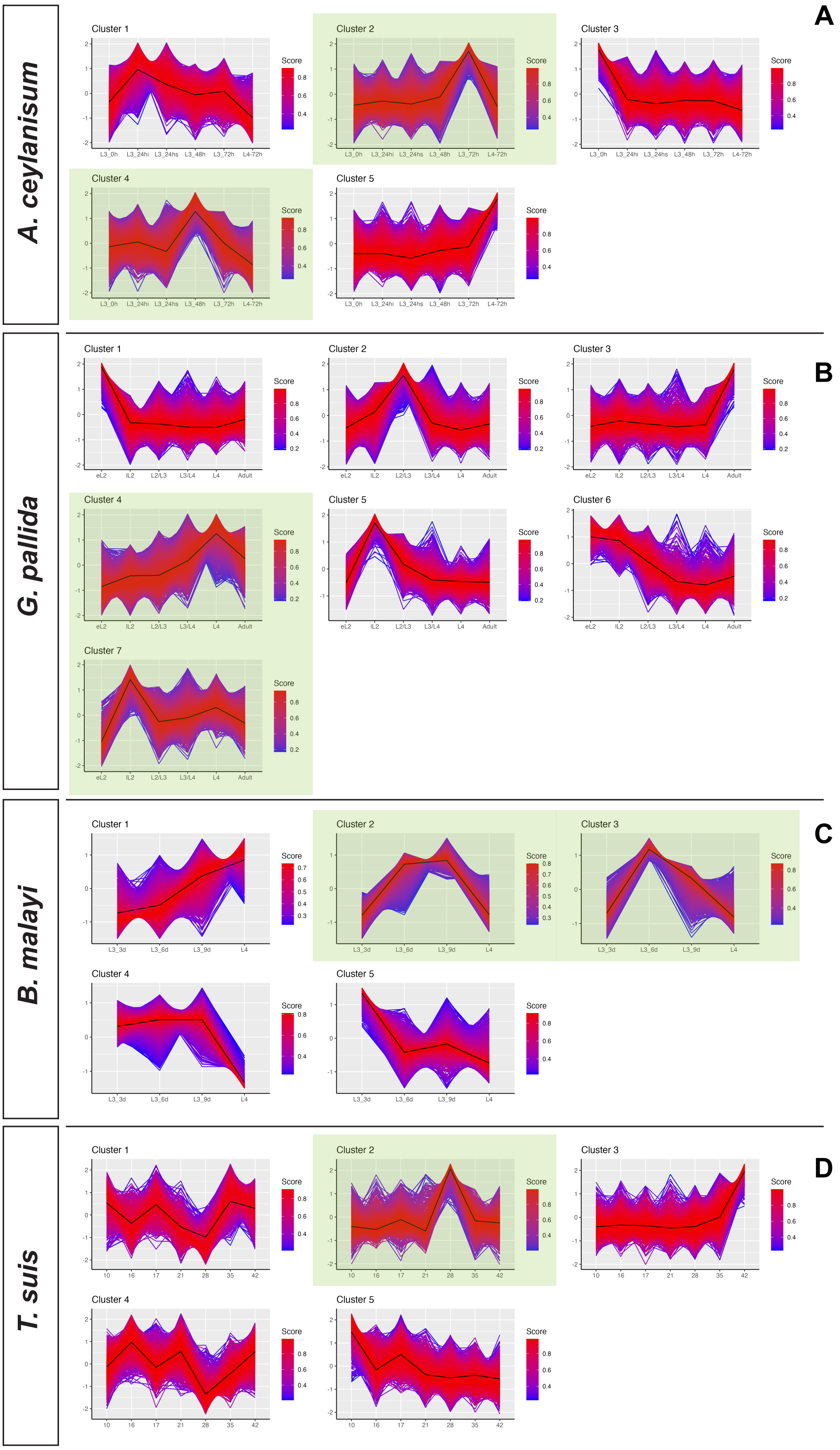


**Figure S10. RNA-seq clustering of *A. ceylanisum*, *G. pallida*, *B. Malayi*, and *T. suis***

**A–D**: Fuzzy c-mean clustering of each transcriptome data set (*A. ceylanisum*, *G. pallida*, *B. Malayi*, and *T. suis*, respectively). Green shades indicate the clusters showing correlated expression with molting cycle.

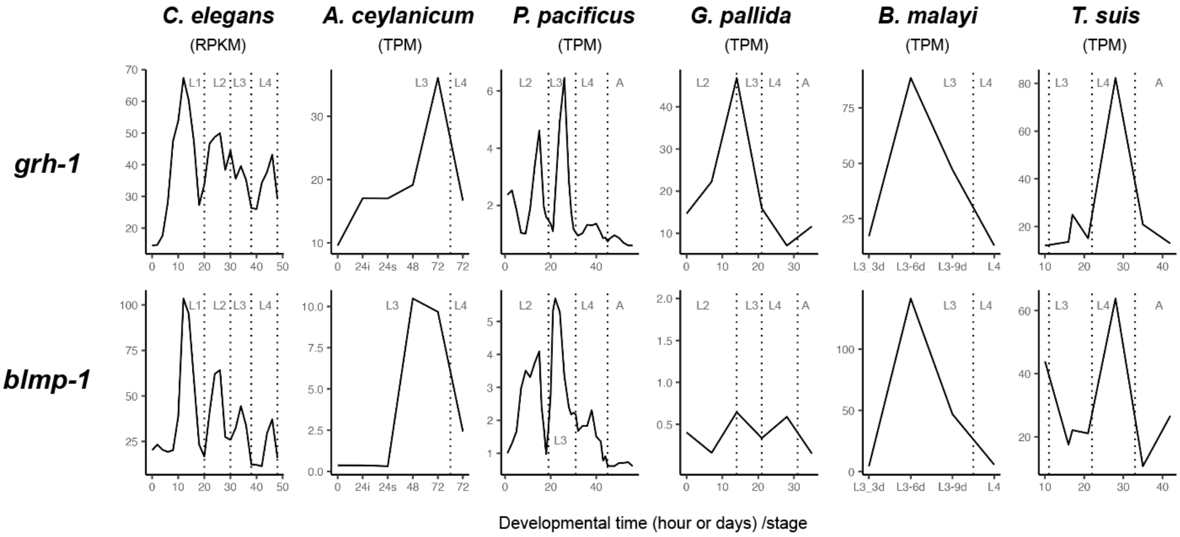


**Figure S11. Temporal expression of *grh-1* and *blmp-1* in six nematode species**

The x-axis indicates developmental time (hours for *C. elegans*, *A. ceylanicum*, [i: intestine, s: stomach] and *P. pacificus*; days for *G. pallida* and *T. suis*, and stages/days for *B. malayi*), and the y-axis shows expression levels (RPKM or TPM). Dotted lines show the molting timing (L1–L4: first to fourth larval stages, A: adult stage).

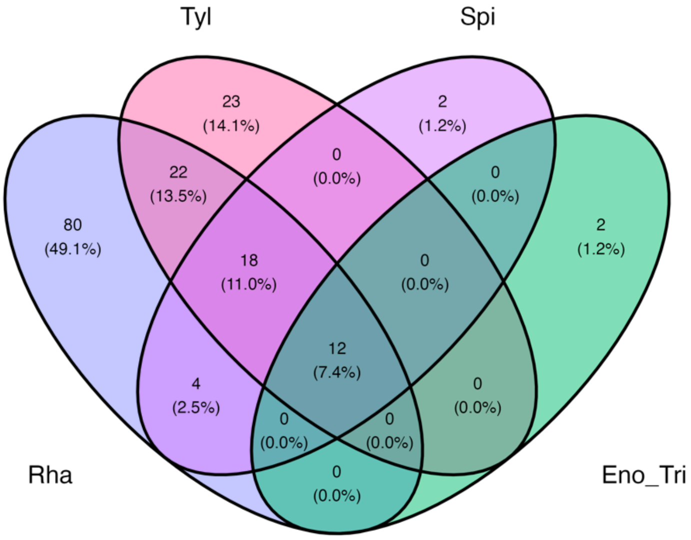


**Figure S12. Venn diagram of orthogroups of nuclear receptors in the representative nematode groups**

Rha: Rhabditina, Tyl: Tylenchina, Spi: Spirurina, Eno_Tri: Enoplida and Trichinellida.

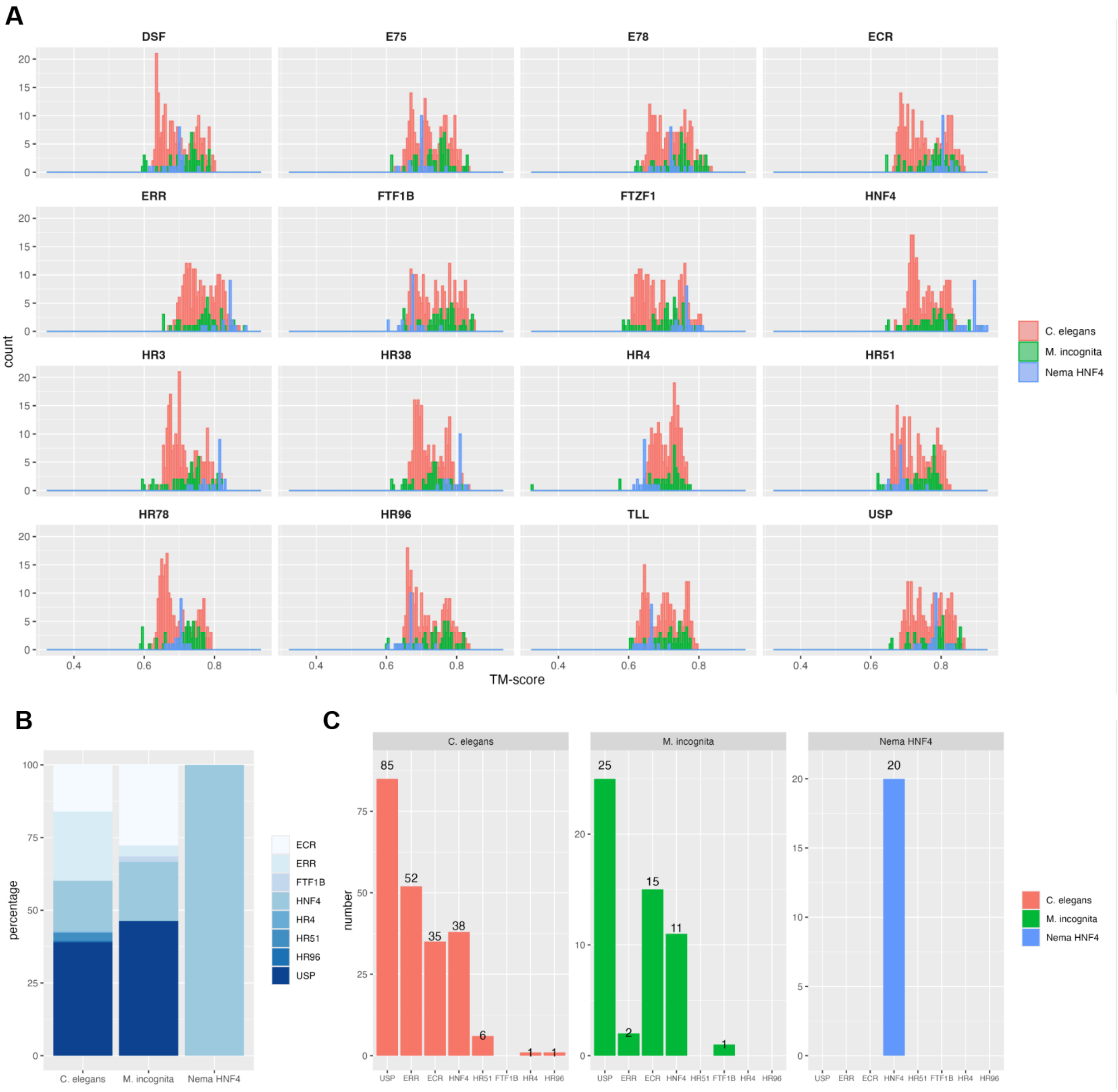


**Figure S13. Subsequent analysis of pairwise TM-align analysis**

**A**: Histograms showing the TM-score in pairwise comparison with each *D. melanogaster* NR. **B**, **C**: the percentage (**B**) and actual number (**C**) of reference NRs that have the highest TM-score in each pairwise alignment. For example, 100% (20 out of 20) of the HNF4 proteins from the early-branching nematode taxa ("Nema HNF4" in the panels) show the highest TM-score with the *D. melanogaster* HNF4 protein in the pairwise comparison. Conversely, 40–50% of the expanded HNF4 proteins from *C. elegans* (85 proteins) and *M. incognita* (25 proteins) have the highest TM-scores with USP.

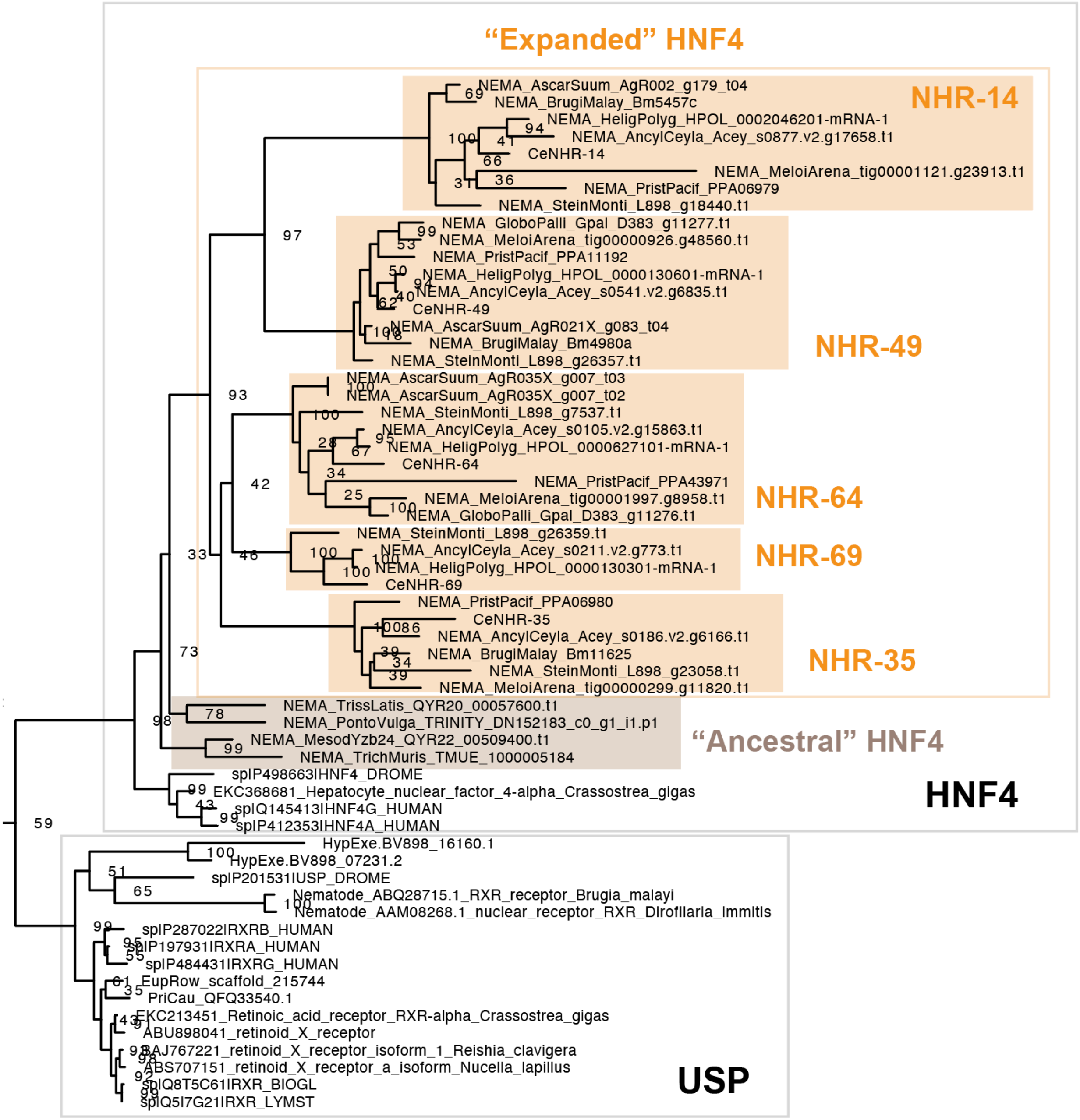


**Figure S14. Phylogeny of expanded nematode HNF4 genes**

A clade of HNF4 and USP was extracted from the tree to test the identity of the target proteins (nematode HNF4 and NHR-64, NHR-69, NHR-35, NHR-14, and NHR-49). The original tree is available in the supplementary dataset. The orange colors indicate the "expanded" HNF4s, which specifically evolved within the Spirurina/Tylenchina/Rhabditina clade of nematodes. "Ancestral" HNF4s represent proteins that evolved before NR expansion events (brown color). Gray squares indicate the clade of HNF4 and USP.

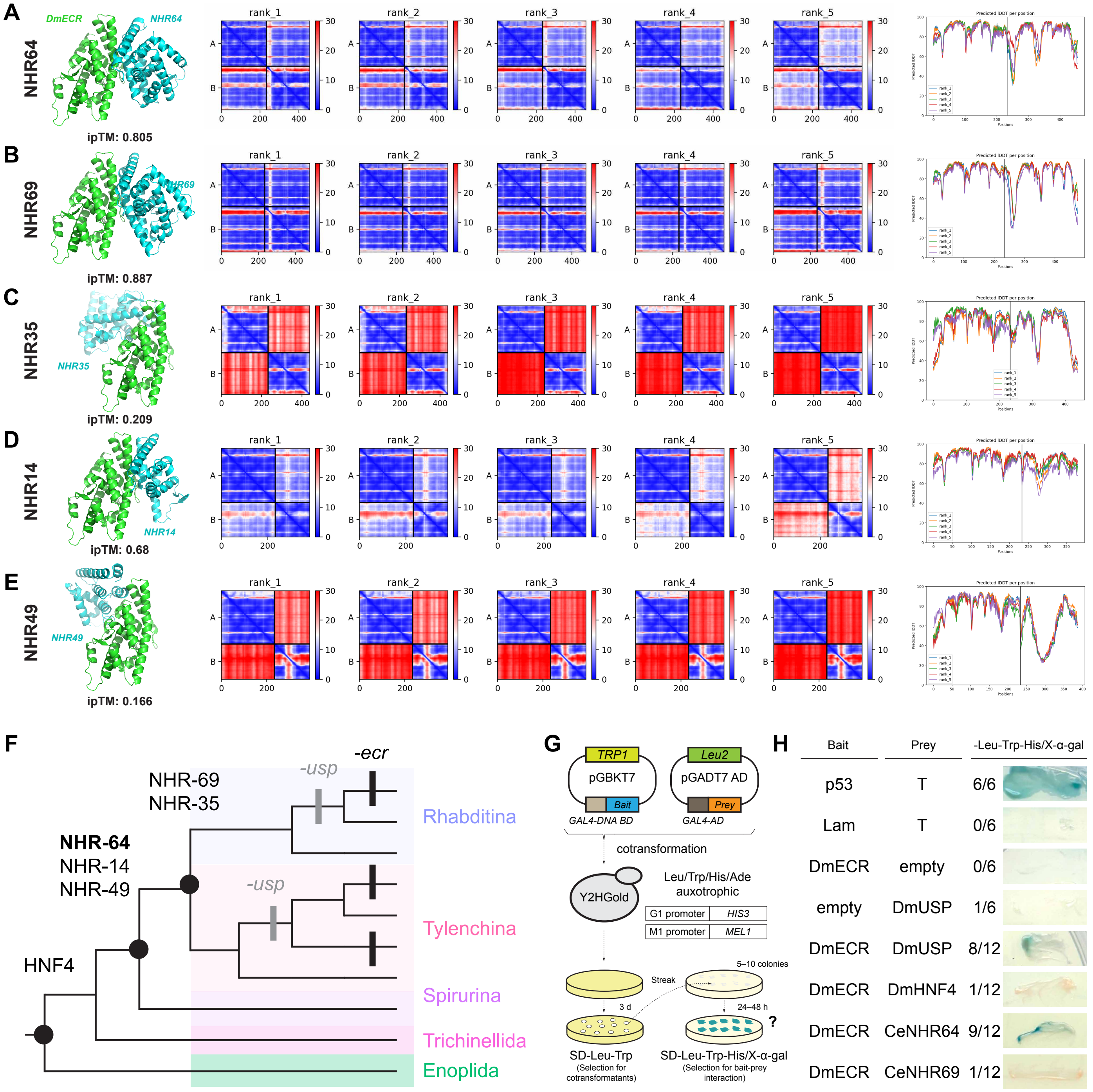
A
NHR64
DmECR
NHR64
ipTM: 0.805
B
NHR69
NHR69
ipTM: 0.887
C
NHR35
NHR35
ipTM: 0.209
D
NHR14
NHR14
ipTM: 0.68
E
NHR49
NHR49
ipTM: 0.166
rank_1
rank_2
rank_3
rank_4
rank_5
Predicted lDDT per position
F
-ecr
-usp
NHR-69
NHR-35
Rhabditina
NHR-64
NHR-14
NHR-49
-usp
Tylenchina
HNF4
Spirurina
Trichinellida
Enoplida
G
TRP1
pGBKT7
Bait
GAL4-DNA BD
Leu2
pGADT7 AD
Prey
GAL4-AD
cotransformation
Y2HGold
Leu/Trp/His/Ade auxotrophic
G1 promoter HIS3
M1 promoter MEL1
Streak
5–10 colonies
3 d
24–48 h
?
SD-Leu-Trp
(Selection for cotransformatants)
SD-Leu-Trp-His/X-α-gal
(Selection for bait-prey interaction)
H
Bait
Prey
-Leu-Trp-His/X-α-gal
p53 T 6/6
Lam T 0/6
DmECR empty 0/6
empty DmUSP 1/6
DmECR DmUSP 8/12
DmECR DmHNF4 1/12
DmECR CeNHR64 9/12
DmECR CeNHR69 1/12

**Figure S15. Screening of *C. elegans* HNF4 proteins which can interact with *D. melanogaster* ECR**

**A**–**E**: The left column shows the 3D models and ipTM values of the highest-scoring predicted heterodimer structures (*C. elegans* NHR-64, NHR-69, NHR-35, NHR-14, and NHR-49, respectively). All target proteins were investigated with ECR from *Drosophila melanogaster* (DmECR, shown in green). The center columns show the PAE heatmaps of the top five highest-scoring predicted models for each target protein. The right column shows predicted lDDT per position in all models. Raw results are also available in the supplementary dataset. **F**: Phylogeny of five nematode taxa and the origin of the target HNF4 proteins. **G**, **H**: a summary of the yeast two-hybrid assay. **G** shows an overview of the assay, including schematic diagrams of the plasmid maps (pGBKT7/Bait and pGADT7/Prey), the auxotroph and the reporter genes of the yeast strain used in this study (Y2HGold), and the selective condition (SD-Leu-Trp-His/X-alpha-Gal). **H**: the results of our screening. The target proteins for the bait and prey are shown in the two columns on the left. The number of positive/observed colonies is shown in the right column, along with a photo of representative colonies. P53-T and Lam-T were used as positive and negative controls, respectively. DmECR-empty and empty-DmUSP were tested for autoactivation. See the methods for more details.

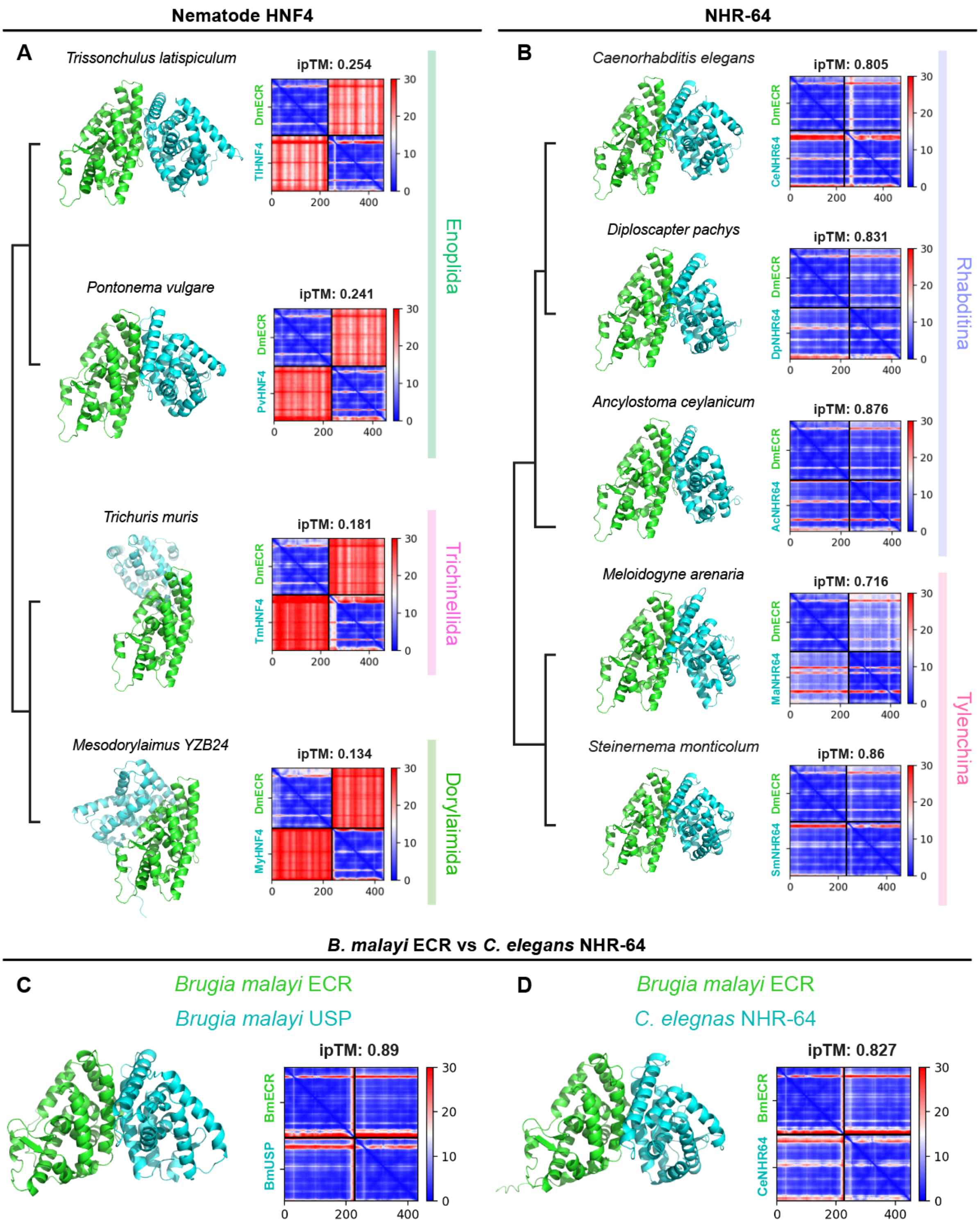


**Figure S16. Additional predictions of heterodimer structure with ECR**

**A**–**E**: The 3D models, ipTM values, and PAE heatmap depict the highest-scoring predicted heterodimer structure (**A**: nematode HNF4; **B**: NHR-64; and **C**, **D**: *Brugia malayi* ECR/USP and *Caenorhabditis elegans* NHR-64 comparison). The ipTM values and PAE heatmaps are shown to the right of the 3D models. The target proteins were investigated with ECR from *D. melanogaster* (DmECR) in panels **A** and **B**, and with ECR from *B. malayi* (BmECR) in panels **C**–**D** (**C**: *B. malayi* USP; **D**: *C. elegans* NHR-64).
