## Supplementary table for "Multiple losses of ecdysone receptor genes in nematodes: an alternative evolutionary scenario of molting regulation": Fig_S6.R

```
library(patchwork)
library(tidyverse)
library(ggtree)
library(ape)
library(ggstance)

tree <- read.tree('tree')

sum <- read.table("sum_table")
colnames(sum) <- c("sp","taxa","label")

sum <-  sum %>% as_tibble() %>% mutate(ECR = 1, USP = 1, HR3 =1)
sum <- sum %>% mutate(ECR = ifelse(label=="Caenorhabditis_elegans",0,ECR), USP = ifelse(label=="Caenorhabditis_elegans",0,USP))

p <- ggtree(tree)  + geom_tiplab(size=3, align=T, linesize=0.3) + ggplot2::xlim(0, 30)
tip_labels <- p$data[p$data$isTip, ]
tip_labels <- tip_labels[order(tip_labels$y), "label"]

gene <- sum %>% pivot_longer(-c(sp,label,taxa)) %>% mutate(alpha = ifelse(value>0,10,0), label = fct_relevel(label, tip_labels$label), 
gene = fct_relevel(name, c("ECR","USP"))) 


g_g <- gene %>% ggplot(aes(x=gene, y=label, color=taxa,alpha=alpha)) + geom_point(size=4,shape=15)  + theme_classic()+ theme(axis.text.y=element_blank(), 
axis.ticks=element_blank(), axis.title=element_blank(), legend.position="none", axis.line.y=element_blank(), axis.line.x=element_line(size=0.5), 
axis.text.x=element_text(angle=60, hjust=1) )

sum <- p + g_g + plot_layout(widths = c(1,0.3))
ggsave(plot=sum, file="ecdysozoan_ECR_USP_HR3.png", width=4, height=7)
```
