## Supplementary table for "Multiple losses of ecdysone receptor genes in nematodes: an alternative evolutionary scenario of molting regulation": results_summary_figure.R

```
library(patchwork)
library(tidyverse)
library(ggtree)
library(ape)
library(ggstance)

tree <- read.tree('nematode_phylogeny_newick')

sum <- read.table("results_summary")
colnames(sum) <- c("Style","Habitat","sp","label","id","clade","clade_n","taxa","file","nr","ecr","usp","hr3","lin42","e75","ftzf1","kin20")

sum <- sum %>% as_tibble() %>% select(-c(id,clade,clade_n,file))

p <- ggtree(tree)  + geom_tiplab(size=3, align=T, linesize=0.3) + ggplot2::xlim(0, 45)
tip_labels <- p$data[p$data$isTip, ]
tip_labels <- tip_labels[order(tip_labels$y), "label"]

life <-  sum %>% select(label,Style,Habitat) %>% pivot_longer(-label) %>% 
mutate(label = fct_relevel(label, tip_labels$label), name=fct_relevel(name, c("Style","Habitat"))) 
nr <- sum %>%  mutate(label = fct_relevel(label, tip_labels$label))


l <- read.table("lacking_list", header=F)
colnames(l) <- c("sp","loss")

gene <- sum %>% 
	full_join(l) %>%
	pivot_longer(-c(sp,loss,Style,Habitat,label,taxa,nr)) %>% 
	mutate(
		alpha = ifelse(value>0,10,0), label = 
		fct_relevel(label, tip_labels$label), 
		gene = fct_relevel(name, c("ecr","usp","hr3","lin42","kin20","e75","ftzf1")),
		quest = ifelse(loss==3 & value<1,"?", ""),
		quest = ifelse(gene != "ecr" & gene != "usp"  & value==0, "?", quest))

g_l <- life %>% ggplot(aes(x=name,y=label,color=value,label=value)) + 
geom_text(size=3) + theme_classic() + 
theme(axis.text.y=element_blank(), axis.ticks=element_blank(), 
axis.title=element_blank(), axis.line.y=element_blank(), axis.line.x=element_line(size=0.5), axis.text=element_text(angle = 60, hjust = 1),legend.position="none") + 
scale_color_manual(values=c("#b28c6e","black","#507ea4","#3a5b52","#0f2350","#bb5548","#839b5c","black"))

g_nr <-  nr %>%  ggplot(aes(x=nr, y=label, fill=taxa)) + geom_barh(stat="identity") + 
theme(axis.text.y=element_blank(), axis.ticks.y=element_blank(), axis.title=element_blank(), legend.position="none", 
axis.text=element_text(angle = 60, hjust = 1), axis.line.y=element_blank(),  
axis.line.x=element_line(size=0.5), panel.background=element_rect(fill="transparent"), panel.grid.major.x=element_line(color="black", size=0.2), 
panel.grid.minor.x=element_line(color="black", size=0.1))

g_g1 <- gene %>% filter(gene == "ecr" | gene == "usp") %>%
ggplot() + geom_point(aes(x=gene, y=label, color=taxa,alpha=alpha),size=4,shape=15) + geom_text(aes(x=gene, y=label,label=quest))  + 
theme_classic()+ 
theme(axis.text.y=element_blank(), 
axis.ticks=element_blank(),axis.text=element_text(angle = 60, hjust = 1), axis.title=element_blank(), legend.position="none", axis.line.y=element_blank(), 
axis.line.x=element_line(size=0.5))

g_g2 <- gene %>% filter(gene != "ecr" & gene != "usp") %>%
ggplot() + geom_point(aes(x=gene, y=label, color=taxa,alpha=alpha),size=4,shape=15) + geom_text(aes(x=gene, y=label,label=quest))  +
theme_classic()+
theme(axis.text.y=element_blank(),
axis.ticks=element_blank(),axis.text=element_text(angle = 60, hjust = 1), axis.title=element_blank(), legend.position="none", axis.line.y=element_blank(),
axis.line.x=element_line(size=0.5))

sum1 <- p + g_l + g_g1  + g_g2 + plot_layout(widths = c(2.2,1,0.2,0.4))
ggsave(plot=sum1, file="gene_sum.png", width=7.5, height=25)

sum2 <- p + g_l + g_nr + plot_layout(widths = c(3,1,2))
ggsave(plot=sum2, file="nr_sum.png", width=8, height=25)
```
