## Supplementary table for "Multiple losses of ecdysone receptor genes in nematodes: an alternative evolutionary scenario of molting regulation": genome_quality_figure.R

```
library(patchwork)
library(tidyverse)
library(ggtree)
library(ape)
library(ggstance)

tree <- read.tree('nematode_phylogeny_newick_noPontonema')

sum <- read.table("species.list_sum")
colnames(sum) <- c("Style","Habitat","sp","label","id","clade","clade_n","taxa","file","nr","ECR","USP","HR3","LIN42","E75","FTZF1")

sum <- sum %>% as_tibble() %>% select(-c(id,clade,clade_n,file))

p <- ggtree(tree)  + geom_tiplab(size=3, align=T, linesize=0.3) + ggplot2::xlim(0, 45)
tip_labels <- p$data[p$data$isTip, ]
tip_labels <- tip_labels[order(tip_labels$y), "label"]

life <-  sum %>% select(label,Style,Habitat) %>% pivot_longer(-label) %>% 
mutate(label = fct_relevel(label, tip_labels$label), name=fct_relevel(name, c("Style","Habitat"))) 
nr <- sum %>%  mutate(label = fct_relevel(label, tip_labels$label))

l <- read.table("lacking_list", header=F)
colnames(l) <- c("sp","loss")

gene <- sum %>% 
	full_join(l) %>%
	pivot_longer(-c(sp,loss,Style,Habitat,label,taxa,nr)) %>% 
	mutate(
		alpha = ifelse(value>0,10,0), label = 
		fct_relevel(label, tip_labels$label), 
		gene = fct_relevel(name, c("ECR","USP","HR3","LIN42","E75","FTZF1")),
		quest = ifelse(loss==3 & value<1,"?", ""),
		quest = ifelse(gene != "ECR" & gene != "USP"  & value==0, "?", quest)) %>%
	filter(label != "Pontonema_vulgare")

l2 <- read.table("name_change")
colnames(l2) <- c("Species_Name","sp")

add <- sum %>% select(label, taxa)
colnames(add) <- c("sp","taxa")

d <- read.csv("wormbase_parasite_select")

quality <- d %>% as_tibble()  %>% select(-c(Provider,Assembly, Clade,Genome_Browser)) %>% mutate(N50 = as.numeric(gsub(",", "",N50)), Genome_Size = as.numeric(gsub(",", "",Genome_Size)), 
Number_of_Scaffolds = as.numeric(gsub(",", "",Number_of_Scaffolds))) %>% group_nest(Species_Name) %>% mutate(N50 = map_dbl(data, ~mean(.$N50)), Genome_size = map_dbl(data, ~mean(.$Genome_Size)), 
Scaffold_number = map_dbl(data, ~mean(.$Number_of_Scaffolds))) %>% select(-data) %>% pivot_longer(-Species_Name) 

quality <- quality %>% full_join(l2) %>% left_join(add)  %>% mutate(label = fct_relevel(sp, tip_labels$label)) 

g_qu1 <- 
quality %>% full_join(l2) %>% left_join(add)  %>% mutate(label = fct_relevel(sp, tip_labels$label)) %>% ggplot(aes(x=value, y=label, fill=taxa)) + geom_barh(stat="identity") + facet_wrap(~name, 
scale="free") + theme(axis.text.y=element_blank(), axis.ticks.y=element_blank(), axis.title=element_blank(), legend.position="none",axis.text=element_text(angle = 60, hjust = 1), 
axis.line.y=element_blank(),axis.line.x=element_line(size=0.5), strip.background=element_blank(),panel.background=element_rect(fill="transparent"), panel.grid.major.x=element_line(color="black", 
size=0.2),panel.grid.minor.x=element_line(color="black", size=0.1)) + scale_x_log10()

g_qu2 <-
quality %>% full_join(l2) %>% left_join(add)  %>% mutate(label = fct_relevel(sp, tip_labels$label)) %>% ggplot(aes(x=value, y=label, fill=taxa)) + geom_barh(stat="identity") + facet_wrap(~name,
scale="free") + theme(axis.text.y=element_blank(), axis.ticks.y=element_blank(), axis.title=element_blank(), legend.position="none",axis.text=element_text(angle = 60, hjust = 1),
axis.line.y=element_blank(),axis.line.x=element_line(size=0.5), strip.background=element_blank(),panel.background=element_rect(fill="transparent"), panel.grid.major.x=element_line(color="black",
size=0.2),panel.grid.minor.x=element_line(color="black", size=0.1)) 


e <- read.table("busco_score")
colnames(e) <- c("Species_Name","id","an","as")

busco <-  e %>% group_nest(Species_Name) %>% mutate(BUSCO_annotation = map_dbl(data, ~mean(.$an)), BUSCO_assembly = map_dbl(data, ~mean(.$as)))  %>% select(-data) %>% left_join(l2) %>% 
left_join(add)  %>% mutate(label = fct_relevel(sp, tip_labels$label)) %>% pivot_longer(-c(sp,label,Species_Name,taxa)) 

g_bu <- busco %>% ggplot(aes(x=value, y=label, fill=taxa)) + geom_barh(stat="identity") + facet_wrap(~name,
scale="free") + theme(axis.text.y=element_blank(), axis.ticks.y=element_blank(), axis.title=element_blank(), legend.position="none",axis.text=element_text(angle = 60, hjust = 1),
axis.line.y=element_blank(),axis.line.x=element_line(size=0.5), panel.background=element_rect(fill="transparent"), strip.background=element_blank(),
panel.grid.major.x=element_line(color="black",size=0.2),panel.grid.minor.x=element_line(color="black", size=0.1)) 

g_g1 <- gene %>% filter(gene == "ECR" | gene == "USP") %>% ggplot() + geom_point(aes(x=gene, y=label, color=taxa,alpha=alpha),size=4,shape=15) + geom_text(aes(x=gene, y=label,label=quest))  + 
theme_classic()+ 
theme(axis.text.y=element_blank(), 
axis.ticks=element_blank(),axis.text=element_text(angle = 60, hjust = 1), axis.title=element_blank(), legend.position="none", axis.line.y=element_blank(), 
axis.line.x=element_line(size=0.5))

sum1 <- p + g_g1  + g_qu1 + plot_layout(widths = c(2.2,0.2,6))
ggsave(plot=sum1, file="Fig.S3_genome_quality_sum_log10.png", width=18, height=25)

sum2 <- p + g_g1  + g_qu2 + plot_layout(widths = c(2.2,0.2,6))
ggsave(plot=sum2, file="Fig.S4_genome_quality_sum.png", width=18, height=25)

sum3 <- p + g_g1  + g_bu  + plot_layout(widths = c(2.2,0.2,4))
ggsave(plot=sum3, file="Fig.S5_busco_sum.png", width=13, height=25)
```
