## Supplementary table for "Multiple losses of ecdysone receptor genes in nematodes: an alternative evolutionary scenario of molting regulation": blast_phylo.sh

```
#mkdir blast_num
#mkdir phylo
#mkdir db

for i in EcR usp Hr3 

do

rm blast_num/"blast_num_"$i
rm $i"_select.fa"

	 cat species.list2 | awk -F"\t" '{print $6}' | while read line
		do 
	
                rm top & rm top.fa & rm r.top & rm r.top.T & rm r.top.fa & rm r.top.HMM & rm hmm

		makeblastdb -in ../all_seq2/$line -out db/$line"_db" -dbtype prot -parse_seqids 
		blastp -query ../query/$i  -db db/$line"_db" -outfmt 7 -num_threads 6 > result
		cat result |  grep -v "#" | awk '$11<1e-20 {print $2}' > top

		cat ../all_seq2/$line | seqkit grep -nrf top > top.fa
		tblastn -query top.fa -db ../test/fmel.db -outfmt 7 -num_threads 6 | awk '/hits found/{getline;print}' | grep -v "#" | awk '$11<1e-10 {print $1" "$2}' | grep $i > r.top
	
		cat r.top | grep $i | awk '{print $1}' | sort | uniq > r.top.T	
		cat ../all_seq2/$line | seqkit grep -nrf r.top.T > r.top.fa
	
		cat r.top.fa | grep '>' | wc -l > n
		echo $line > nn
		paste n nn >> blast_num/"blast_num_"$i

		cat r.top.fa  >> $i"_select.fa"

		done
	
	cat species.list2 | awk -F"\t" '{print $1}' | while read line
		do 
		cat blast_num/"blast_num_"$i  | grep $line > a
                cat species.list2 | grep $line | awk -F"\t" 'BEGIN{OFS="\t"}{print $2,$5}' > b
                paste b a | awk -F"\t" 'BEGIN{OFS="\t"}{print $2,$3,$1}' >>c
		done

	cat c | sort -k 1 > blast_num/"blast_num_sort_"$i

	echo $i > a

	rm c

cat $i"_select.fa" ../phylo/NR.fa > ./phylo/$i"_set.fa"
mafft --auto ./phylo/$i"_set.fa" > ./phylo/$i"_set.fa_mafft"
trimal -in ./phylo/$i"_set.fa_mafft" -out ./phylo/$i"_set.fa_mafft_tr" -gt 0.8

cd ./phylo
raxmlHPC-PTHREADS-SSE3 -f a -x 12345 -p 12345 -# 1000 -m PROTGAMMAAUTO —-auto-prot=aic -s $i"_set.fa_mafft_tr"  -n $i"_set.fa_mafft_tr_240609_bp1000"  -T 7
cd ../

done

rm a
rm b
rm n
rm nn
rm result
rm top & rm top.fa & rm r.top & rm r.top.T & rm r.top.fa & rm r.top.HMM & rm hmm
```
