## Supplementary table for "Multiple losses of ecdysone receptor genes in nematodes: an alternative evolutionary scenario of molting regulation": phylo_sub1.sh

```
for i in EcR usp Hr3
do 
	cat $i"_select_S1_240613" | grep NEMA | awk '{print $2}' | sed -e s/\"//g > a
        cat ../$i"_select.fa" | seqkit grep -nrf a > a.fa

	cat a | awk '{print $1"____"}' > a2 
	cat a.fa | grep '>'  | awk '{print $1"____"}' | sed -e s/">"//g > b2
	cat a2 b2 | sort | uniq -c | sort -n | awk '$1==1 {print $2}' | sed -e s/____//g > c 

	cat a.fa | seqkit grep -v -nrf c > $i"_select.fa_1st_screen_1"

	rm a
	rm a2
	rm b2
	rm c
	rm a.fa

	cat ../../phylo/NR.fa $i"_select.fa_1st_screen_1" > $i"_select.fa_1st_screen_1_NR.set"
	mafft --thread 7 --auto $i"_select.fa_1st_screen_1_NR.set" >  ./subsequent1/$i"_select.fa_1st_screen_1_NR.set.mafft"
	trimal -in ./subsequent1/$i"_select.fa_1st_screen_1_NR.set.mafft"  -out ./subsequent1/$i"_select.fa_1st_screen_1_NR.set.mafft.tr0.8"  -gt 0.8


	cd ./subsequent1/
	raxmlHPC-PTHREADS-SSE3 -f a -x 12345 -p 12345 -# 1000 -m PROTGAMMAAUTO —-auto-prot=aic -s $i"_select.fa_1st_screen_1_NR.set.mafft.tr0.8" -n $i"_select.fa_1st_screen_1_NR.set.mafft.tr0.8"  -T 7
	cd ../

done
```
