## Supplementary table for "Multiple losses of ecdysone receptor genes in nematodes: an alternative evolutionary scenario of molting regulation": phylo_sub2.sh

```
cat ../species.list2| grep Rhabditina | awk '{print $1}' > Rha
cat ../species.list2| grep Tylenchina | awk '{print $1}'  > Tyl
cat ../species.list2| grep Spiruri | awk '{print $1}' > Spi
cat ../species.list2| grep -v Spiruri | grep -v Rhabditina | grep -v Tylenchina | awk '{print $1}' > Others

for i in Rha Tyl Spi Others
do

	cat Hr3_select.fa_1st_screen_1 | seqkit grep -nrf $i > "Hr3_select.fa_1st_screen_2_"$i

	cat ../../phylo/NR.fa "Hr3_select.fa_1st_screen_2_"$i > "Hr3_select.fa_1st_screen_2_"$i"_NR.set"
	mafft --thread 7 --auto  "Hr3_select.fa_1st_screen_2_"$i"_NR.set" >  ./subsequent2/"Hr3_select.fa_1st_screen_2_"$i"_NR.set.mafft"
	trimal -in ./subsequent2/"Hr3_select.fa_1st_screen_2_"$i"_NR.set.mafft"   -out ./subsequent2/"Hr3_select.fa_1st_screen_2_"$i"_NR.set.mafft0.8"  -gt 0.8

	cd ./subsequent2/
	raxmlHPC-PTHREADS-SSE3 -f a -x 12345 -p 12345 -# 1000 -m PROTGAMMAAUTO —-auto-prot=aic -s "Hr3_select.fa_1st_screen_2_"$i"_NR.set.mafft0.8" -n "Hr3_select.fa_1st_screen_2_"$i"_NR.set.mafft0.8" -T 7
	cd ../

done
```
