## Supplementary table for "Multiple losses of ecdysone receptor genes in nematodes: an alternative evolutionary scenario of molting regulation": tree_select_1.R

```
library(ggtree)
library(ape)
library(tidyverse)

#EcR
tree <- read.tree("phylo/RAxML_bipartitions.EcR_set.fa_mafft_tr_240609_bp1000")

get_node_number <- function(tree, tip_label) {
  tip_nodes <- which(tree$tip.label == tip_label)
  if (length(tip_nodes) == 0) {
    cat("Tip label", tip_label, "not found in the tree\n")
    return(NULL)
  } else {
    return(tip_nodes)
  }
}

tip_label <- "DrPPARAatr"
outgroup <- get_node_number(tree, tip_label)

rtree_EcR <- root(tree, outgroup = outgroup)
g <- ggtree(rtree_EcR)+ geom_text(aes(label=node), hjust=-.2) + geom_tiplab()

ggsave(file="./tree/EcR_root_S1_bp1000_240613.png", plot=g, width=20, height=120, limitsize = FALSE)

#Hr3
tree <- read.tree("phylo/RAxML_bipartitions.Hr3_set.fa_mafft_tr_240609_bp1000")

get_node_number <- function(tree, tip_label) {
  tip_nodes <- which(tree$tip.label == tip_label)
  if (length(tip_nodes) == 0) {
    cat("Tip label", tip_label, "not found in the tree\n")
    return(NULL)
  } else {
    return(tip_nodes)
  }
}

tip_label <- "DrPPARAatr"
outgroup <- get_node_number(tree, tip_label)

rtree_Hr3 <- root(tree, outgroup = outgroup)
g <- ggtree(rtree_Hr3)+ geom_text(aes(label=node), hjust=-.2) + geom_tiplab()

ggsave(file="./tree/Hr3_root_S1_bp1000_240613.png", plot=g, width=20, height=120, limitsize = FALSE)

#usp
tree <- read.tree("phylo/RAxML_bipartitions.usp_set.fa_mafft_tr_240609_bp1000")

get_node_number <- function(tree, tip_label) {
  tip_nodes <- which(tree$tip.label == tip_label)
  if (length(tip_nodes) == 0) {
    cat("Tip label", tip_label, "not found in the tree\n")
    return(NULL)
  } else {
    return(tip_nodes)
  }
}

tip_label <- "DrPPARAatr"
outgroup <- get_node_number(tree, tip_label)

rtree_usp <- root(tree, outgroup = outgroup)
g <- ggtree(rtree_usp)+ geom_text(aes(label=node), hjust=-.2) + geom_tiplab()

ggsave(file="./tree/usp_root_S1_bp1000_240613.png", plot=g, width=20, height=120, limitsize = FALSE)
```
