## Supplementary table for "Multiple losses of ecdysone receptor genes in nematodes: an alternative evolutionary scenario of molting regulation": tree_select_2.R

```
#EcR
EcR <- extract.clade(rtree_EcR, 440)$tip.label

sink("tree/EcR_select_S1_240613")
EcR
sink()

#Hr3
Hr3 <- extract.clade(rtree_Hr3, 860)$tip.label

sink("tree/Hr3_select_S1_240613")
Hr3
sink()

#usp
usp <- extract.clade(rtree_usp, 533)$tip.label

sink("tree/usp_select_S1_240613")
usp
sink()
```
