## Supplementary table for "Multiple losses of ecdysone receptor genes in nematodes: an alternative evolutionary scenario of molting regulation": blast.sh

```
for i in EcR usp Hr3
do

cat $i"_having.sp"  |while read line 
	do 

	cat ../screen_1st/$i"_select.fa_1st_screen_1" | seqkit grep -nrp $line > a
	cat $i"_lacking.sp" |  while read line2
		do 
		
		blastp -query a -db ../db/$line2".fa_db" -outfmt 7 -num_threads 7 > r
		cat r | awk '/hits found/{getline;print}' | grep -v "#" |  awk '$11<1e-20 {print}' | cut -f 1,2 > b
		cat b | awk '{print $2}' > c 
		cat c | while read line3
			do 

			cat ../../all_seq2/$line2".fa" | seqkit grep -nrp $line3 >> c.fa
		
			done 

		blastp -query c.fa -db ../db/$line".fa_db"  -outfmt 7 -num_threads 7 > r2
		cat r2 | awk '/hits found/{getline;print}' | grep -v "#" | awk '$11<1e-20 {print}'  |cut -f 1,2 > d

		paste b d | awk '{print $1"\t"$2"\t"$3"\t"$4}' >> "blast.tophit".$i

		rm c.fa

		done
	done

done
```
