## Supplementary table for "Multiple losses of ecdysone receptor genes in nematodes: an alternative evolutionary scenario of molting regulation": final_phylo.sh

```
mafft --thread 7 --auto screen2-1.set.fa >  screen2-1.set.fa.mafft
trimal -in screen2-1.set.fa.mafft   -out screen2-1.set.fa.mafft.tr0.8  -gt 0.8


raxmlHPC-PTHREADS-SSE3 -f a -x 12345 -p 12345 -# 1000 -m PROTGAMMAAUTO —-auto-prot=aic -s screen2-1.set.fa.mafft.tr0.8 -n screen2-1.set.fa.mafft.tr0.8.1000 -T 7
```
