## Supplementary table for "Multiple losses of ecdysone receptor genes in nematodes: an alternative evolutionary scenario of molting regulation": tree_select_1.R

```
library(tidyverse)
library(ape)
library(ggtree)

tree <- read.tree("../RAxML_bipartitions.screen2_candidate_NR_set.mafft.tr.0.8_1000")

get_node_number <- function(tree, tip_label) {
  tip_nodes <- which(tree$tip.label == tip_label)
  if (length(tip_nodes) == 0) {
    cat("Tip label", tip_label, "not found in the tree\n")
    return(NULL)
  } else {
    return(tip_nodes)
  }
}

tip_label <- "DrPPARAatr"
outgroup <- get_node_number(tree, tip_label)

rtree <- root(tree, outgroup = outgroup)
g <- ggtree(rtree)+ geom_text(aes(label=node), hjust=-.2) + geom_tiplab()

ggsave(file="root.png", plot=g, width=20, height=120, limitsize = FALSE)


EcR <- extract.clade(rtree, 587)$tip.label
usp <- extract.clade(rtree, 822)$tip.label
Hr3 <- extract.clade(rtree, 1038)$tip.label


sink("select")
EcR
usp
Hr3
sink()
```
