## Supplementary table for "Multiple losses of ecdysone receptor genes in nematodes: an alternative evolutionary scenario of molting regulation": tree_select_2.R

```
library(tidyverse)
library(ape)
library(ggtree)

tree <- read.tree("RAxML_bipartitions.screen2-1.set.fa.mafft.tr0.8.1000")

get_node_number <- function(tree, tip_label) {
  tip_nodes <- which(tree$tip.label == tip_label)
  if (length(tip_nodes) == 0) {
    cat("Tip label", tip_label, "not found in the tree\n")
    return(NULL)
  } else {
    return(tip_nodes)
  }
}

tip_label <- "DrPPARAatr"
outgroup <- get_node_number(tree, tip_label)

rtree <- root(tree, outgroup = outgroup)
g <- ggtree(rtree)+ geom_text(aes(label=node), hjust=-.2) + geom_tiplab()

ggsave(file="select_root.png", plot=g, width=20, height=120, limitsize = FALSE)


EcR <- extract.clade(rtree, 543)$tip.label
usp <- extract.clade(rtree, 327)$tip.label
Hr3 <- extract.clade(rtree, 488)$tip.label


sink("EcR_2.1_select")
EcR
sink()

sink("usp_2.1_select")
usp
sink()

sink("Hr3_2.1_select")
Hr3
sink()
```
