## Supplementary table for "Multiple losses of ecdysone receptor genes in nematodes: an alternative evolutionary scenario of molting regulation": tophit_select_hmm_phylo.sh

```
cat blast.tophit.* | awk '$1==$4"_" {print $2}' | sort | uniq  > Rha_Tyl_EcR_usp_reciprocal_tophit.select_list

rm Rha_Tyl_EcR_usp_reciprocal_tophit.select_list.fa

cat Rha_Tyl_EcR_usp_reciprocal_tophit.select_list | while read line
do

echo $line | awk -F"_" '{print $2}' | while read line2
	do
	cat ../../all_seq2/$line2".fa" | seqkit grep -nrp $line >> Rha_Tyl_EcR_usp_reciprocal_tophit.select_list.fa
	done

done

hmmscan --cpu 7 -E 1e-05  --domtblout Rha_Tyl_EcR_usp_reciprocal_tophit.select_list.fa_HMM_NR ../../HMM/NR.hmm Rha_Tyl_EcR_usp_reciprocal_tophit.select_list.fa

cat Rha_Tyl_EcR_usp_reciprocal_tophit.select_list.fa_HMM_NR | grep PF00104.33 | awk '{print $4}' | sort | uniq |  grep -v -f ../screen_2nd/screen2_candidate_list> screen3_candidate_list

cat screen3_candidate_list | awk '{print $1"_"}' > rename.list
cat Rha_Tyl_EcR_usp_reciprocal_tophit.select_list.fa | sed -e s/" "/"_ "/g > rename.fa

cat rename.fa | seqkit grep -nrf rename.list > screen3_candidate_list.fa

cat ../../phylo/NR.fa screen3_candidate_list.fa > screen3_candidate_NR_set.fa

mafft --thread 7 --auto screen3_candidate_NR_set.fa >  screen3_candidate_NR_set.mafft
trimal -in screen3_candidate_NR_set.mafft  -out screen3_candidate_NR_set.mafft.tr.0.8  -gt 0.8

#raxmlHPC-PTHREADS-SSE3 -f a -x 12345 -p 12345 -# 1000 -m PROTGAMMAAUTO —-auto-prot=aic -s screen3_candidate_NR_set.mafft.tr.0.8 -n screen3_candidate_NR_set.mafft.tr.0.8_1000 -T 7
```
