## Supplementary table for "Multiple losses of ecdysone receptor genes in nematodes: an alternative evolutionary scenario of molting regulation": blast_phylo_common.sh

```
########BLAST

###making blast database
makeblastdb -in <input> -out <output> -dbtype prot -parse_seqids 

###blastp analysis (tubler format output result)
blastp -query <query>  -db <db> -outfmt 7 -num_threads 6 > result

###create top hit list with specific threshold (e.g. 1E-20)
cat result |  grep -v "#" | awk '$11<1e-20 {print $2}' > tophit_list


#########Alignment and RAxML

###alignment using mafft (--auto function)
mafft --auto input  > output

###Trimming of aligned sequences (threshold: 0.8)
trimal -in input  -out output -gt 0.8

###Making a gene tree using RAxML (ex. bootstrap: 1000 times)
raxmlHPC-PTHREADS-SSE3 -f a -x 12345 -p 12345 -# 1000 -m PROTGAMMAAUTO —-auto-prot=aic -s <input>  -n <output>  -T 7
```
