## Supplementary table for "Multiple losses of ecdysone receptor genes in nematodes: an alternative evolutionary scenario of molting regulation": Clustering_ac.R

```
ac_cl <- ac %>% select(id,stage,mean) %>% group_nest(id) %>% mutate(max = map_dbl(data, ~max(.$mean))) %>% 
filter(max>1) %>% select(-max) %>% unnest(data) %>% pivot_wider(id, names_from=stage, values_from=mean) 

ac_cl2 <- ac_cl %>% select(-id)
ac_cl3 <- as.matrix(ac_cl2)
rownames(ac_cl3) <- ac_cl$id

scaledata <- t(scale(t(ac_cl3))) 
scaledata <- scaledata[complete.cases(scaledata),]

mestimate<- function(df){
  N <-  dim(df)[[1]]
  D <- dim(df)[[2]]
  m.sj <- 1 + (1418/N + 22.05)*D^(-2) + (12.33/N +0.243)*D^(-0.0406*log(N) - 0.1134)
  return(m.sj)
}

m <- mestimate(scaledata)
m
#[1] 1.719721

sumsqr <- function(x, clusters){
  sumsqr <- function(x) sum(scale(x, scale = FALSE)^2)
  wss <- sapply(split(as.data.frame(x), clusters), sumsqr)
  return(wss)
}

#get the wss for repeated clustering
iterate_fcm_WSS <- function(df,m){
  totss <- numeric()
  for (i in 2:20){
    FCMresults <- cmeans(df,centers=i,m=m)
    totss[i] <- sum(sumsqr(df,FCMresults$cluster))
  }
  return(totss)
}
wss_2to20 <- iterate_fcm_WSS(scaledata,m)
plot(1:20, wss_2to20[1:20], type="b", xlab="Number of Clusters", ylab="wss")

k = 5
fcm_results <- cmeans(scaledata,centers=k,m=m)

library(reshape2)
library(tidyr)
library(dplyr)

#get the centroids into a long dataframe:
fcm_centroids <- fcm_results$centers
fcm_centroids_df <- data.frame(fcm_centroids)
fcm_centroids_df$cluster <- row.names(fcm_centroids_df)
centroids_long <- tidyr::gather(fcm_centroids_df,"sample",'value',1:6)

ggplot(centroids_long, aes(x=sample,y=value, group=cluster, colour=as.factor(cluster))) + 
  geom_line() +
  xlab("Time") +
  ylab("Expression") +
  labs(title= "Cluster Expression by Time",color = "Cluster")

cor(t(fcm_centroids))

#start with the input data
fcm_plotting_df <- data.frame(scaledata)

#add genes
fcm_plotting_df$gene <- row.names(fcm_plotting_df)

#bind cluster assinment
fcm_plotting_df$cluster <- fcm_results$cluster
#fetch the membership for each gene/top scoring cluster
fcm_plotting_df$membership <- sapply(1:length(fcm_plotting_df$cluster),function(row){
  clust <- fcm_plotting_df$cluster[row]
  fcm_results$membership[row,clust]
})

k_to_plot = 5

#subset the dataframe by the cluster and get it into long form
#using a little tidyr action
cluster_plot_df <- dplyr::filter(fcm_plotting_df, cluster == k_to_plot) %>%
  dplyr::select(.,1:6,membership,gene) %>%
  tidyr::gather(.,"sample",'value',1:6)

#order the dataframe by score
cluster_plot_df <- cluster_plot_df[order(cluster_plot_df$membership),]
#set the order by setting the factors using forcats
cluster_plot_df$gene = forcats::fct_inorder(cluster_plot_df$gene)

#subset the cores by cluster
core <- dplyr::filter(centroids_long, cluster == k_to_plot)

ac_g5 <- ggplot(cluster_plot_df, aes(x=sample,y=value)) + 
    geom_line(aes(colour=membership, group=gene)) +
    scale_colour_gradientn(colours=c('blue1','red2')) +
    #this adds the core 
    geom_line(data=core, aes(sample,value, group=cluster), color="black",inherit.aes=FALSE) +
    xlab("") +
    ylab("") +
    labs(title= paste0("Cluster ",k_to_plot),color = "Score") +
    theme(axis.line=element_line(color="black")) 

k_to_plot = 4

#subset the dataframe by the cluster and get it into long form
#using a little tidyr action
cluster_plot_df <- dplyr::filter(fcm_plotting_df, cluster == k_to_plot) %>%
  dplyr::select(.,1:6,membership,gene) %>%
  tidyr::gather(.,"sample",'value',1:6)
  
#order the dataframe by score
cluster_plot_df <- cluster_plot_df[order(cluster_plot_df$membership),]
#set the order by setting the factors using forcats
cluster_plot_df$gene = forcats::fct_inorder(cluster_plot_df$gene)

#subset the cores by cluster
core <- dplyr::filter(centroids_long, cluster == k_to_plot)

ac_g4 <- ggplot(cluster_plot_df, aes(x=sample,y=value)) +
    geom_line(aes(colour=membership, group=gene)) +
    scale_colour_gradientn(colours=c('blue1','red2')) +
    #this adds the core
    geom_line(data=core, aes(sample,value, group=cluster), color="black",inherit.aes=FALSE) +
    xlab("") +
    ylab("") +
    labs(title= paste0("Cluster ",k_to_plot),color = "Score") +
    theme(axis.line=element_line(color="black"))

k_to_plot = 3

#subset the dataframe by the cluster and get it into long form
#using a little tidyr action
cluster_plot_df <- dplyr::filter(fcm_plotting_df, cluster == k_to_plot) %>%
  dplyr::select(.,1:6,membership,gene) %>%
  tidyr::gather(.,"sample",'value',1:6)
  
#order the dataframe by score
cluster_plot_df <- cluster_plot_df[order(cluster_plot_df$membership),]
#set the order by setting the factors using forcats
cluster_plot_df$gene = forcats::fct_inorder(cluster_plot_df$gene)

#subset the cores by cluster
core <- dplyr::filter(centroids_long, cluster == k_to_plot)

ac_g3 <- ggplot(cluster_plot_df, aes(x=sample,y=value)) +
    geom_line(aes(colour=membership, group=gene)) +
    scale_colour_gradientn(colours=c('blue1','red2')) +
    #this adds the core
    geom_line(data=core, aes(sample,value, group=cluster), color="black",inherit.aes=FALSE) +
    xlab("") +
    ylab("") +
    labs(title= paste0("Cluster ",k_to_plot),color = "Score") +
    theme(axis.line=element_line(color="black"))

k_to_plot = 1

#subset the dataframe by the cluster and get it into long form
#using a little tidyr action
cluster_plot_df <- dplyr::filter(fcm_plotting_df, cluster == k_to_plot) %>%
  dplyr::select(.,1:6,membership,gene) %>%
  tidyr::gather(.,"sample",'value',1:6)
  
#order the dataframe by score
cluster_plot_df <- cluster_plot_df[order(cluster_plot_df$membership),]
#set the order by setting the factors using forcats
cluster_plot_df$gene = forcats::fct_inorder(cluster_plot_df$gene)

#subset the cores by cluster
core <- dplyr::filter(centroids_long, cluster == k_to_plot)

ac_g1 <- ggplot(cluster_plot_df, aes(x=sample,y=value)) +
    geom_line(aes(colour=membership, group=gene)) +
    scale_colour_gradientn(colours=c('blue1','red2')) +
    #this adds the core
    geom_line(data=core, aes(sample,value, group=cluster), color="black",inherit.aes=FALSE) +
    xlab("") +
    ylab("") +
    labs(title= paste0("Cluster ",k_to_plot),color = "Score") +
    theme(axis.line=element_line(color="black"))

k_to_plot = 2

#subset the dataframe by the cluster and get it into long form
#using a little tidyr action
cluster_plot_df <- dplyr::filter(fcm_plotting_df, cluster == k_to_plot) %>%
  dplyr::select(.,1:6,membership,gene) %>%
  tidyr::gather(.,"sample",'value',1:6)
  
#order the dataframe by score
cluster_plot_df <- cluster_plot_df[order(cluster_plot_df$membership),]
#set the order by setting the factors using forcats
cluster_plot_df$gene = forcats::fct_inorder(cluster_plot_df$gene)

#subset the cores by cluster
core <- dplyr::filter(centroids_long, cluster == k_to_plot)

ac_g2 <- ggplot(cluster_plot_df, aes(x=sample,y=value)) +
    geom_line(aes(colour=membership, group=gene)) +
    scale_colour_gradientn(colours=c('blue1','red2')) +
    #this adds the core
    geom_line(data=core, aes(sample,value, group=cluster), color="black",inherit.aes=FALSE) +
    xlab("") +
    ylab("") +
    labs(title= paste0("Cluster ",k_to_plot),color = "Score") +
    theme(axis.line=element_line(color="black"))

saveRDS(file="ac_g1.RDS", object=ac_g1)
saveRDS(file="ac_g2.RDS", object=ac_g2)
saveRDS(file="ac_g3.RDS", object=ac_g3)
saveRDS(file="ac_g4.RDS", object=ac_g4)
saveRDS(file="ac_g5.RDS", object=ac_g5)
saveRDS(file="ac_clu.RDS", object=fcm_plotting_df)

sink("ac_clu")
fcm_plotting_df %>% as_tibble() %>% select(gene, cluster) %>% print(n=100000)
sink()
```
