## Supplementary table for "Multiple losses of ecdysone receptor genes in nematodes: an alternative evolutionary scenario of molting regulation": AncylCeyla_filter_exp.sh

```
cat sra.list| awk '{print $1}' | while read line
do

java -jar /Users/shumpei/Trimmomatic-0.36/trimmomatic-0.36.jar  PE -threads 6   -phred33  $line"_1.fastq"  $line"_2.fastq" "PAIRED_"$line"_1.fastq"   "UNPAIRED_"$line"_1.fastq" "PAIRED_"$line"_2.fastq"  "UNPAIRED_"$line"_2.fastq" ILLUMINACLIP:/Users/shumpei/Dropbox/PD1/Tardigrade_RNAseq_data/Trimmomatic-0.38.fa:2:30:10 LEADING:20 TRAILING:20 SLIDINGWINDOW:4:15         MINLEN:36

fastqc --nogroup "PAIRED_"$line"_1.fastq" & fastqc --nogroup "PAIRED_"$line"_2.fastq"

rsem-prepare-reference --gff3 ancylostoma_ceylanicum.PRJNA231479.WBPS19.annotations.gff3_no_empty --bowtie2 -p 6 ancylostoma_ceylanicum.PRJNA231479.WBPS19.genomic.fa ancylostoma_ceylanicum.PRJNA231479.WBPS19.annotations.gff3_no_empty_RSEM.index

done


cat sra.list| awk '{print $1}' | while read i
do

rsem-calculate-expression --paired-end --bowtie2 --num-threads 7 "PAIRED_"$i"_1.fastq" "PAIRED_"$i"_2.fastq" ancylostoma_ceylanicum.PRJNA231479.WBPS19.annotations.gff3_no_empty_RSEM.index ./rsem/$i"_rsem"

cp rsem/$i"_rsem.genes.results" /Volumes/HD-PCFSU3-A/240501_AncCey/rsem/ & mv rsem/$i"_rsem.isoforms.results" /Volumes/HD-PCFSU3-A/240501_AncCey/rsem/ & mv rsem/$i"_rsem.log" /Volumes/HD-PCFSU3-A/240501_AncCey/rsem/ & mv rsem/$i"_rsem.transcript.bam" /Volumes/HD-PCFSU3-A/240501_AncCey/rsem/ & mv rsem/$i"_rsem.stat"/ /Volumes/HD-PCFSU3-A/240501_AncCey/rsem/

rm "PAIRED_"$i"_1.fastq"
rm "PAIRED_"$i"_2.fastq"

done
```
