## Supplementary table for "Multiple losses of ecdysone receptor genes in nematodes: an alternative evolutionary scenario of molting regulation": BrugiMalay_filter_exp.sh

```
for i in SRR9858310 SRR4308251 SRR9858311 SRR9858312 SRR9858313 SRR9858314 SRR9858315 SRR9858316 SRR9858317
do

conda activate /Users/shumpei/opt/miniconda3/envs/bioinfo_basic/

java -jar /Users/shumpei/Trimmomatic-0.36/trimmomatic-0.36.jar  PE -threads 7   -phred33  $i".sra_1.fastq" $i".sra_2.fastq"  "new_PAIRED_"$i".sra_1.fastq"     "new_UNPAIRED_"$i".sra_1.fastq" "new_PAIRED_"$i".sra_2.fastq"     "new_UNPAIRED_"$i".sra_2.fastq" ILLUMINACLIP:Trimmomatic-0.38.fa_3:2:30:10 LEADING:20  TRAILING:20 SLIDINGWINDOW:4:15  MINLEN:36 CROP:150

fastqc --nogroup "new_PAIRED_"$i".sra_1.fastq" & fastqc --nogroup "new_PAIRED_"$i".sra_2.fastq"

rm $i".sra_1.fastq"  & rm $i".sra_2.fastq" 

conda activate /Users/shumpei/opt/miniconda3/envs/bowtie2_rsem

rsem-prepare-reference --bowtie2 -p 6 BrugiaMalayi_rrna.fa_noduplicate BrugiaMalayi_rrna.index
rsem-calculate-expression --paired-end --bowtie2 --num-threads 7 "new_PAIRED_"$i".sra_1.fastq" "new_PAIRED_"$i".sra_2.fastq" BrugiaMalayi_rrna.index ./rsem_rrna/$i"_rsem_rrna"

samtools view -b -f 4  rsem_rrna/$i"_rsem_rrna.transcript.bam" > $i"_rrna"  
samtools view $i"_rrna" | cut -f 1 > $i"_no_rrna"

conda activate /Users/shumpei/opt/miniconda3/envs/bioinfo_basic/

cat "new_PAIRED_"$i".sra_1.fastq" | seqkit grep -f $i"_no_rrna" > $i"._no_rrna_sra_1.fastq"
cat "new_PAIRED_"$i".sra_2.fastq" | seqkit grep -f $i"_no_rrna" > $i"._no_rrna_sra_2.fastq"

fastqc --nogroup $i"._no_rrna_sra_1.fastq"  & fastqc --nogroup $i"._no_rrna_sra_2.fastq" 

conda activate /Users/shumpei/opt/miniconda3/envs/bowtie2_rsem
rsem-calculate-expression --paired-end --bowtie2 --num-threads 7 $i"._no_rrna_sra_1.fastq" $i"._no_rrna_sra_2.fastq" brugia_malayi.PRJNA10729.WBPS19.RSEM_index ./new_rsem/$i"_new_rsem"

done
```
