## Supplementary table for "Multiple losses of ecdysone receptor genes in nematodes: an alternative evolutionary scenario of molting regulation": Fig_Expression_pattern_others.R

```
library(tidyverse)
library(patchwork)


#######annot

annot <- read.table("gene_ids_kin-20_added", header=T)

annot_bm <- annot %>% filter(species=="Brugia_malayi") %>% select(-species)
annot_ac <- annot %>% filter(species=="Ancylostoma_ceylanicum") %>% select(-species)
annot_ts <- annot %>% filter(species=="Trichuris_suis") %>% select(-species)
annot_pp <- annot %>% filter(species=="Pristionchus_pacificus") %>% select(-species)
annot_gp <- annot %>% filter(species=="Globodera_pallida") %>% select(-species)
annot_ce <- annot %>% filter(species=="Caenorhabditis_elegans") %>% select(-species)

annot_colors <- c(   
"ecr" = "#eb6100",   
"usp" = "#004c97",   
"hr3" = "#15722c",   
"lin-42" = "#fccc00",
"e75" = "#914e25",  
"ftz-f1" = "#737061",
"kin-20" = "black"
)


#######Bm
bm <- read.table("../Brugia_malayi/new_rsem/new_rsem.tpm", header=T)


plot_bm <- bm %>% as_tibble() %>% right_join(annot_bm)  %>% filter(annotation != "ftz-f1_1")  %>% pivot_longer(-c(id,annotation)) %>%  separate(name, sep="_",into=c("st","rep")) %>% mutate(st =
str_replace(st,"X","")) %>% group_nest(id, st, annotation) %>% mutate(mean=map_dbl(data, ~mean(.$value)),time=ifelse(st=="3d",1,ifelse(st=="6d",2,ifelse(st=="9d",3,4)))) %>% separate(annotation,
sep="_", into=c("annot","copy")) %>% mutate(annot = fct_relevel(annot,c("ecr","usp","hr3","lin-42","e75")))  %>% group_by(annot) %>% group_split() %>% map(~ {
 ggplot(.x,aes(x=time,y=mean,color=annot,group=id)) +
geom_vline(xintercept=c(3.5),linetype="dotted") +  geom_line() +  scale_x_continuous(breaks=c(1:4), labels=c("L3_3d","L3-6d","L3-9d","L4")) + 
 theme_classic() + theme(legend.position="none", axis.title=element_blank())+ scale_color_manual(values = annot_colors)   + theme(axis.text=element_text(size=7))})


#######Ac
ac <- read.table("../Ancylostoma_ceylanicum/tpm.table",header=T)
ac <- ac %>% as_tibble() %>% pivot_longer(-id) %>% separate(name, into=c("st","time","bi","te"),sep="\\.") %>% group_nest(id,st,time,bi) %>% 
mutate(mean.bi = map_dbl(data, ~mean(.$value)))
ac <- ac %>% select(-data) %>% group_nest(id,st,time) %>% mutate(mean = map_dbl(data, ~mean(.$mean.bi)) )
ac <- ac %>% mutate(stage = ifelse(st=="L4",6,ifelse(time=="0h",1,ifelse(time=="24hint",2,ifelse(time=="24hstomach",3,ifelse(time=="48h",4,5)))))) 

plot_ac <- ac %>% as_tibble() %>% right_join(annot_ac) %>% separate(annotation, sep="_", into=c("annot","copy")) %>% complete(annot = c("ecr","usp","hr3","lin-42","e75","ftz-f1","kin-20")) %>% 
mutate(annot = 
fct_relevel(annot,c("ecr","usp","hr3","lin-42","e75"))) %>% group_by(annot) %>% group_split() %>% map(~ {ggplot(.x,aes(x=stage,y=mean,color=annot,group=id)) + 
geom_vline(xintercept=c(5.5),linetype="dotted") + geom_line(na.rm = TRUE)  + scale_x_continuous(breaks=c(1:6), labels=c("0","24i","24s","48","72","72"))  + theme_classic() + 
theme(legend.position="none", axis.title=element_blank()) + scale_color_manual(values = annot_colors)  + theme(axis.text=element_text(size=7))})


#######Ts
ts <- read.table("../exp.level.sum/SRP039506.tpm_per_run.tsv", header=T, sep="\t")
colnames(ts) <- c("id","10_1","16_1","17_1","21_1","28_1","35_1","42_1","a_8","a_7","a_1","a_2","a_3","a_4","a_5","a_6")
ts <- ts %>% as_tibble() %>% pivot_longer(-id) %>% separate(name, into=c("st","rep"), sep="_") %>% filter(st!="a") %>% mutate(st = 
as.numeric(st))


plot_ts <- ts %>% as_tibble() %>% right_join(annot_ts)%>% filter(annotation != "e75_1") %>% separate(annotation, sep="_", into=c("annot","copy")) %>% complete(annot = 
c("ecr","usp","hr3","lin-42","e75","ftz-f1","kin-20")) %>% mutate(annot = fct_relevel(annot,c("ecr","usp","hr3","lin-42","e75"))) %>%  group_by(annot) %>% group_split() %>% map(~ {
ggplot(.x,aes(x=st, y=value, color=annot, group=id)) + 
geom_vline(xintercept=c(11,22,33),linetype="dotted") + geom_line()   + theme_classic() + theme(legend.position="none", axis.title=element_blank())  + theme(axis.text=element_text(size=7)) +
 scale_color_manual(values = annot_colors) })


########Pp
Pp <- read.table("../Pristionchus_pacificus/Sun_et_al/Sun_et_al_tpm.table", header=T)
colnames(Pp) <- c("id",1:38)

Pp <- Pp %>% as_tibble() %>% 
mutate(ave_2 = (`1`+`2`+`3`)/3) %>%
mutate(ave_3 = (`2`+`3`+`4`)/3) %>%
mutate(ave_4 = (`3`+`4`+`5`)/3) %>%
mutate(ave_5 = (`4`+`5`+`6`)/3) %>%
mutate(ave_6 = (`5`+`6`+`7`)/3) %>%
mutate(ave_7 = (`6`+`7`+`8`)/3) %>%
mutate(ave_8 = (`7`+`8`+`9`)/3) %>%
mutate(ave_9 = (`8`+`9`+`10`)/3) %>%
mutate(ave_10 = (`9`+`10`+`11`)/3) %>%
mutate(ave_11 = (`10`+`11`+`12`)/3) %>%
mutate(ave_12 = (`11`+`12`+`13`)/3) %>%
mutate(ave_13 = (`12`+`13`+`14`)/3) %>%
mutate(ave_14 = (`13`+`14`+`15`)/3) %>%
mutate(ave_15 = (`14`+`15`+`16`)/3) %>%
mutate(ave_16 = (`15`+`16`+`17`)/3) %>%
mutate(ave_17 = (`16`+`17`+`18`)/3) %>%
mutate(ave_18 = (`17`+`18`+`19`)/3) %>%
mutate(ave_19 = (`18`+`19`+`20`)/3) %>%
mutate(ave_20 = (`19`+`20`+`21`)/3) %>%
mutate(ave_21 = (`20`+`21`+`22`)/3) %>%
mutate(ave_22 = (`21`+`22`+`23`)/3) %>%
mutate(ave_23 = (`22`+`23`+`24`)/3) %>%
mutate(ave_24 = (`23`+`24`+`25`)/3) %>%
mutate(ave_25 = (`24`+`25`+`26`)/3) %>%
mutate(ave_26 = (`25`+`26`+`27`)/3) %>%
mutate(ave_27 = (`26`+`27`+`28`)/3) %>%
mutate(ave_28 = (`27`+`28`+`29`)/3) %>%
mutate(ave_29 = (`28`+`29`+`30`)/3) %>%
mutate(ave_30 = (`29`+`20`+`31`)/3) %>%
mutate(ave_31 = (`30`+`31`+`32`)/3) %>%
mutate(ave_32 = (`31`+`32`+`33`)/3) %>%
mutate(ave_33 = (`32`+`33`+`34`)/3) %>%
mutate(ave_34 = (`33`+`34`+`35`)/3) %>%
mutate(ave_35 = (`34`+`35`+`36`)/3) %>%
mutate(ave_36 = (`35`+`36`+`37`)/3) %>%
mutate(ave_37 = (`36`+`37`+`38`)/3) 

Pp <- Pp %>% pivot_longer(-id) %>% separate(name, into=c("ave","time"), sep="_") %>% filter(ave=="ave")

Pp <- Pp %>% mutate(time=as.numeric(time), tpm=2^value) %>% select(id,time,tpm) %>% pivot_wider(id_cols=id, names_from=time, values_from=tpm) 

colnames(Pp) <- c("id",1,3,5,7,9,11,13,15,16,17,18,19:22,24,26,28:32,34,36,38,40,42:46,48,50,52,54,56)


plot_pp <- Pp %>% as_tibble() %>% right_join(annot_pp) %>% filter(annotation != "kin-20_1") %>% separate(annotation, sep="_", into=c("annot","copy")) %>% 
complete(annot =c("ecr","usp","hr3","lin-42","e75","ftz-f1","kin-20")) %>% mutate(annot 
= 
fct_relevel(annot,c("ecr","usp","hr3","lin-42","e75"))) %>% pivot_longer(-c(id,annot,copy)) %>% mutate(time=as.numeric(name), tpm=value) %>%  group_by(annot) %>% 
group_split() %>% map(~ {
ggplot(.x,aes(x=time,y=tpm, group=id, color=annot)) + 
geom_vline(xintercept=c(19,31,45), linetype="dotted") + geom_line() + theme_classic() + 
theme(legend.position="none", strip.background=element_blank(), 
strip.text=element_blank(), axis.title=element_blank()) + scale_color_manual(values = annot_colors)  + theme(axis.text=element_text(size=7))})


#########Gp
gp <- read.table("../exp.level.sum/ERP001236.tpm_per_run.tsv",header=T, sep="\t")
colnames(gp) <- c("id","14_1","14_2","21_1","21_2","28_1","28_2","35_1","35_2","7_1","35_3","35_4","0_1","0_2","0_3","a","b")
gp <- gp %>% as_tibble() %>% pivot_longer(-c(id)) %>% filter(name!="a" & name!="b") %>% separate(name, into=c("time","rep"), sep="_") %>% 
group_nest(id,time) %>% mutate(time=as.numeric(time), tpm=map_dbl(data,~mean(.$value)))


plot_gp <-  gp %>% as_tibble() %>% right_join(annot_gp) %>% separate(annotation, sep="_", into=c("annot","copy")) %>% 
complete(annot =c("ecr","usp","hr3","lin-42","e75","ftz-f1","kin-20")) %>% 
mutate(annot = 
fct_relevel(annot,c("ecr","usp","hr3","lin-42","e75","ftz-f1","kin-20"))) %>% group_nest(annot,time) %>% mutate(tpm2 = map_dbl(data, ~sum(.$tpm))) %>% unnest(data) %>% 
group_by(annot) %>% group_split() %>% map(~ {
ggplot(.x,aes(x=time,y=tpm2,color=annot,group=annot)) + geom_vline(xintercept=c(14,21,31),linetype="dotted") + geom_line()   + 
theme_classic() + theme(legend.position="none", axis.title=element_blank()) + scale_color_manual(values = annot_colors)  + theme(axis.text=element_text(size=7)) 
})


##########Ce
ce <- read.table("../Caenorhabditis_elegans/GSE49043_data_rpkm.txt",header=T)
h <- read.table("../Caenorhabditis_elegans/header", header=T)
colnames(ce) <- colnames(h)

ce <- ce %>% as_tibble() %>% pivot_longer(-c(GENEID, GENENAME)) %>% filter(grepl("DH2|DH5",name)) %>% 
separate(name,into=c("lin","time"),sep="N2_") %>% mutate(time=as.numeric(time)) 

ce <- ce %>% group_nest(GENEID,GENENAME,time) %>% mutate(rpkm = map_dbl(data, ~mean(.$value)))

annot_ce <- annot_ce %>% rename(GENENAME=id)

plot_ce <- ce  %>% as_tibble() %>% right_join(annot_ce) %>% separate(annotation, sep="_", into=c("annot","copy")) %>% 
complete(annot =c("ecr","usp","hr3","lin-42","e75","ftz-f1","kin-20")) %>% 
mutate(annot = fct_relevel(annot,c("ecr","usp","hr3","lin-42","e75","ftz-f1","kin-20")))   %>% 
group_by(annot) %>% group_split() %>% map(~ {
ggplot(.x,aes(x=time, y=rpkm, color=annot)) + geom_vline(xintercept=c(20,30,38,48),linetype="dotted") + geom_line()  + 
theme_classic() + theme(legend.position="none", axis.title=element_blank()) +  scale_color_manual(values = annot_colors)  + theme(axis.text=element_text(size=7))})

###########sum

g_sum <- 
plot_ce[[3]] + plot_ce[[4]] + plot_ce[[7]] + plot_ce[[5]] + plot_ce[[6]] +
plot_ac[[3]] + plot_ac[[4]] + plot_ac[[7]] + plot_ac[[5]] + plot_ac[[6]] +
plot_pp[[3]] + plot_pp[[4]] + plot_pp[[7]] + plot_spacer()+ plot_pp[[6]] + 
plot_gp[[3]] + plot_gp[[4]] + plot_gp[[7]] + plot_gp[[5]] + plot_gp[[6]] + 
plot_bm[[3]] + plot_bm[[4]] + plot_bm[[7]] + plot_bm[[5]] + plot_bm[[6]] +
plot_ts[[3]] + plot_ts[[4]] + plot_ts[[7]] + plot_ts[[5]] + plot_ts[[6]] + 
plot_layout(ncol=5)

ggsave(file="five_genes_matrix.png",plot=g_sum, width=8,height=8)
```
