## Supplementary figures and images for "Multiple losses of ecdysone receptor genes in nematodes: an alternative evolutionary scenario of molting regulation"

### ecdysozoan_ECR_USP_HR3.png

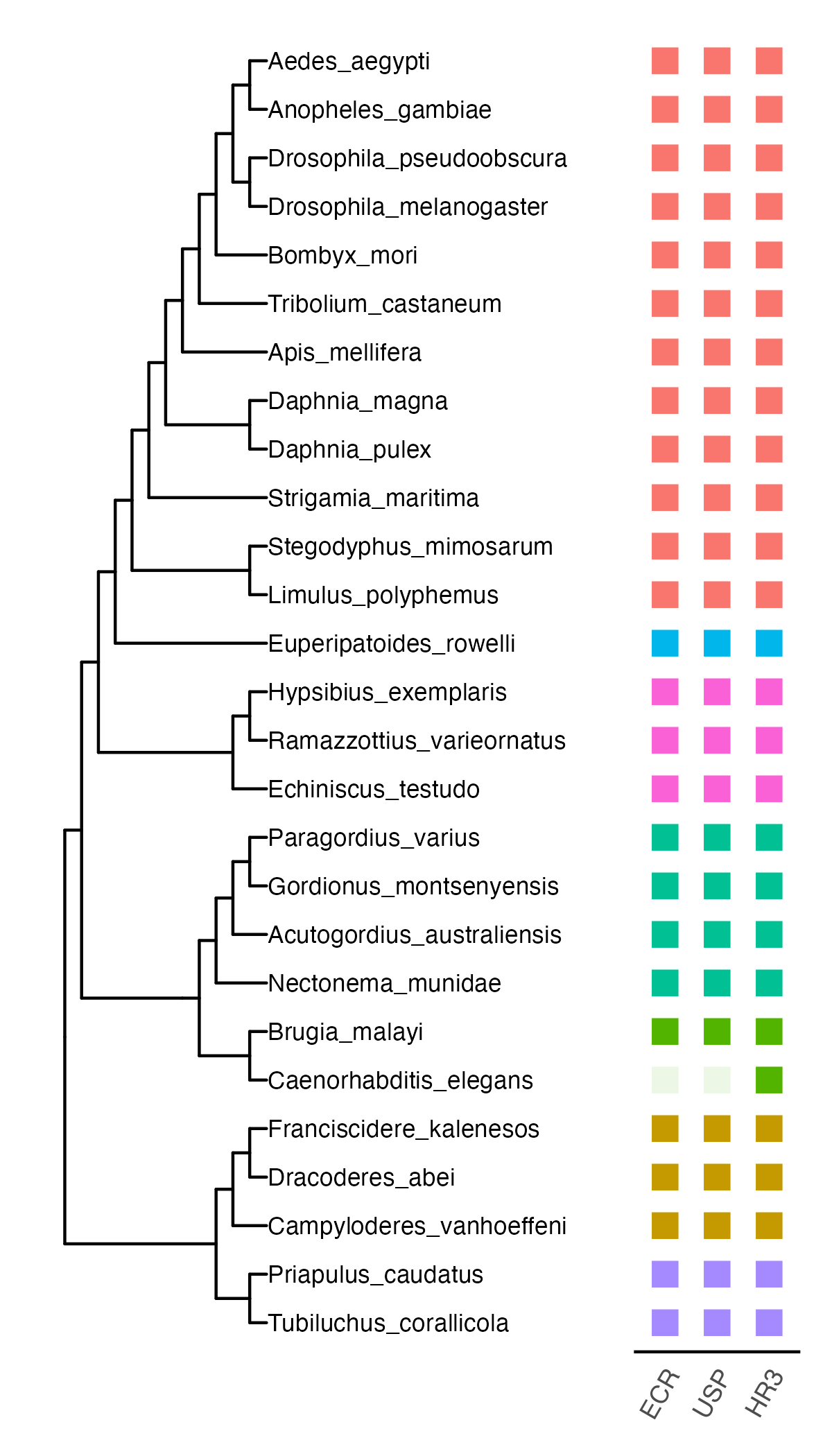

### EcR_root_S1_bp1000_240613.png

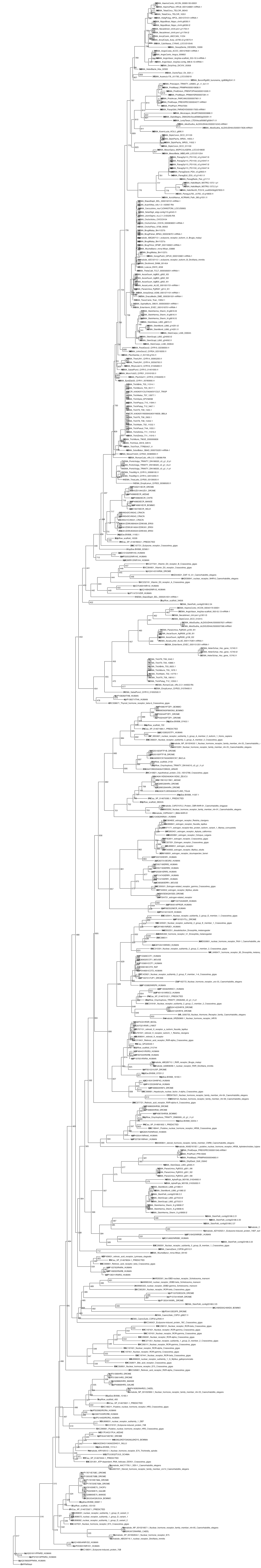

### Fig.2.png

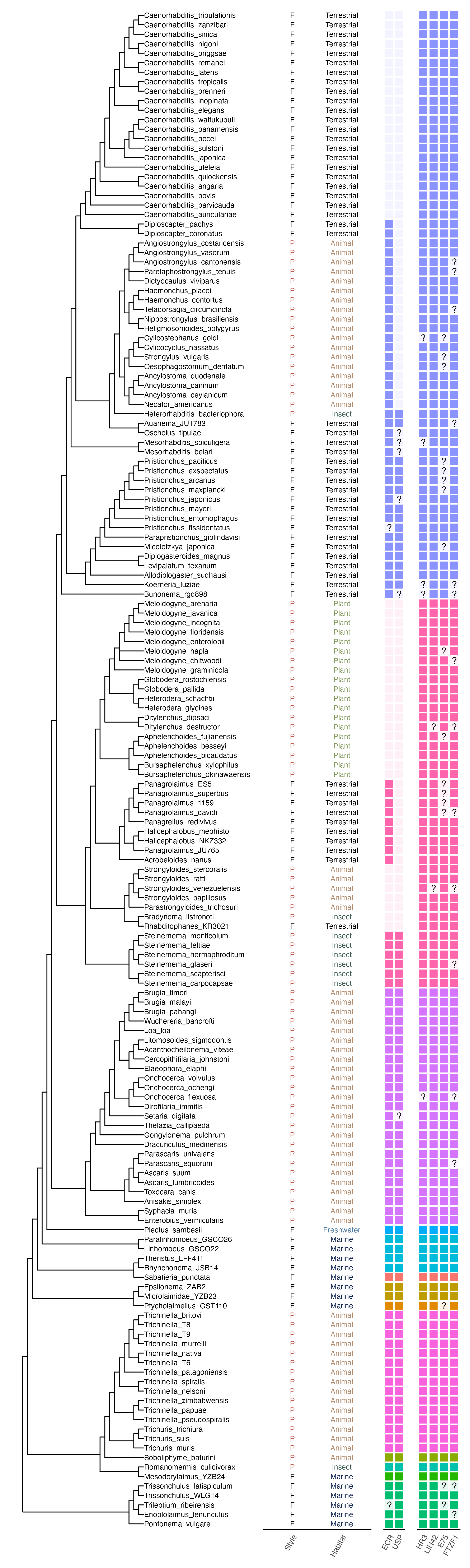

### Fig.6.png

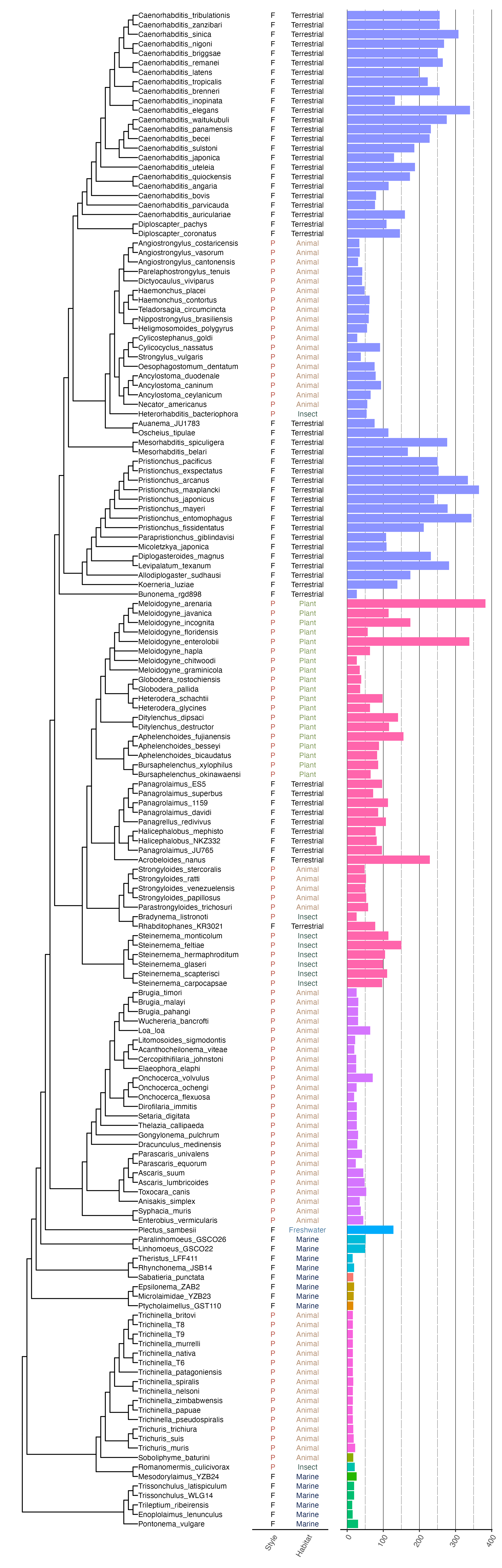

### Fig.S3_genome_quality_sum_log10.png

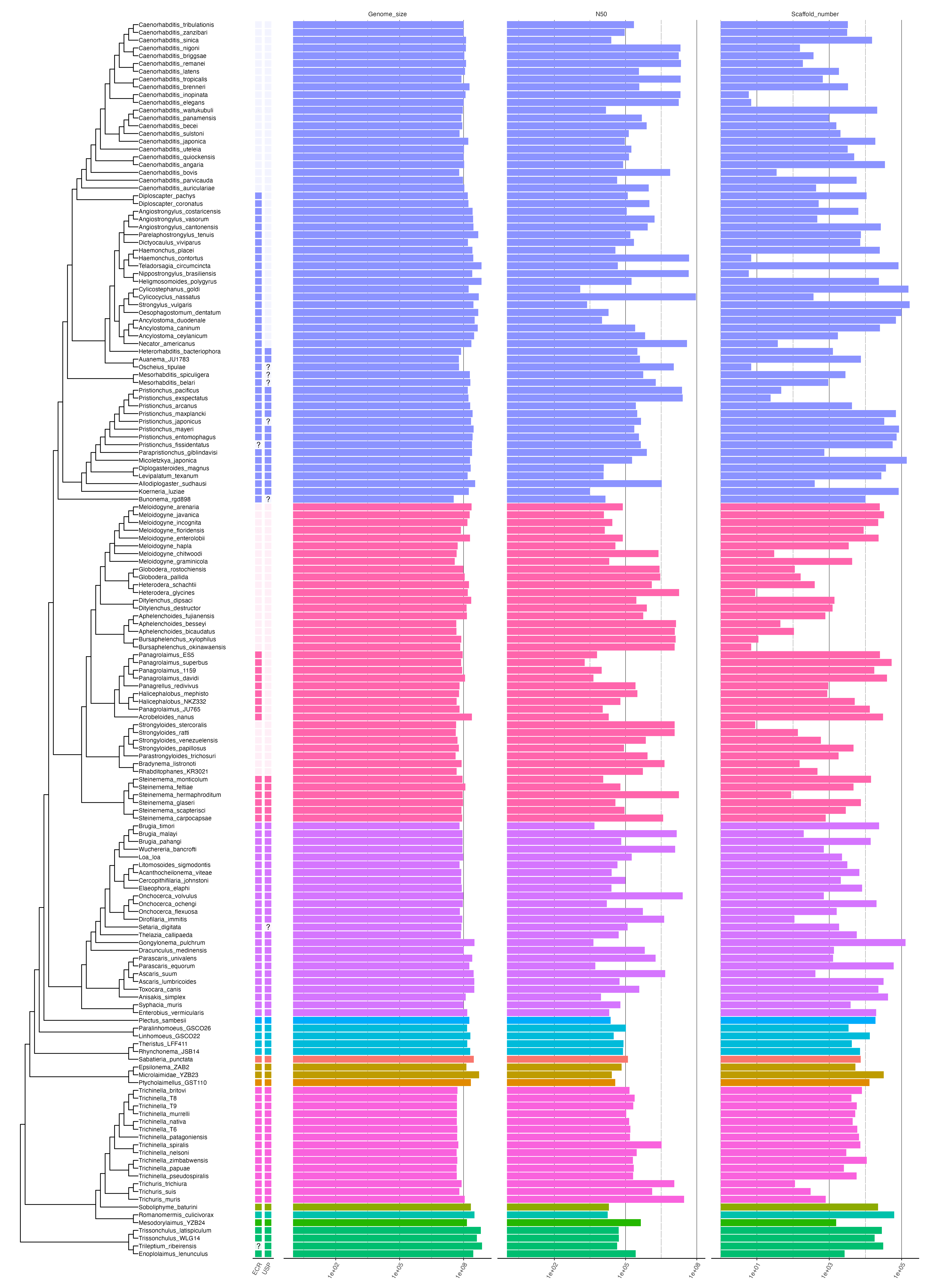

### Fig.S4_genome_quality_sum.png

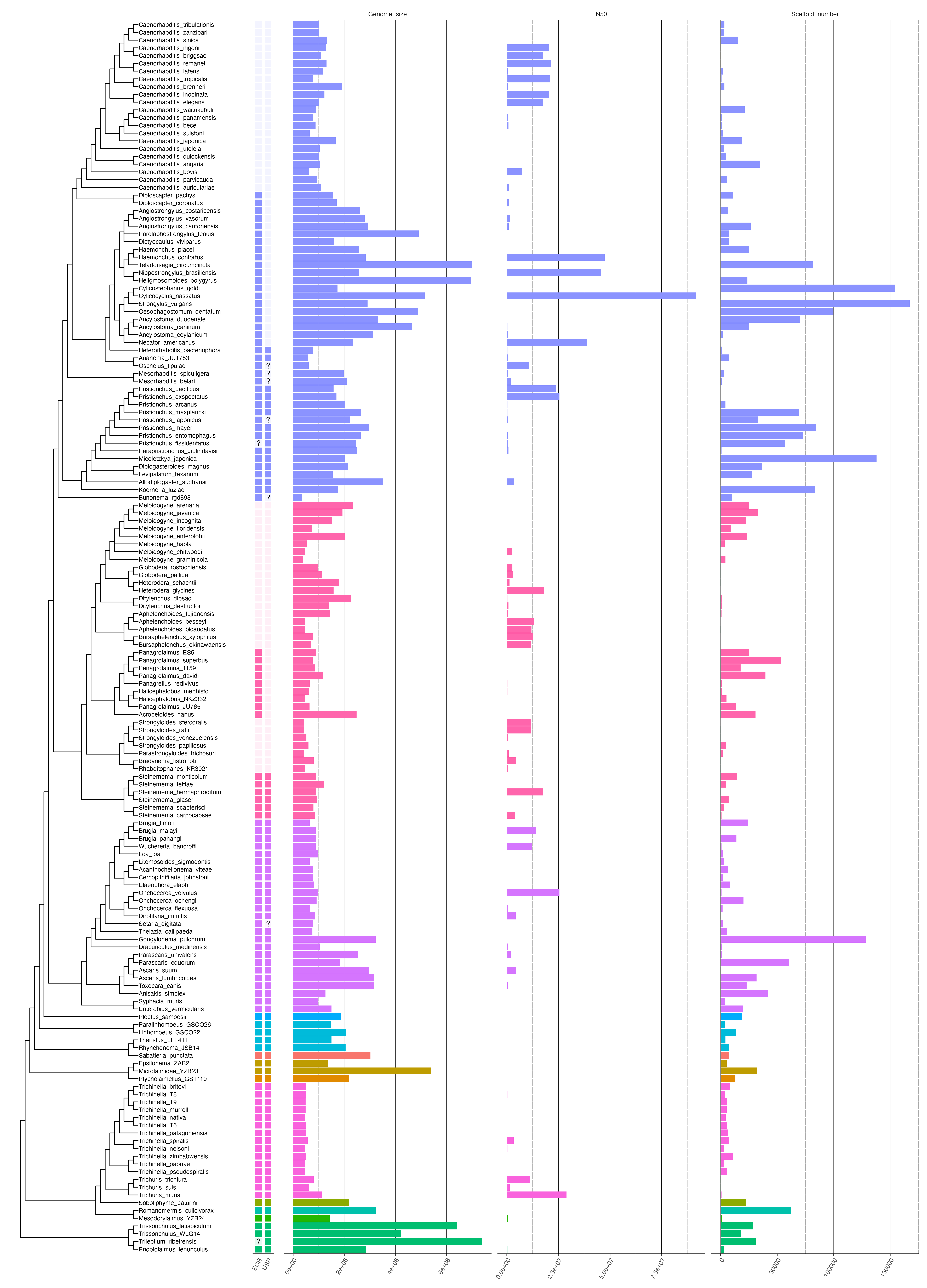

### Fig.S5_busco_sum.png

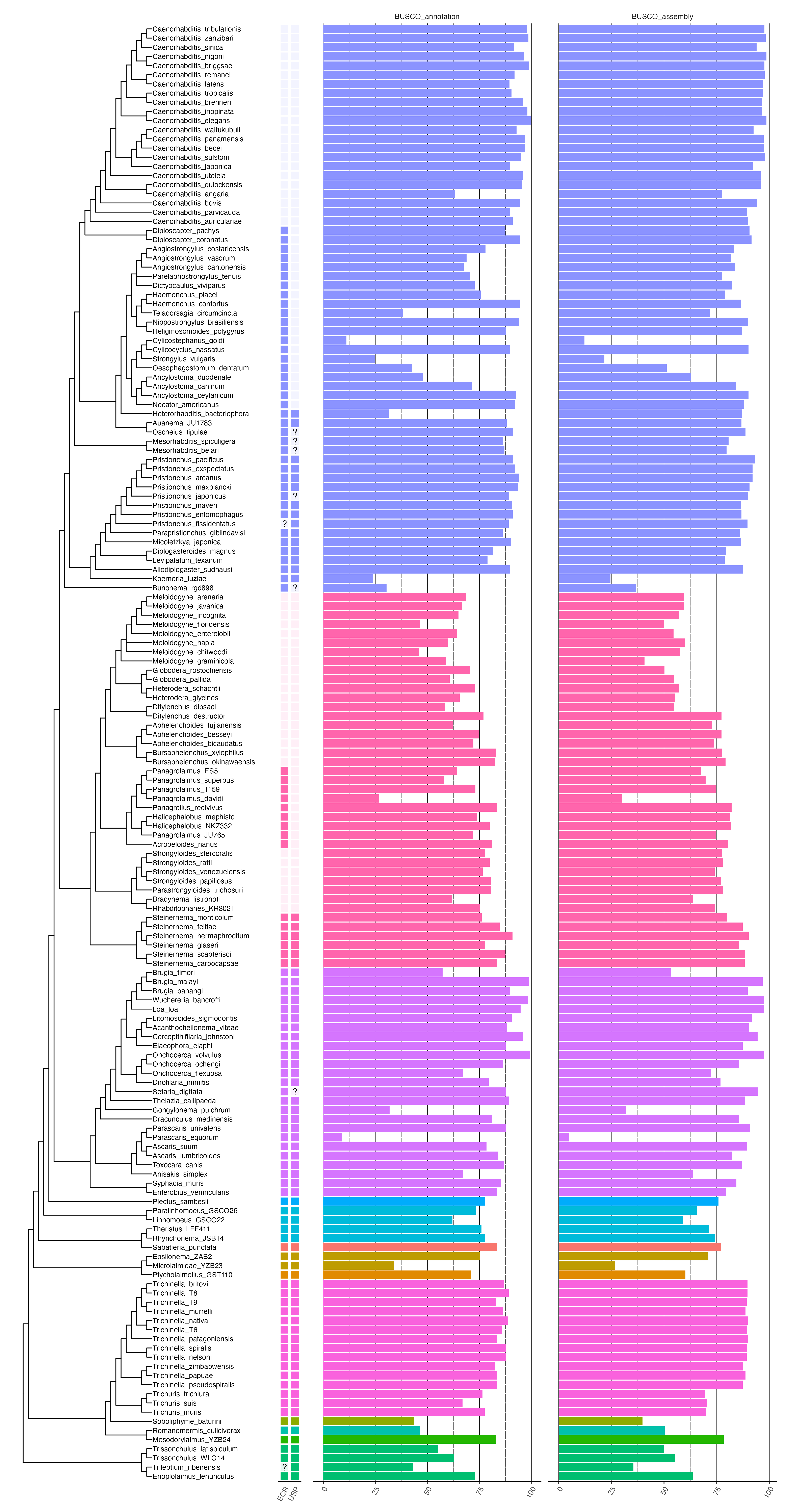

### Hr3_root_S1_bp1000_240613.png

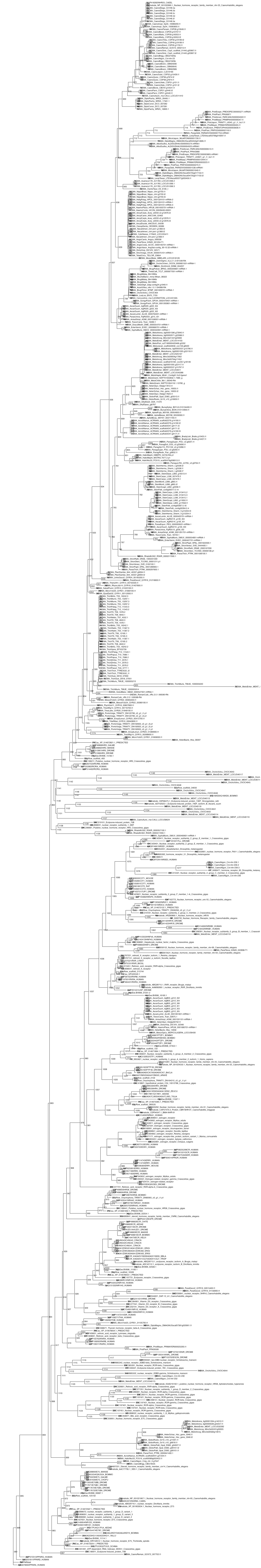

### root.png

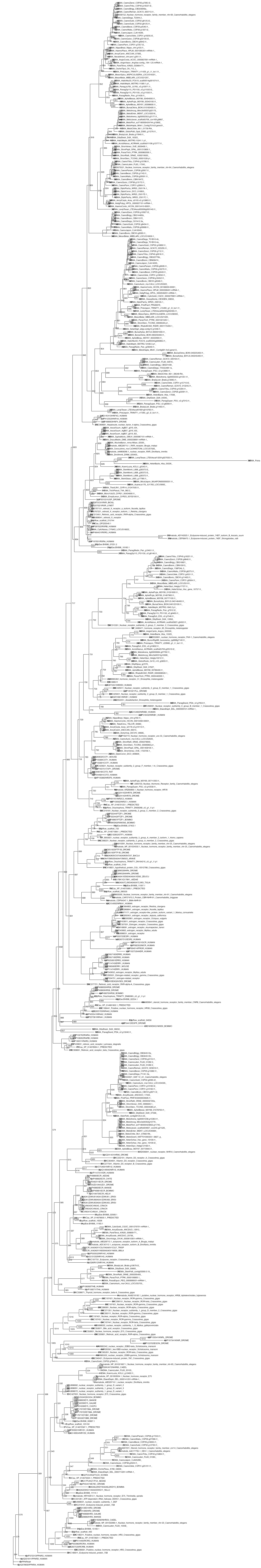

### select_root.png

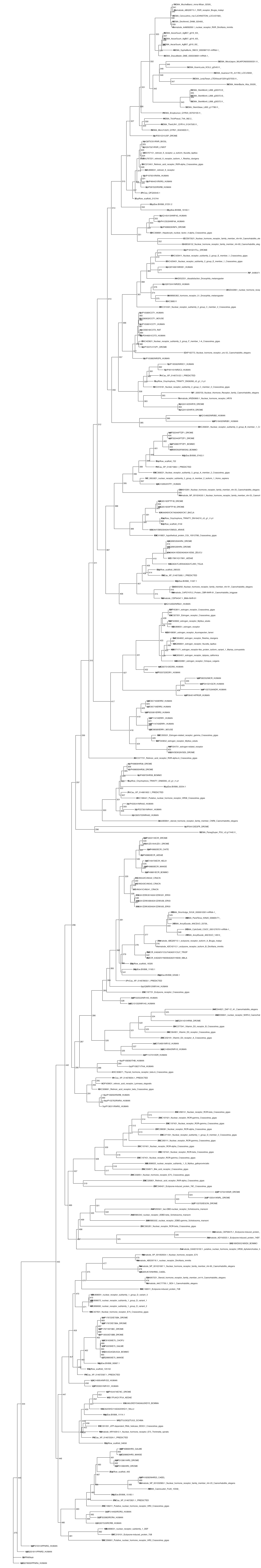

### tree_3rd_screening_E75_ftz-f1.pdf

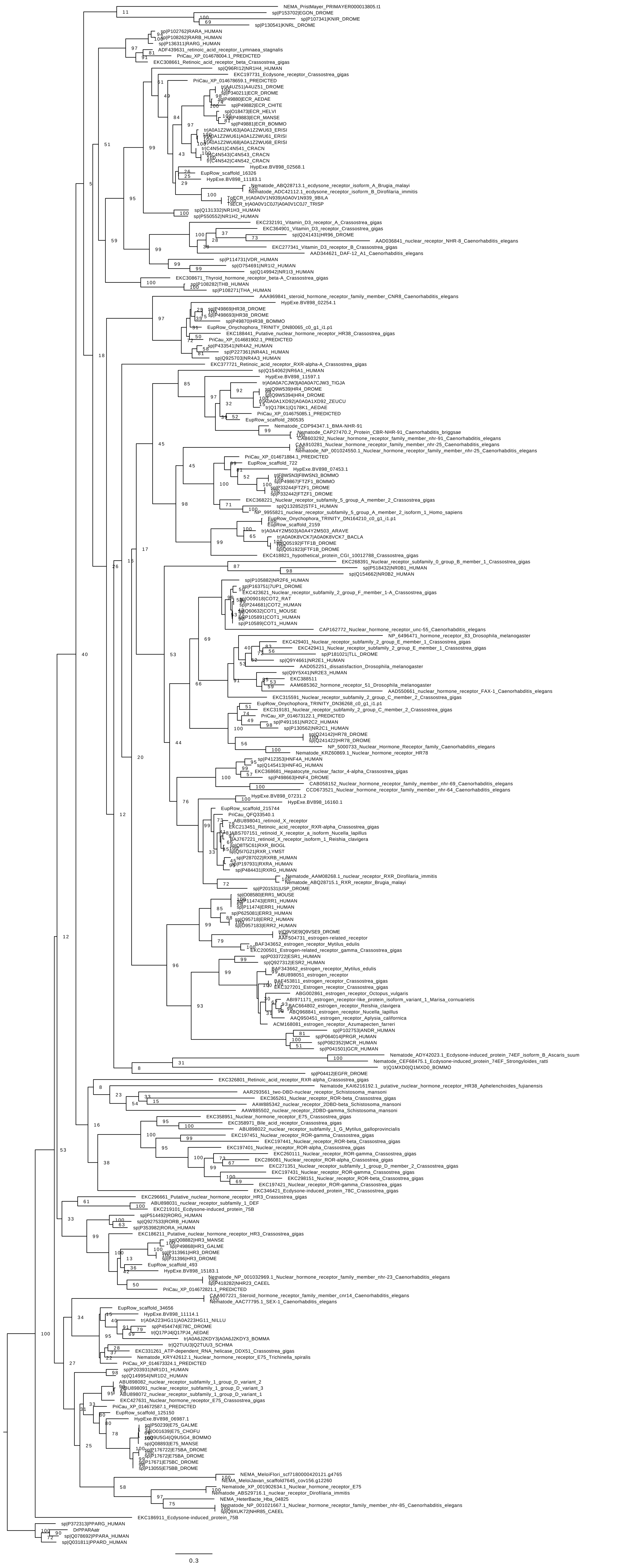

### tree_3rd_screening_ECR_USPHR3.pdf

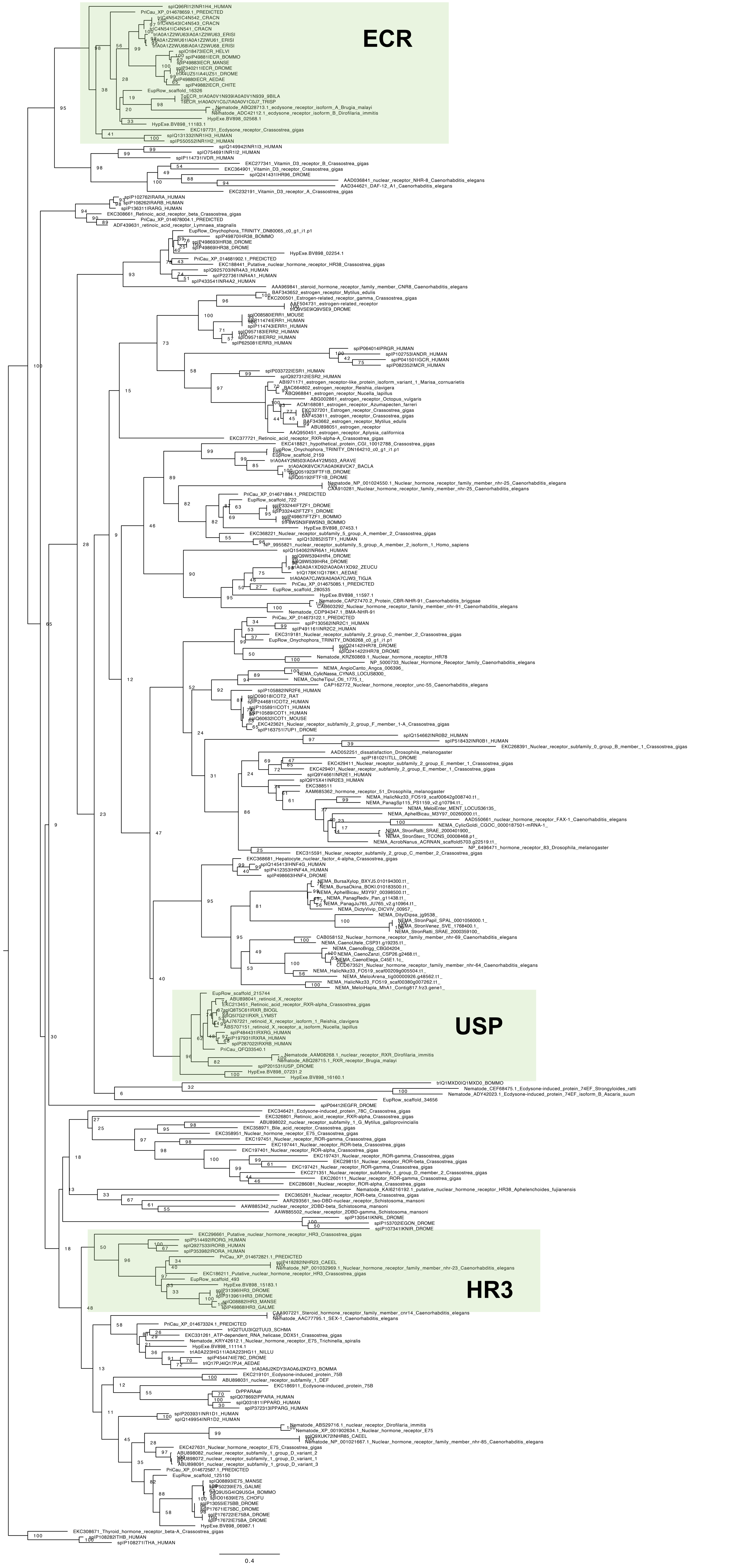

### tree_4th_screening.pdf

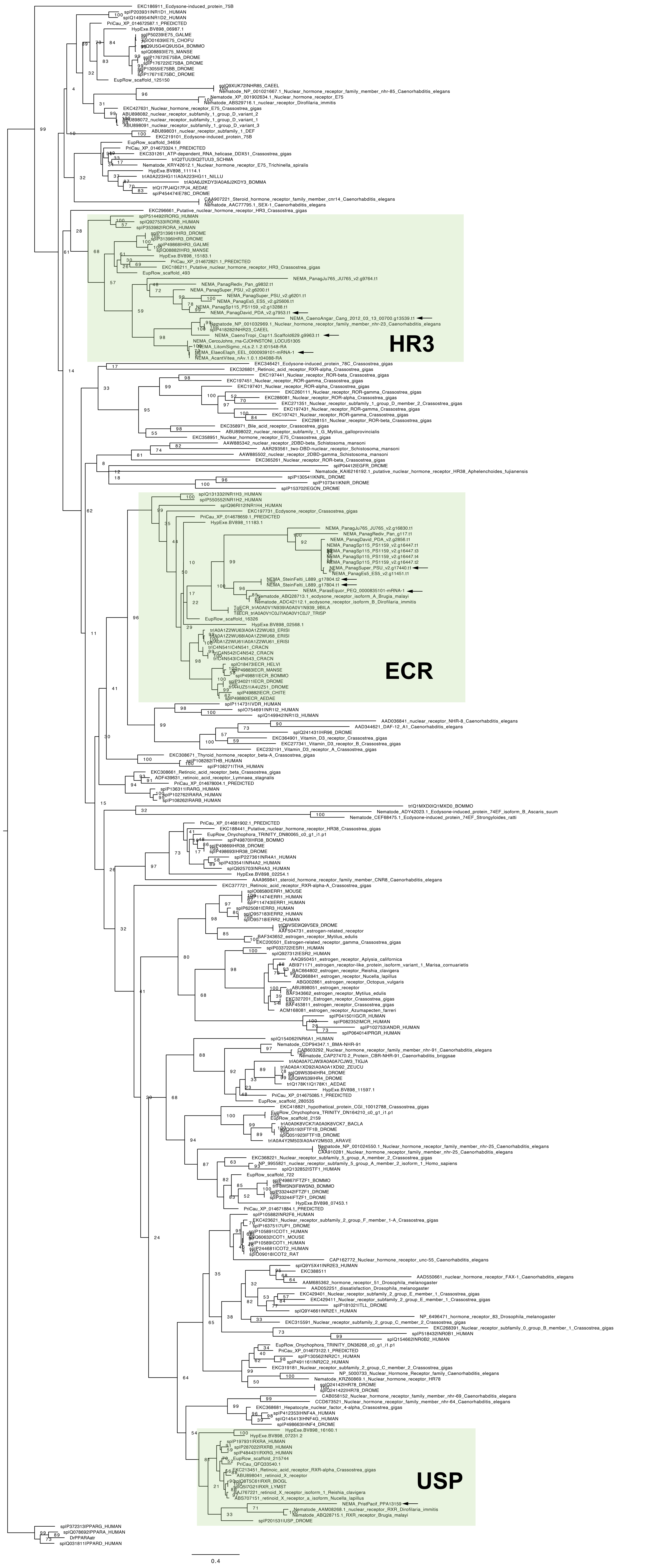

### tree_E75_screening.pdf

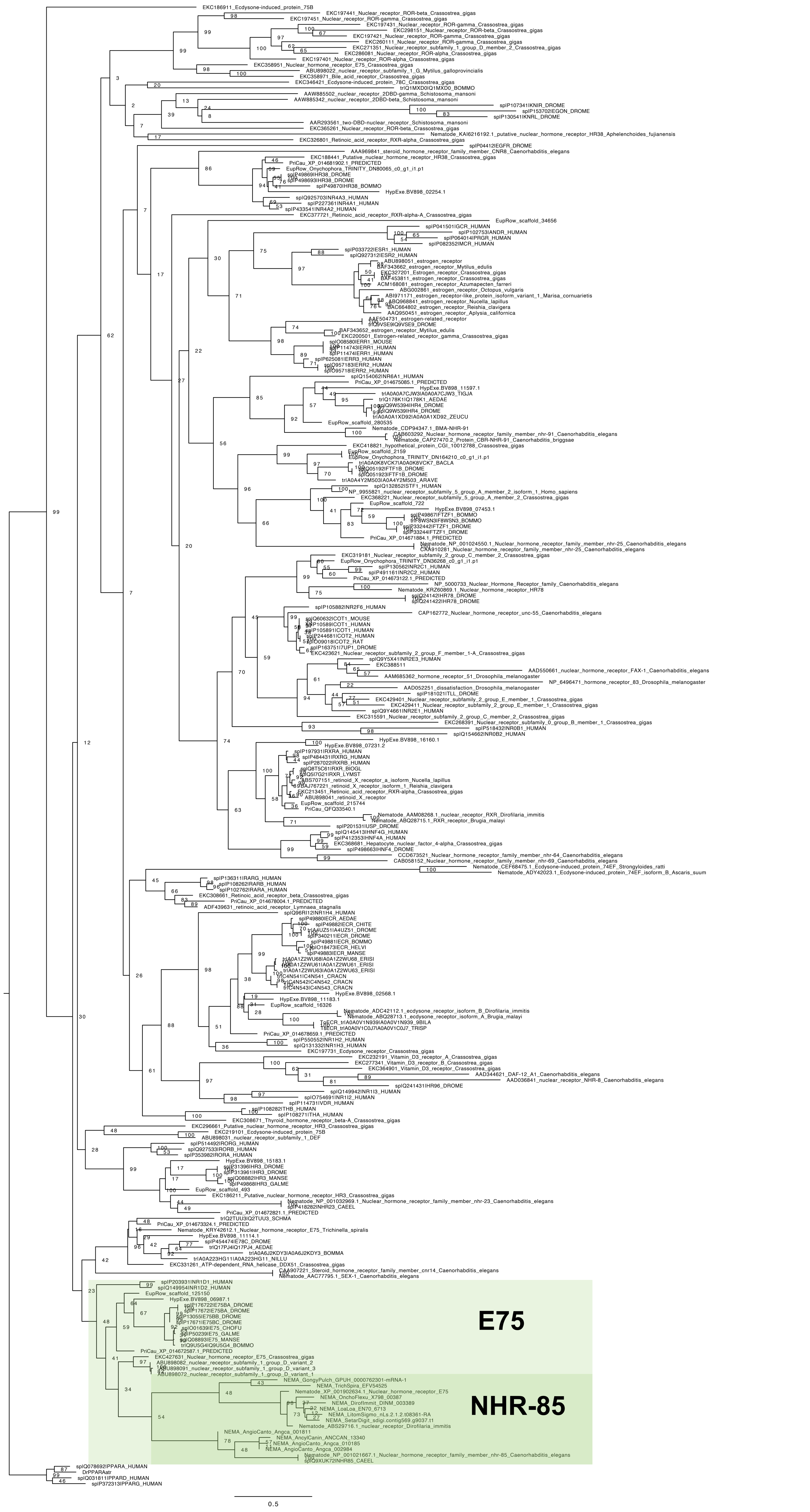

### tree_E75_screening_1_others.pdf

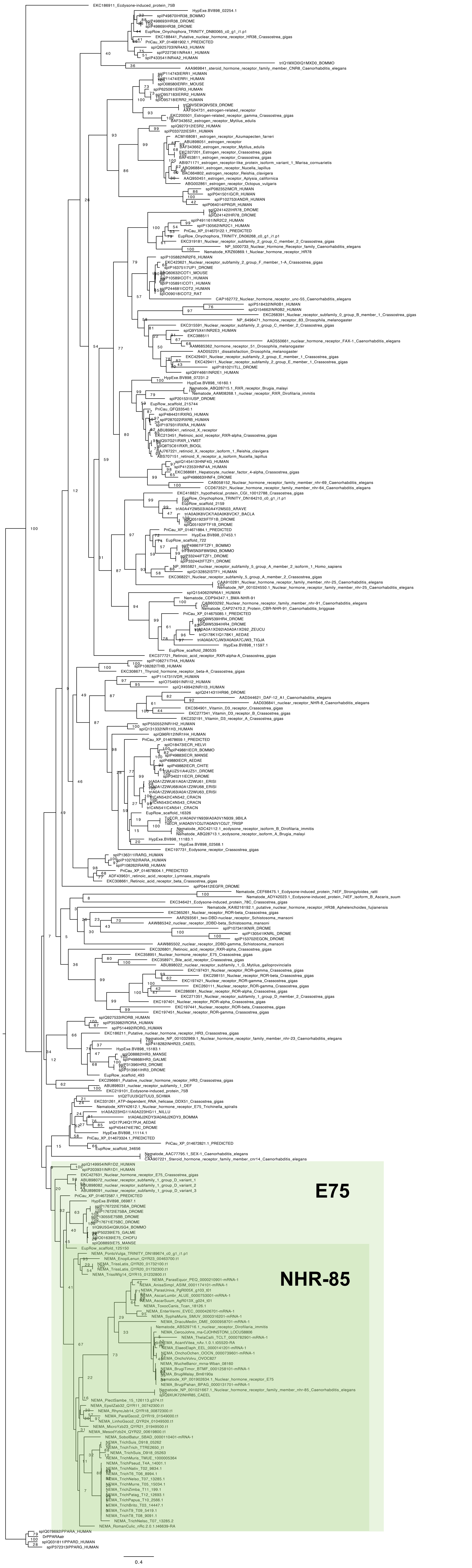

### tree_E75_screening_1_Rhabditina.pdf

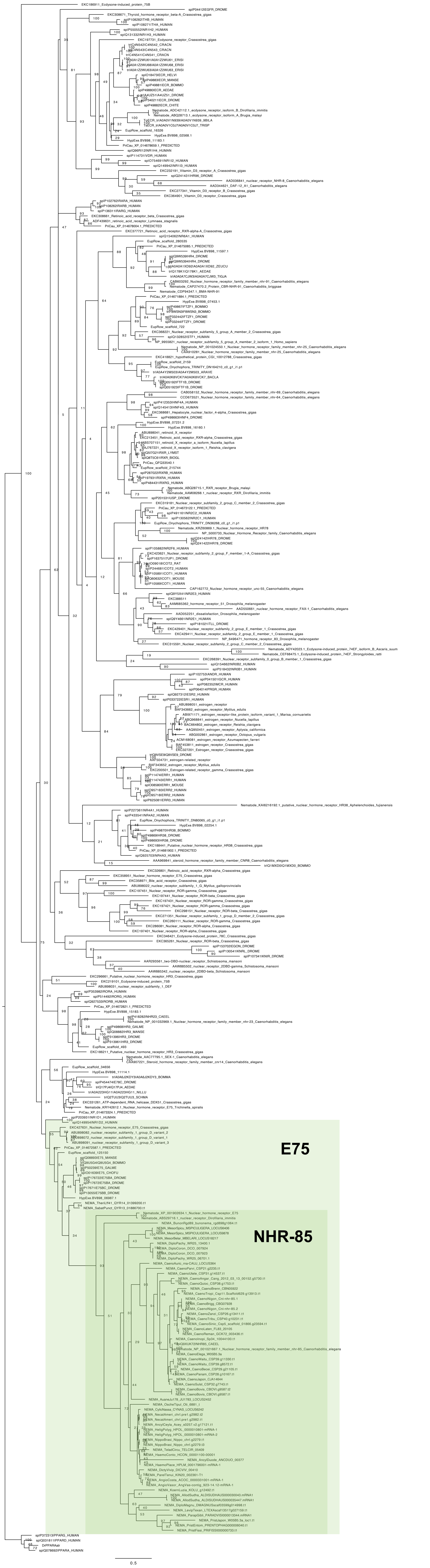

### tree_E75_screening_1_Tylenchina.pdf

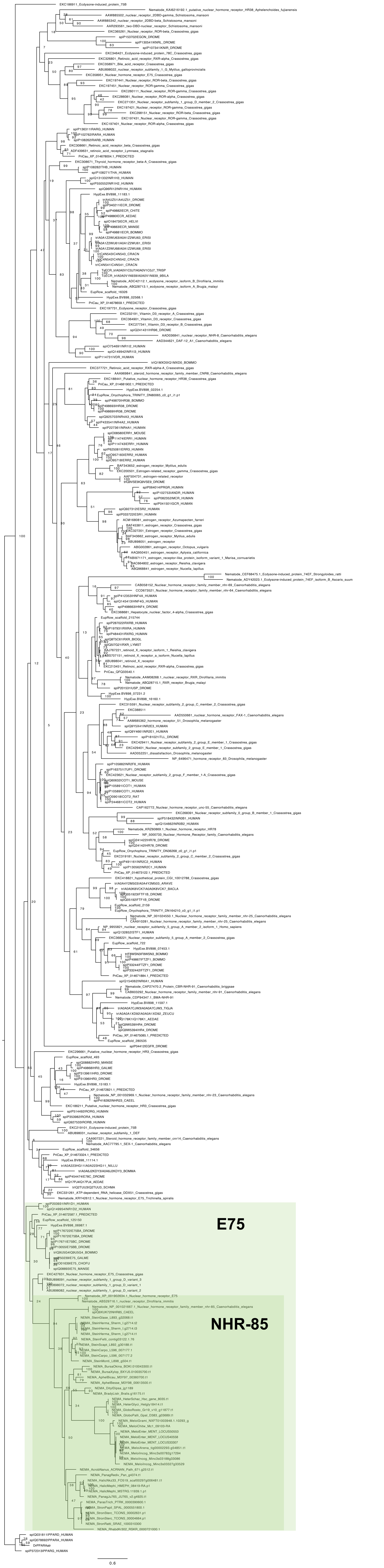

### tree_ECR_screening_1.pdf

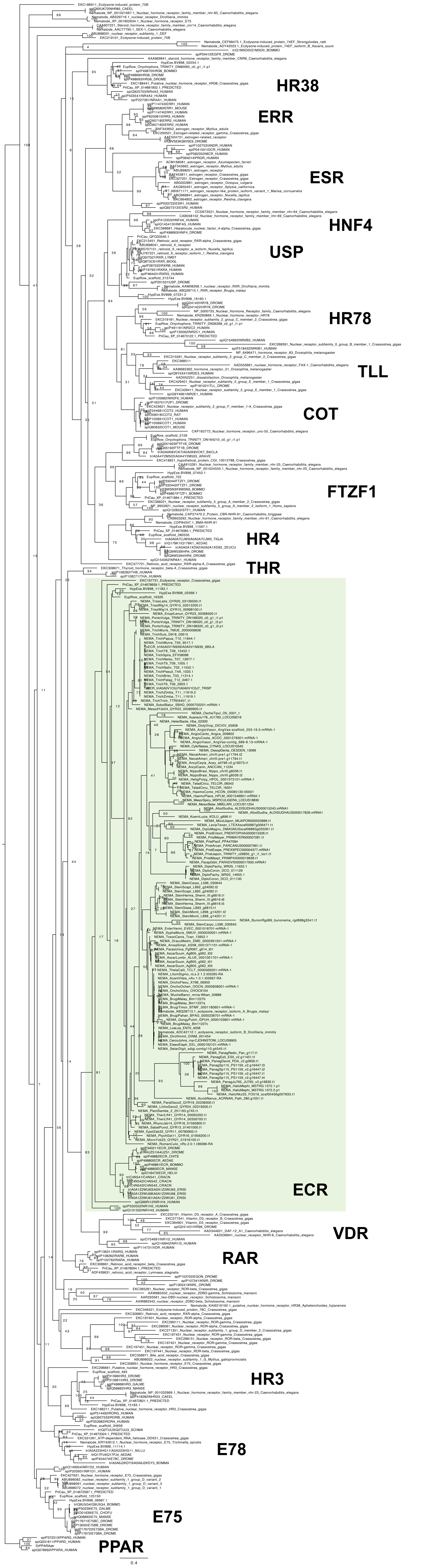

### tree_EcR_usp_Hr3.pdf

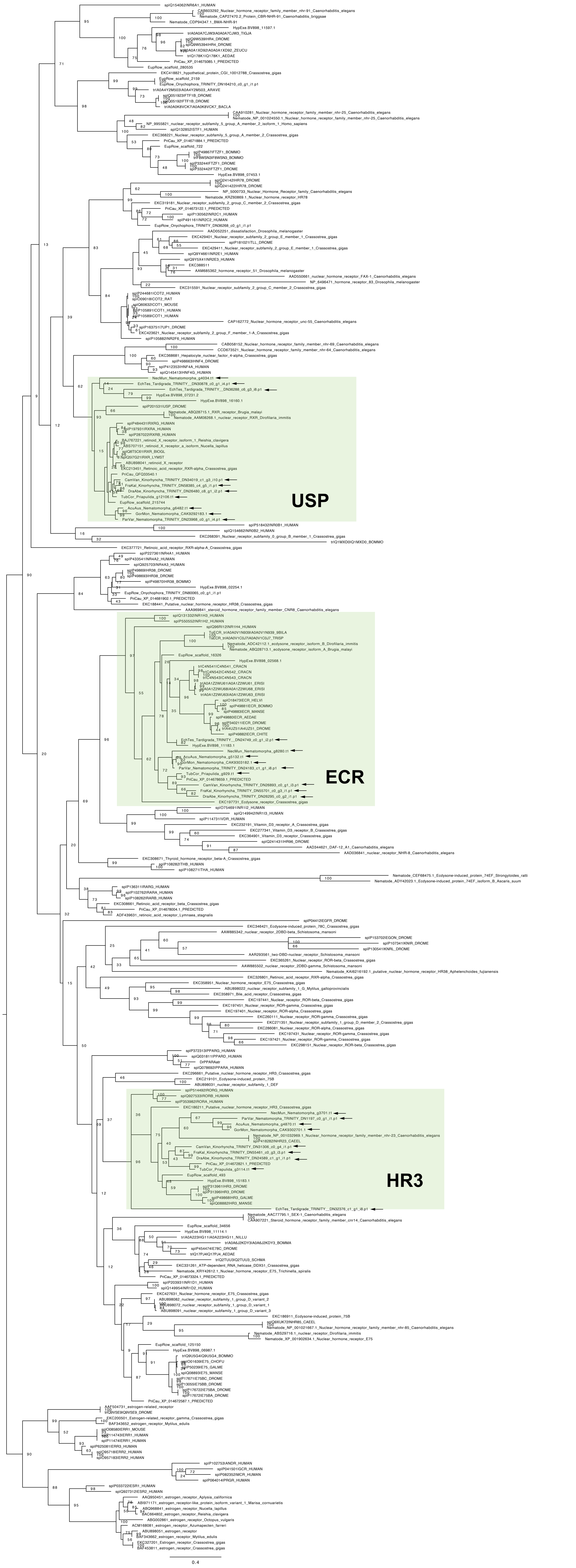

### tree_ECR_USP_HR3_screening.pdf

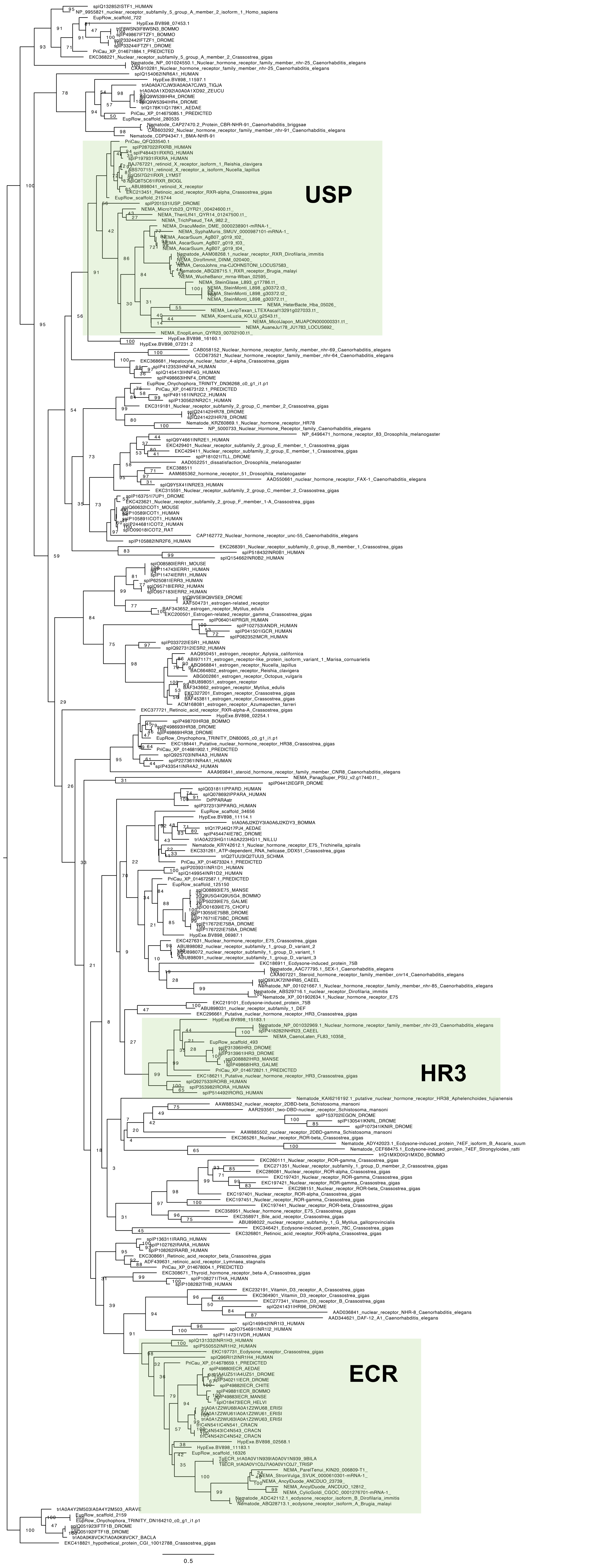

### tree_ftz-f1_screening.pdf

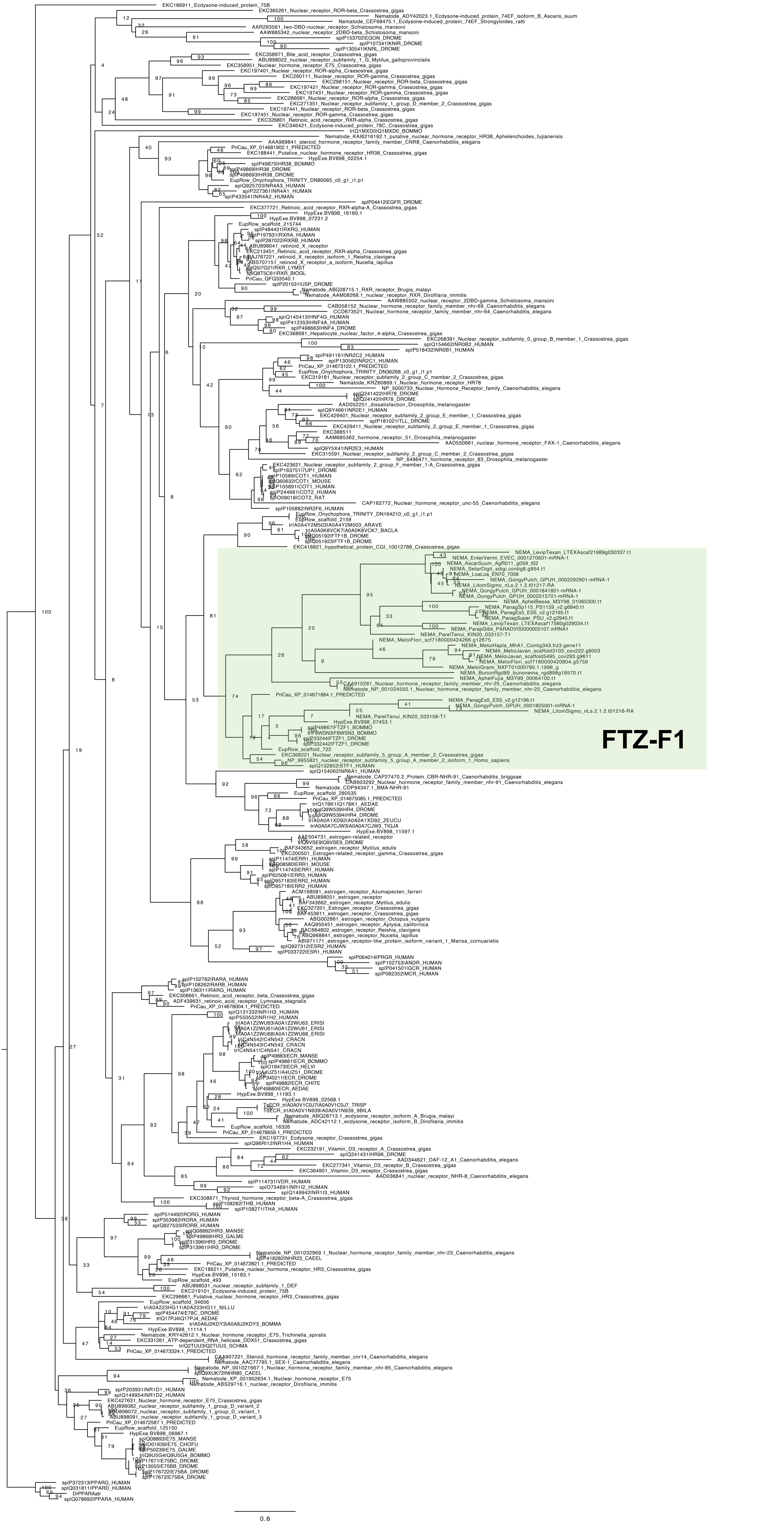

### tree_ftz-f1_screening_1_others.pdf

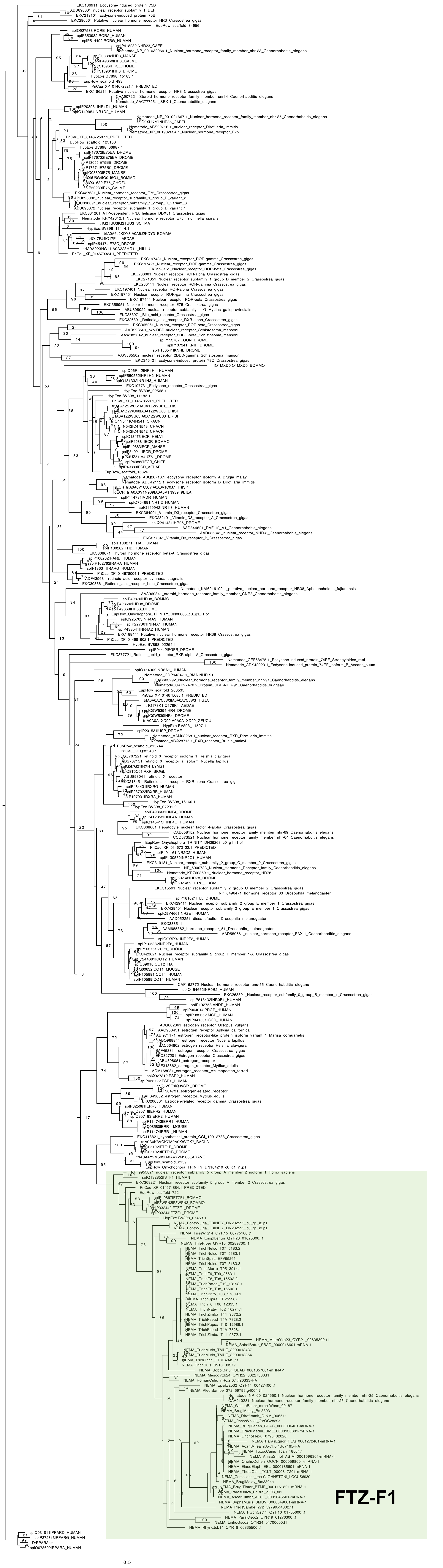

### tree_ftz-f1_screening_1_Rhabditina.pdf

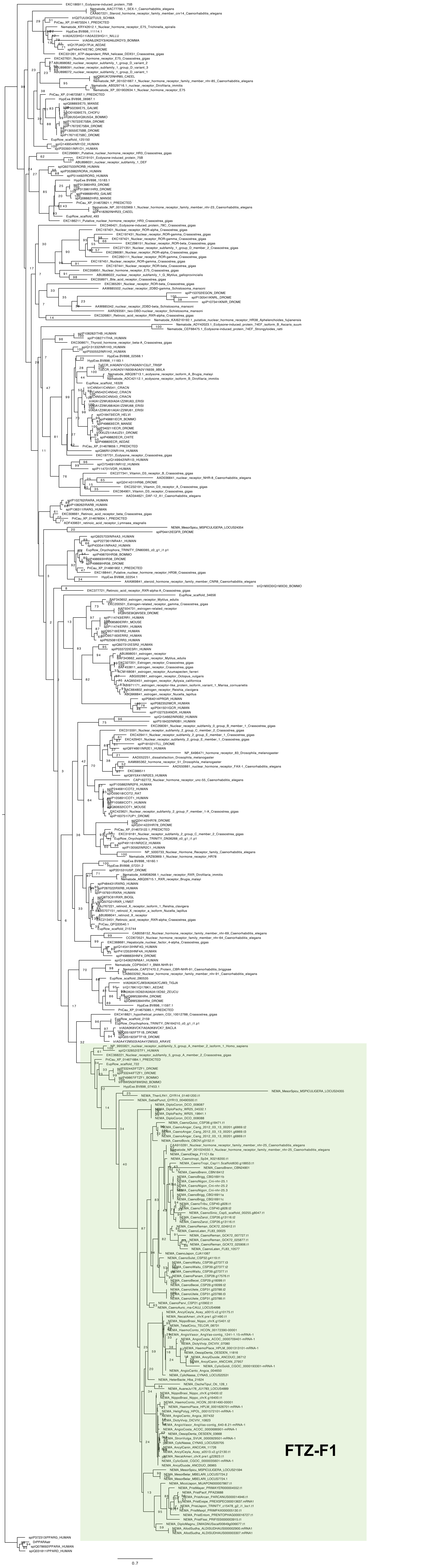

### tree_ftz-f1_screening_1_Rhabditina_Caenorhabditis.pdf

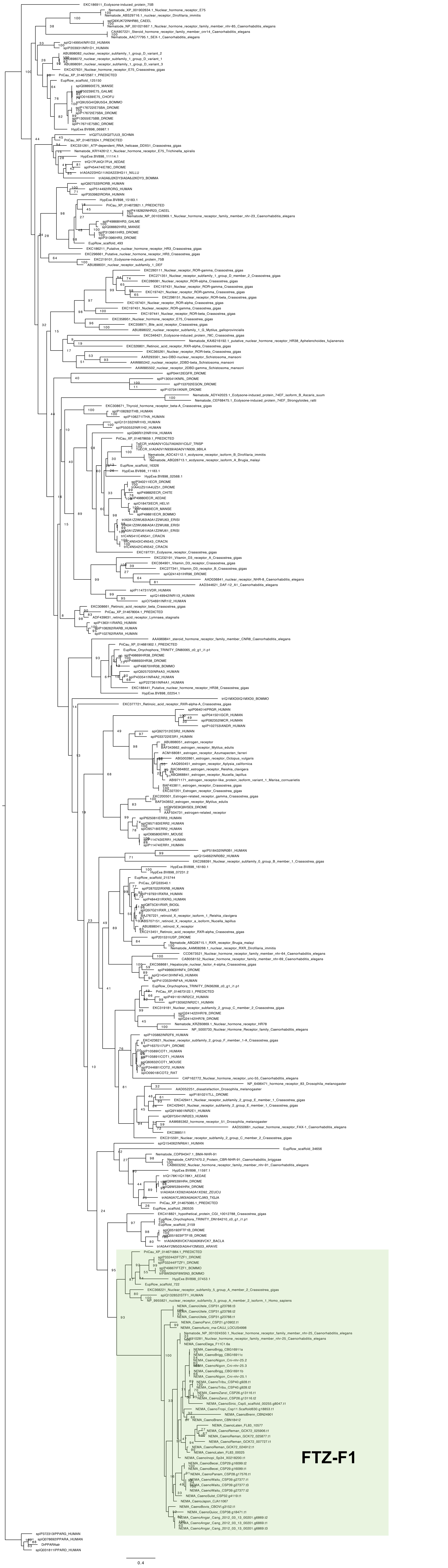

### tree_ftz-f1_screening_1_Rhabditina_nonCaenorhabditis.pdf

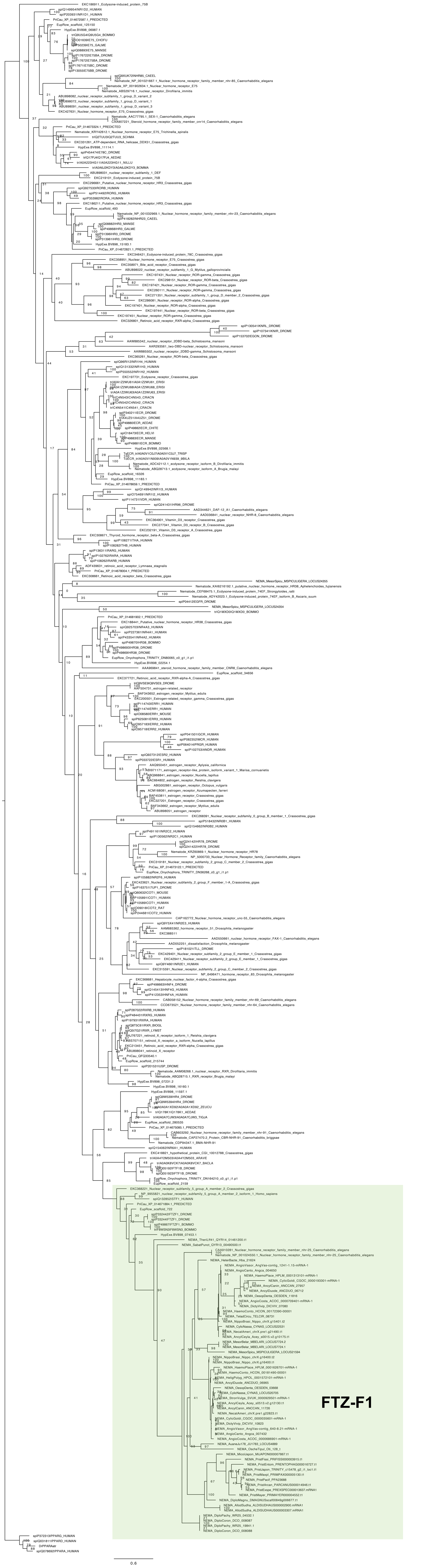

### tree_ftz-f1_screening_1_Rhabditina_Tylenchina.pdf

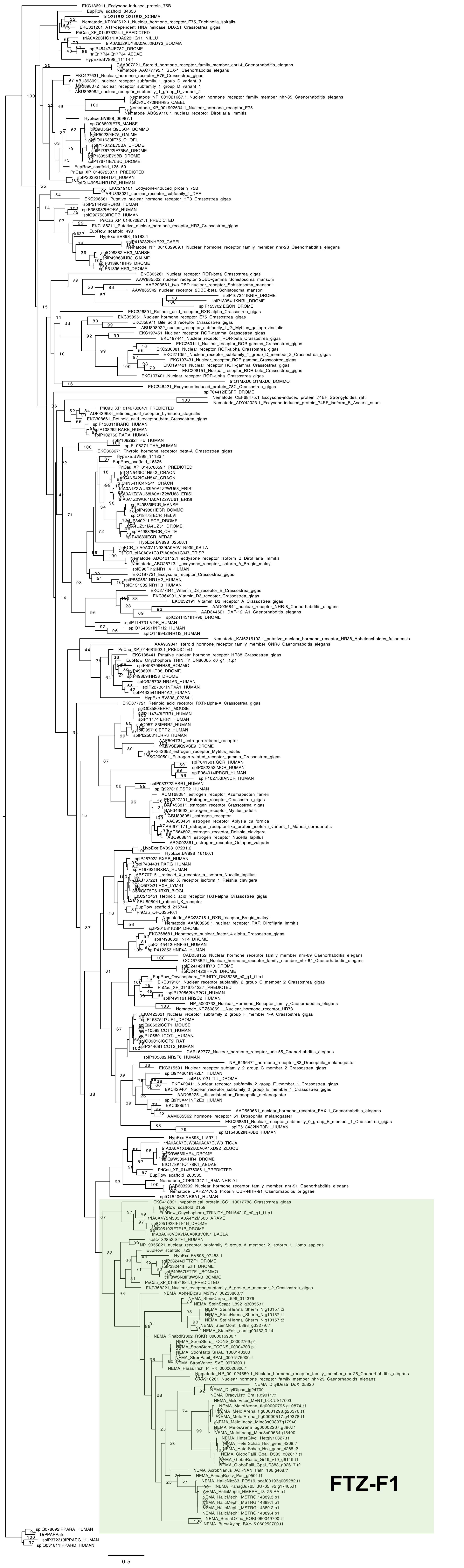

### tree_HR3_screening_1_all.pdf

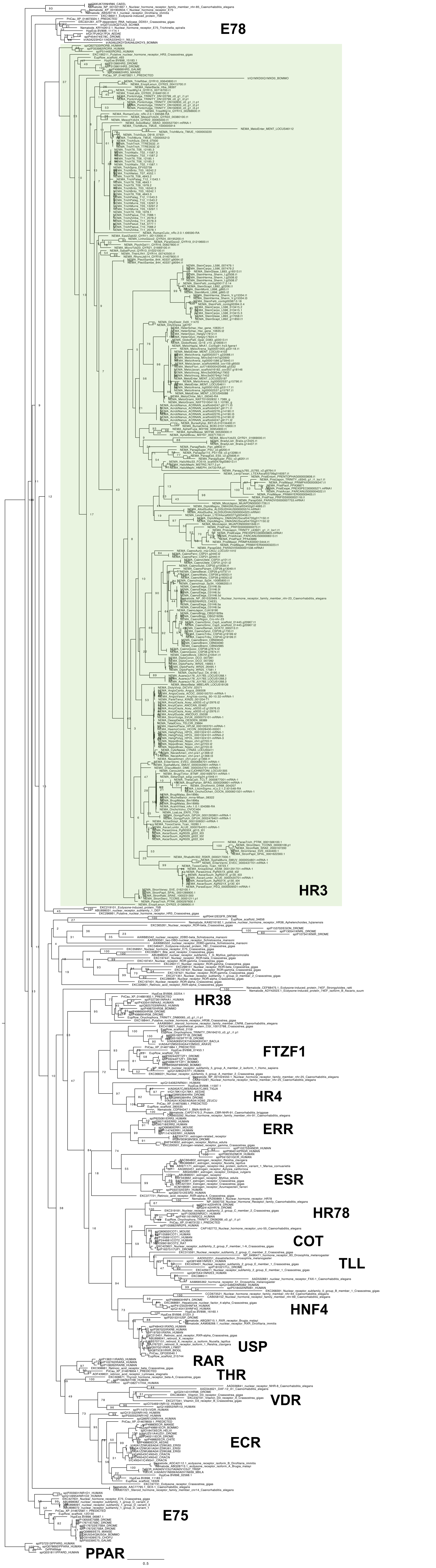

### tree_HR3_screening_1_others.pdf

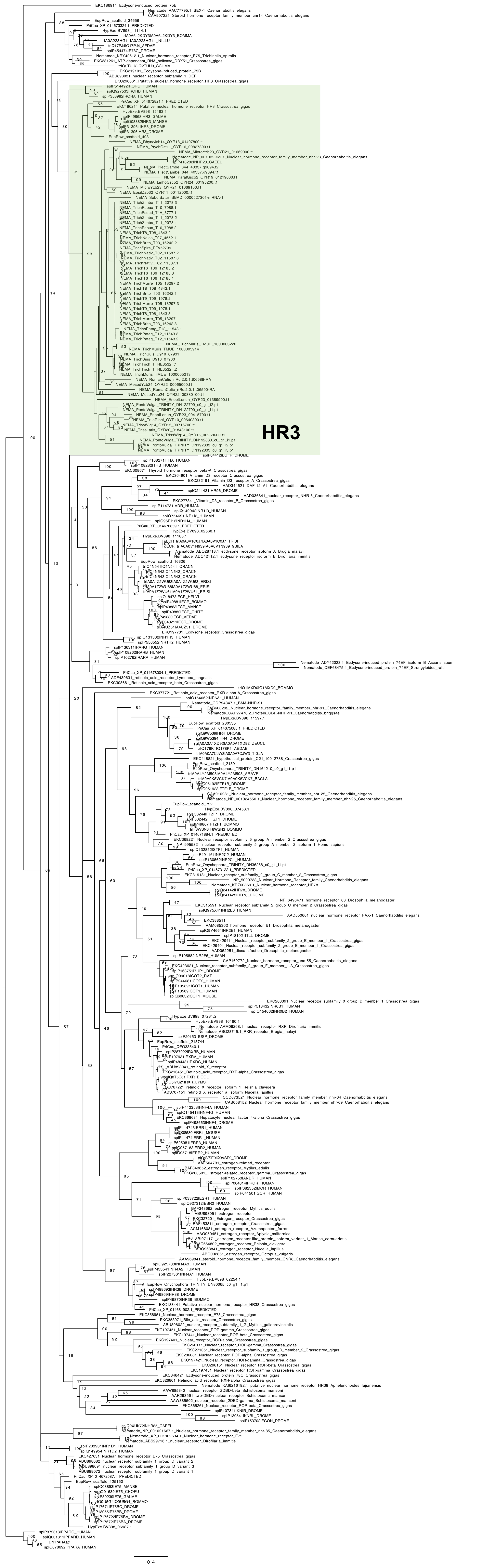

### tree_HR3_screening_1_Rhabditina.pdf

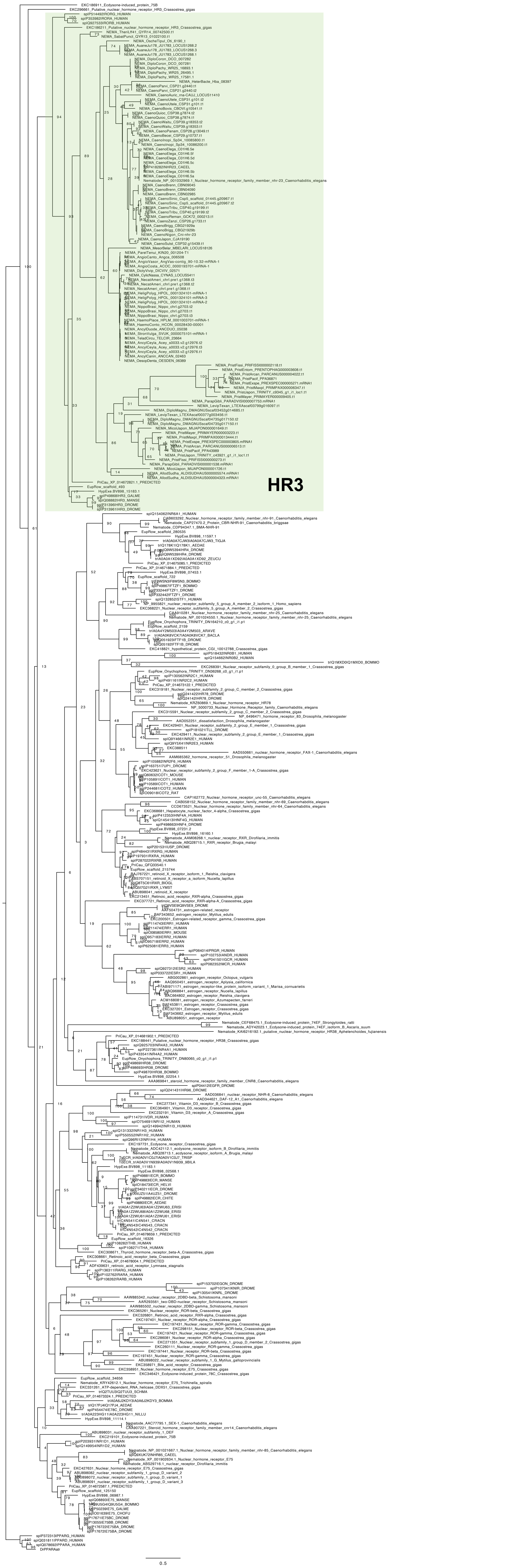

### tree_HR3_screening_1_Spirurira.pdf

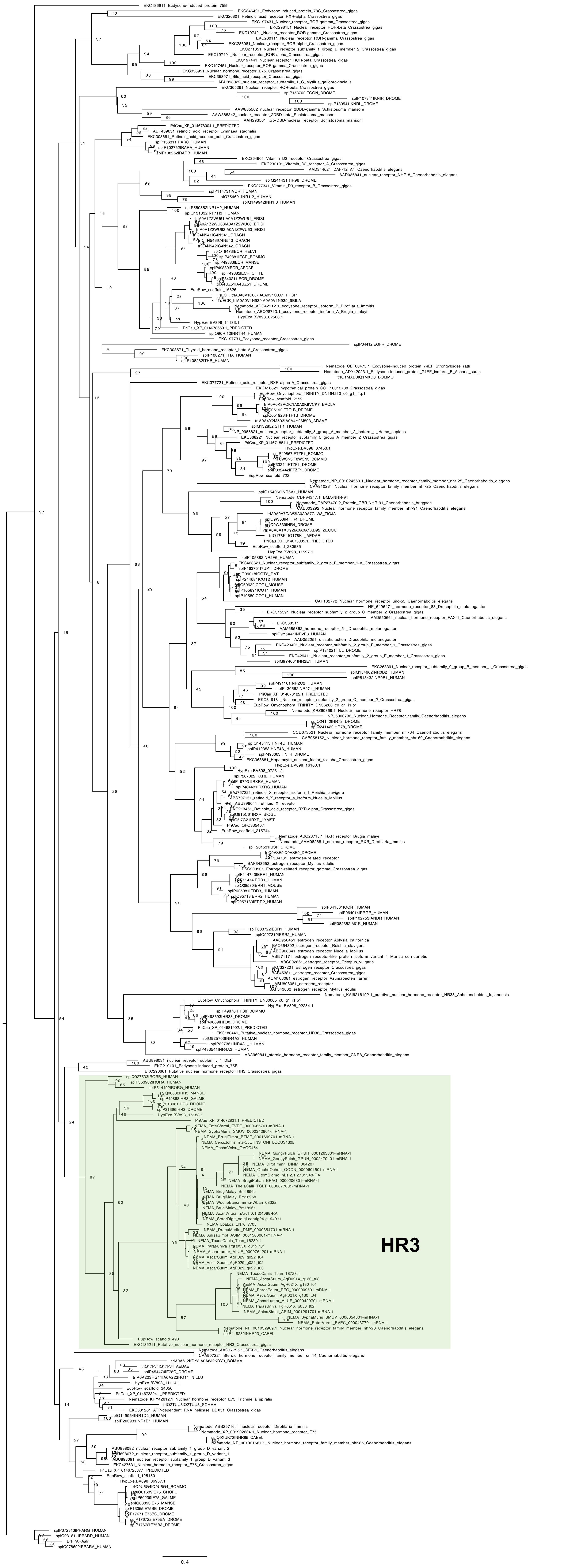

### tree_HR3_screening_1_Tylenchina.pdf

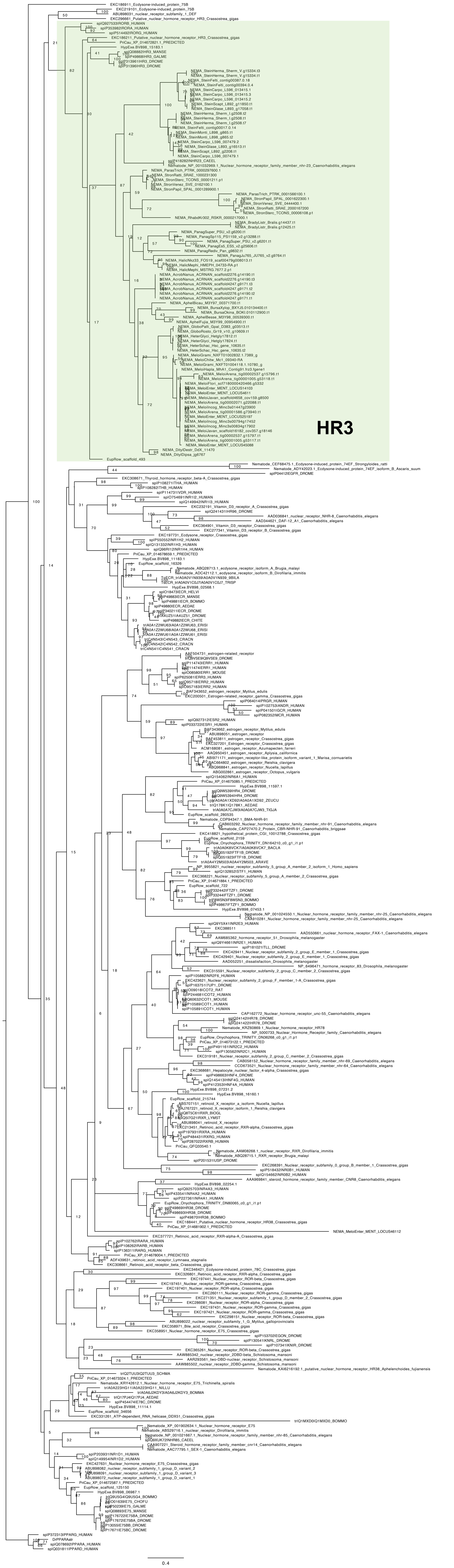

### tree_kin-20_screening_1.pdf

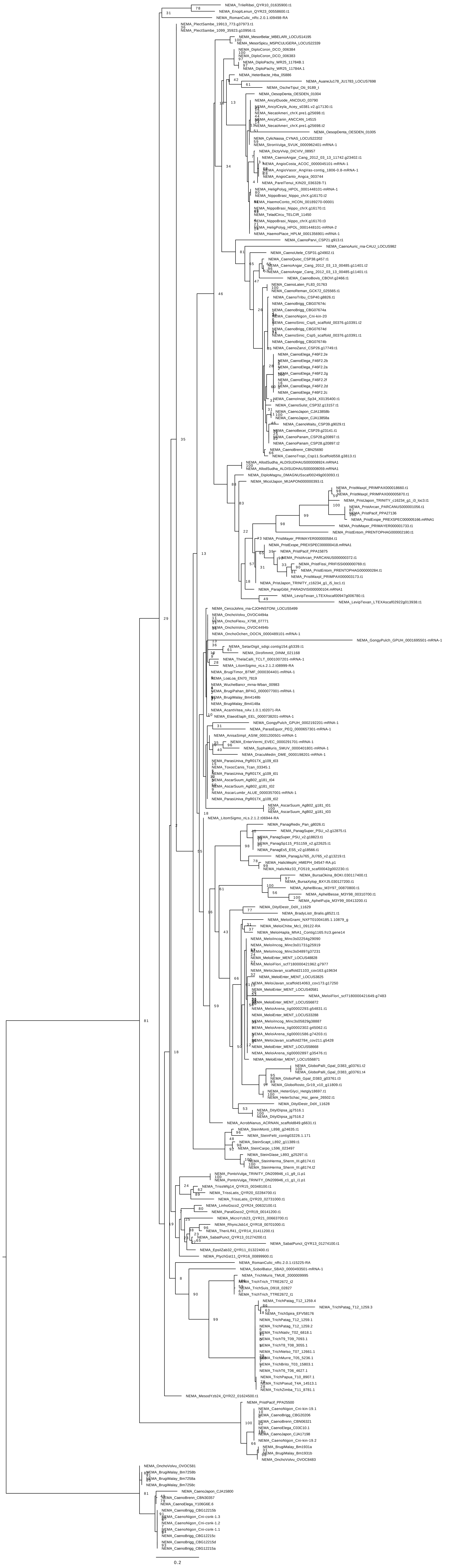

### tree_lin-42_screening.pdf

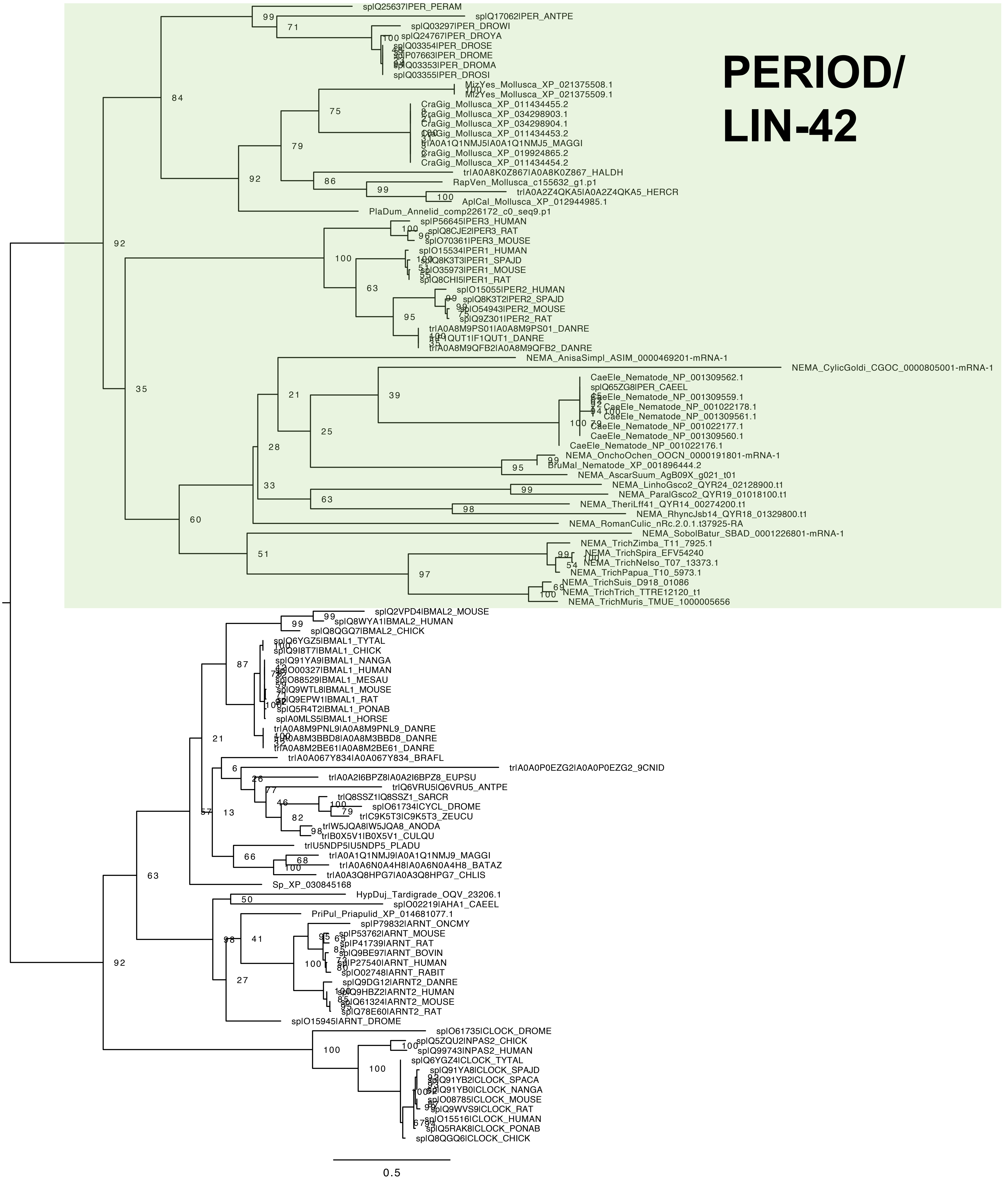

### tree_lin-42_screening_1.pdf

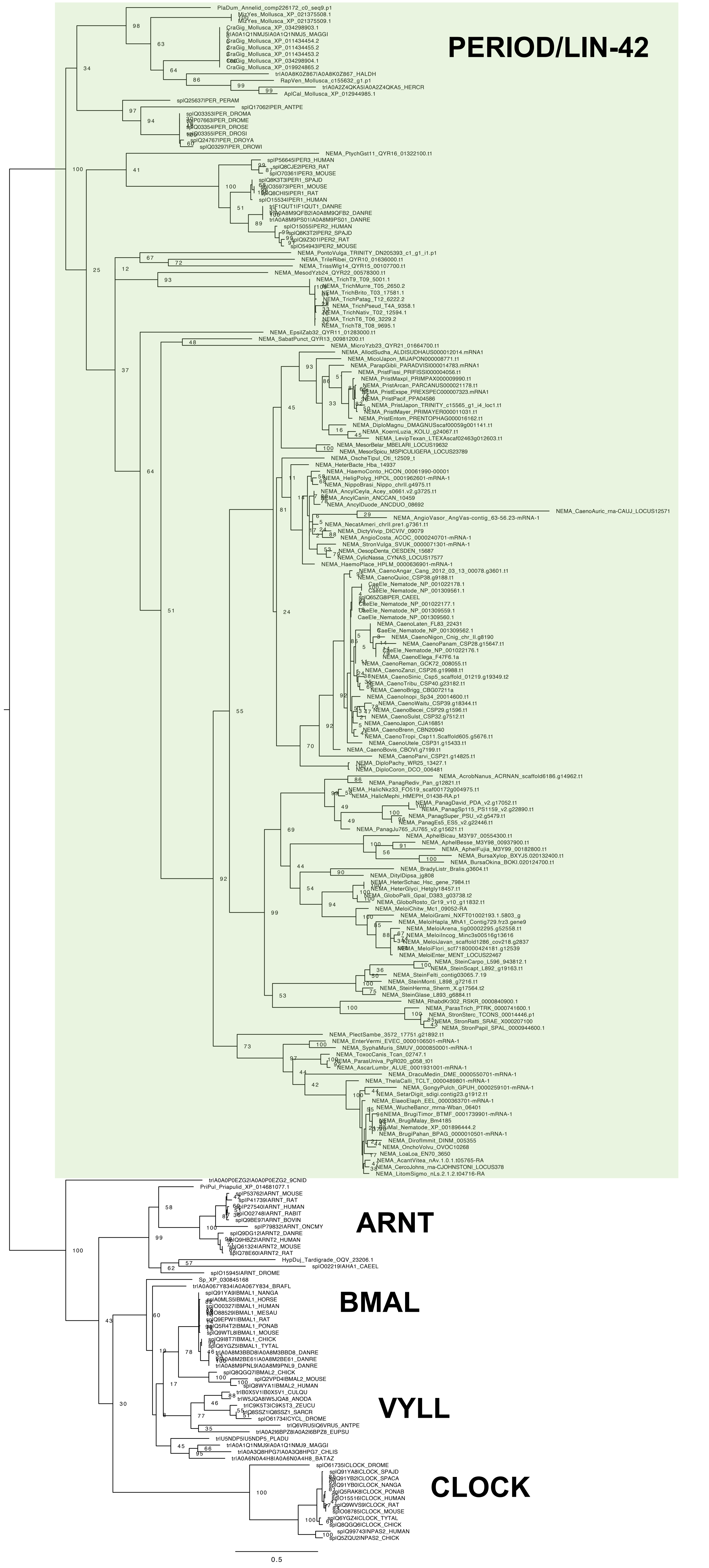

### tree_USP_screening_1.pdf

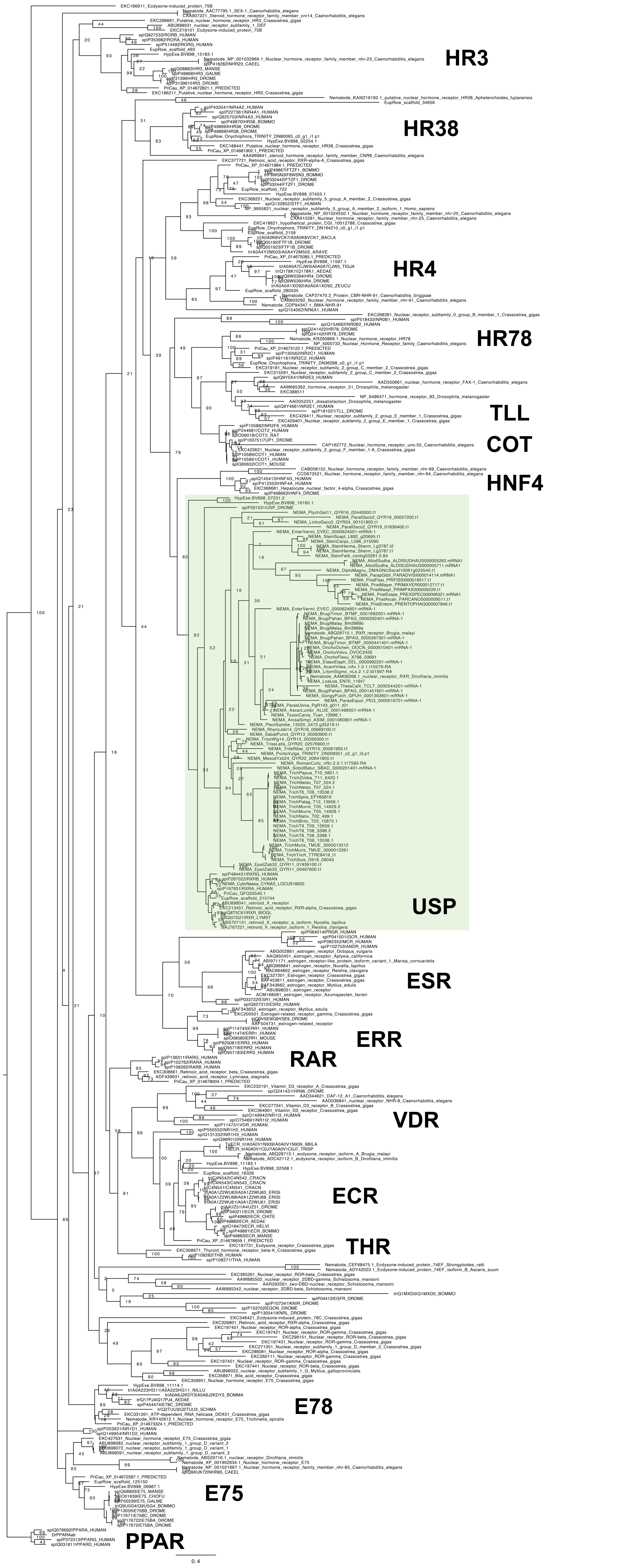

### usp_root_S1_bp1000_240613.png

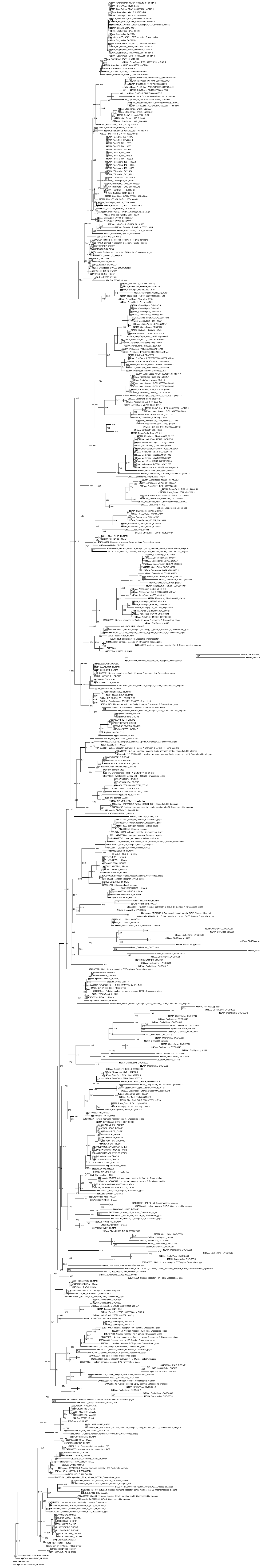
